# High glucose confers senescence resistance via GLUT1 epigenetic rewiring to blunt immunotherapy responses in esophageal squamous cell carcinoma

**DOI:** 10.64898/2026.07.15.738372

**Authors:** Jin-Xiu Dong, Jianian Zhou, Jia-Jie Hao, Shuai Kong, Chuntong Yin, Dan-Dan Wei, Fei Wang, Junboya Ma, Jinman Fang, Yuan-Wei Zhang, Huaguang Pan, Wen-Qiang Wei, Mingrong Wang, Kai Ma, Yuan Jiang, Yan-Yi Jiang

## Abstract

Therapeutic resistance and undefined predictive biomarkers severely hinder the clinical popularization of immunotherapy in esophageal squamous cell carcinoma (ESCC). Herein, we identify the glucose transporter 1 (GLUT1) as a critical determinant of immunotherapy resistance. Elevated expression of GLUT1 correlates with poor immunotherapy response and unfavorable prognosis in ESCC patients. GLUT1 deletion or inhibition enhances CD8⁺ T cell infiltration and cytotoxicity, and sensitizes ESCC tumors to anti-PD-1 (α-PD1) therapy. Importantly, dietary glucose restriction exhibits equivalent antitumor efficacy to GLUT1 inhibition when combined with α-PD1. Mechanistically, GLUT1 establishes a positive feedback loop with HAT1 and FOXM1, which epigenetically remodels chromatin accessibility to suppress tumor cell senescence, thereby impeding CD8⁺ T cell-mediated antitumor immunity. Our findings highlight GLUT1 as a predictive biomarker of immunotherapy resistance and suggest dietary glucose restriction as a viable strategy to potentiate immunotherapy efficacy in ESCC.

**Graphical abstract:** 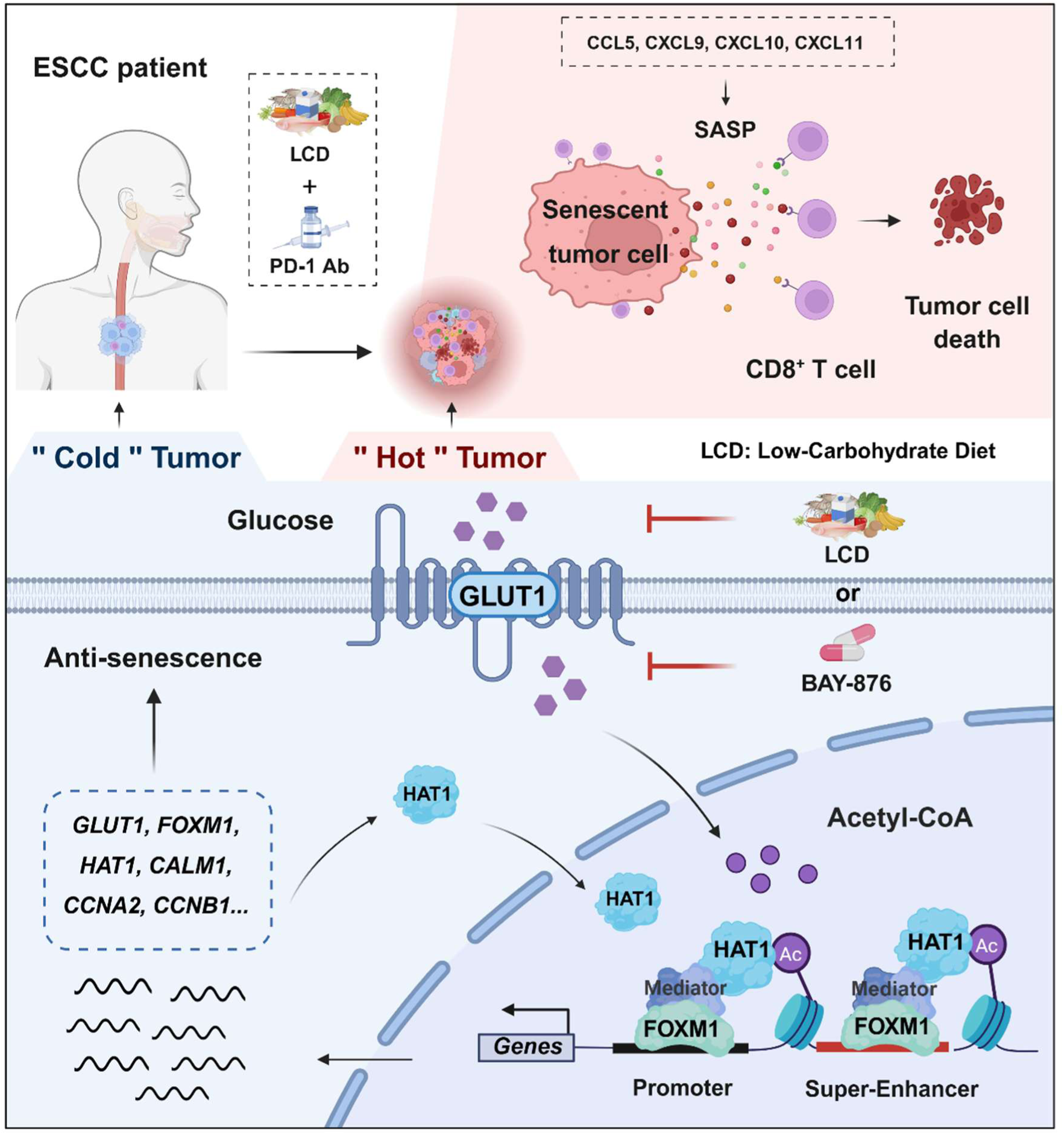

**SHORT SUMMARY:** Dong et al. identify glucose transporter 1 (GLUT1) as a predictor of poor response to immunotherapy in esophageal squamous cell carcinoma. GLUT1 promotes immune evasion by forming a positive feedback loop with the HAT1/FOXM1 anti-senescence axis, thereby epigenetically remodeling chromatin accessibility. GLUT1 inhibition or dietary glucose restriction restores CD8⁺ T-cell-mediated antitumor immunity and improves the efficacy of anti-PD-1 therapy in preclinical models.

## Main

Esophageal squamous cell carcinoma (ESCC) remains a major cause of cancer mortality in East and Central Asia, especially in China ^1^. Despite advances in multimodality management, including surgery, radiotherapy, and chemotherapy, clinical outcomes remain poor due to lack of specific therapeutic targets and the heterogeneity of the ESCC tumor microenvironment (TME) ^1,2^. In recent years, immunotherapy has brought substantial advancements in the treatment of various solid tumors ^3–5^. Notably, PD-1/PD-L1 blockade combined with chemotherapy has become the standard first-line and even neoadjuvant regimen for advanced ESCC ^6–8^. However, clinical responses remain limited to a subset of patients ^9^. Therefore, it is imperative to elucidate the molecular mechanisms underlying immune responsiveness and to develop predictive biomarkers for identifying eligible beneficiaries and optimizing therapeutic strategies.

Cellular senescence is a state of stable proliferative arrest in which viable cells remain metabolically active ^10^. In cancer, senescent cells have been recognized as a “hallmark of cancer” owing to their tumor-promoting effect ^11^. They can remodel the immunosuppressive microenvironment by expressing PD-L1 and CD80 ^12,13^ or secreting extracellular vesicles that induce T-cell senescence ^14^. However, the immunogenic potential of senescent cells enables them to elicit antitumor immunity, thereby potentially complementing antitumor immunotherapies ^15–18^. Such dual beneficial and detrimental effects on tissue biology appear highly dependent on tumor type, induction mode, and the TME ^19^. In ESCC, a recent study has identified senescent EGR1^+^ B cells as a key contributor to immunosuppression and neoadjuvant immunotherapy resistance ^20^. However, the crosstalk and molecular mechanisms connecting tumor cell senescence to tumor immunity remain largely underappreciated. GLUT1 (glucose transporter 1), encoded by *SLC2A1*, is a high-affinity glucose transporter ^21^. As a core regulator of tumor metabolic reprogramming, GLUT1 plays an indispensable role in the development and immune evasion of tumors, as well as therapy resistance. Studies in glioblastoma have demonstrated that GLUT1 not only promotes glycolysis and malignant progression through its S-palmitoylation ^22^, but also facilitates immune evasion by enhancing immunosuppressive activity of monocyte-derived macrophages via glucose-driven histone lactylation, and upregulating PD-L1 expression through HK2-mediated NF-κB activation ^23,24^. In melanoma, enhanced glycolysis in tumor cells is one of the key characteristics leading to immune resistance to adoptive T cell therapy ^25^. Although several studies have reported the high expression of GLUT1 in ESCC ^26,27^, the mechanistic role of GLUT1, particularly in regulating tumor cell senescence-mediated immune responses, has not been fully characterized.

In this study, we demonstrate that GLUT1, a key regulator of glucose metabolism, promotes tumor immune evasion by resisting tumor cell senescence in ESCC. Clinically, high GLUT1 expression independently predicts poor prognosis and confers immunochemotherapy resistance in ESCC patients. Mechanistically, GLUT1 forms a positive feedback loop with HAT1 and FOXM1, thereby cooperatively driving epigenetic remodeling of chromatin accessibility to suppress tumor cell senescence and senescence-associated secretory phenotype (SASP) chemokine production, consequently inhibiting CD8⁺ T cell function and ultimately impairing antitumor adaptive immunity. Our findings expand the functional landscape of GLUT1 beyond canonical glucose transport, uncover a previously unrecognized crosstalk between glucose metabolism and senescence-mediated antitumor immune responses, and suggest that glucose restriction represents a tractable strategy to enhance immunotherapy efficacy in ESCC.

## Results

### Elevated expression of GLUT1 in tumor tissues correlates with poor response to immunotherapy in ESCC patients

To systematically identify potential predictive and functional biomarkers associated with immunotherapy resistance in ESCC, we first analyzed publicly available scRNA-seq data of 42 pretreatment patient samples from three independent ESCC cohorts (Discovery cohort: HRA003312, HRA004396, and HRA004740) ^28–30^, in which patients received neoadjuvant anti-PD-1 (α-PD1) therapy plus chemotherapy (Figure 1A). We identified 319 genes that could effectively distinguish non-complete response (NCR) from pathological complete response (CR). Of these, 116 candidate genes exhibited elevated expression in tumor epithelium relative to normal counterparts. Notably, correlation analysis revealed that GLUT1 showed the strongest negative correlation with CD8⁺ T cell infiltration (Figures 1B and S1A).

**Figure 1.**
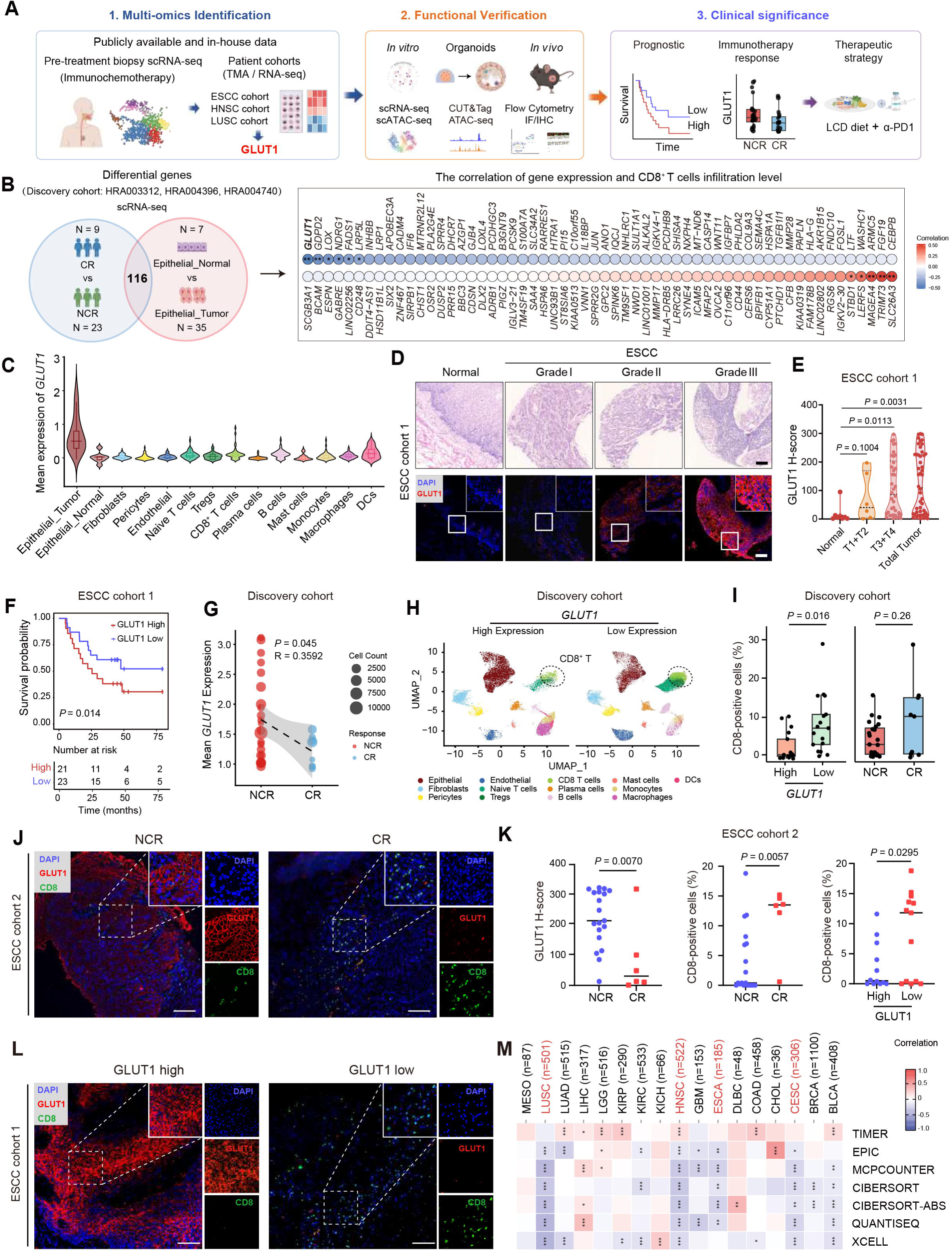
Elevated GLUT1 in tumor tissues correlates with poor response to immunochemotherapy in ESCC patients. (A) Study overview. (B) Left: Venn diagram showing overlapping differential genes (116 candidates). Right: ranking correlations of 116 differential genes with CD8⁺ T cell infiltration, with GLUT1 showing the most significant negative correlation. scRNA-seq and matched clinical data from three independent ESCC datasets (Discovery cohort: HRA003312, HRA004396, and HRA004740). (C) scRNA-seq analysis showing the distribution of *GLUT1* expression across different cell populations in ESCC. (D) Representative hematoxylin-eosin (H&E) staining (top) and immunofluorescence (IF) staining (bottom) of ESCC tissues with different pathological grades. Insets indicate magnified regions. Scale bar, 100 μm. (E) Quantification of GLUT1 protein expression (H-score) in (D). Statistical significance was determined as indicated. (F) Kaplan-Meier analysis of overall survival in the ESCC cohort 1, with patients stratified into high- and low-GLUT1 expression groups based on the median GLUT1 expression level (top 50% vs. bottom 50%). The log-rank test *P* value is shown, with numbers at risk displayed below. (G) Comparison of mean GLUT1 expression between non-complete response (NCR) and complete response (CR). Each dot represents a sample; point size reflects cell count. Linear regression statistics (*P* and R) are shown. Data from Discovery cohort.| (H) UMAP visualization of scRNA-seq data annotated by cell types, separately presented in *GLUT1* high- and low-expression groups. (I) Quantification of CD8⁺ T cell proportions according to *GLUT1* expression (high vs. low) and treatment response (NCR vs. CR) to immunochemotherapy. *P* values are indicated. (J) Representative IF images from ESCC patients with NCR and CR. Insets highlight magnified regions. (K) Quantification of GLUT1 H-score and CD8⁺ T cell percentages in NCR and CR patient samples, and CD8⁺ T cell percentages stratified by GLUT1 high vs. low expression. (L) Representative IF images of tumors with GLUT1 high and low expression, stained for DAPI, GLUT1, and CD8a. Insets show magnified regions. Scale bar, 100 μm. (M) Bulk RNA-seq analysis showing a negative correlation between *GLUT1* expression and CD8⁺ T cell infiltration. Data from TCGA. Data of (B, C, G, H, I) are from Discovery cohort: HRA003312, HRA004396, and HRA004740. Data of (D, E, F, L) from ESCC cohort 1. Data of (J, K) from ESCC cohort 2. Discovery cohort: 9 CR, 23 NCR, and 3 NA (response data unavailable); ESCC cohort 2: 6 CR, 19 NCR. Statistical analysis was performed using Student’s t-test (G, I, and K) and one-way ANOVA with Tukey’s multiple comparison test (E).

To further investigate the clinical relevance of GLUT1 in ESCC, we integrated data from 42 scRNA-seq datasets, protein expression data from 137 in-house FFPE tissue specimens, and 1,119 bulk RNA-seq samples. First, GLUT1 had remarkably high expression in epithelial cells, especially in malignant tumor cells when compared with other cell types such as immune cells, fibroblasts, or endothelial cells (Figure 1C). Second, among the 14 GLUT-family members, GLUT1 displayed the highest expression level and the most significant differential expression between tumor and paired normal tissues in patients with ESCC, as well as in other types of squamous cell carcinoma (Figures 1D, 1E, and S1B). Third, GLUT1 expression was gradually elevated from early to advanced ESCC and positively correlated with clinical stage (Figure 1D and 1E). Importantly, high GLUT1 expression was significantly correlated with poor ESCC prognosis, and served as an independent prognostic factor in ESCC patients (Figures 1F and S1C). In addition, *GLUT1* expression was consistently elevated in NCR patients compared with CR patients undergoing neoadjuvant immunochemotherapy across three independent datasets, with statistical significance confirmed by linear regression analysis (*P* = 0.045, R = 0.359) (Figure 1G). Moreover, this association was not observed for *CD274* (*PD-L1*) expression (Figure S1D), suggesting that GLUT1 may represent a superior predictive biomarker for stratifying neoadjuvant immunochemotherapy response in this clinical setting.

Consistent with these findings, tumors with low *GLUT1* expression and those from CR patients contained higher proportions of CD8⁺ T cells, revealed by scRNA-seq data (Figure 1H and 1I). As a validation, we analyzed GLUT1 expression at the protein level in another clinical ESCC cohort of 25 patients receiving α-PD1-based immunochemotherapy (ESCC cohort 2, samples from Hefei Cancer Hospital of Chinese Academy of Sciences and The First Affiliated Hospital of Anhui Medical University). GLUT1 expression was significantly higher in NCR than in CR, accompanied by a reduced infiltration of CD8⁺ T cells (Figure 1J and 1K). This inverse association was further validated by immunofluorescence (IF) staining using our in-house tissue microarrays derived from patients with ESCC and LUSC (Figures 1L, S1E, S1F and S2A-S2C). Moreover, independent bulk RNA-seq datasets also confirmed a significant negative correlation between *GLUT1* level and CD8⁺ T cell infiltration across squamous cell carcinomas (Figure 1M). Thus, these findings identify GLUT1 as a reliable predictive biomarker for immunotherapy resistance in ESCC, potentially by suppressing CD8⁺ T cell infiltration.

### GLUT1 inhibits CD8⁺ T cell infiltration and cytotoxic activity in ESCC tumors

To further explore why GLUT1 may serve as a predictive biomarker of therapeutic response, we first investigated its role in the tumor microenvironment (TME) by establishing syngeneic murine ESCC models in immunocompetent C57BL/6J mice. Notably, *Glut1* knockdown significantly suppressed tumor growth *in vivo* (Figures 2A, S2D, and S2E). To dissect the TME remodeling, we performed scRNA-seq on excised allograft tumors. *Glut1* knockdown markedly reshaped the tumor immune landscape, characterized by increased proportions of CD8⁺ T cells, CD4⁺ T cells, and NK/NKT cells, coupled with a reduction in macrophages and dendritic cells (Figure 2B). The elevated proportion of CD8⁺ T cells was confirmed by flow cytometry (Figure 2C) and further supported by IF staining, which visually demonstrated enhanced CD8⁺ T cell infiltration into *Glut1*-knockdown tumors (Figure 2D). Cell-cell communication analysis revealed that knockdown of *Glut1* led to the most prominent enhancement of crosstalk between tumor epithelial cells and CD8⁺ T cells (Figure 2E and 2F), suggesting that tumor-intrinsic GLUT1 upregulation may actively constrain CD8⁺ T cell recruitment and engagement within the TME.

**Figure 2.**
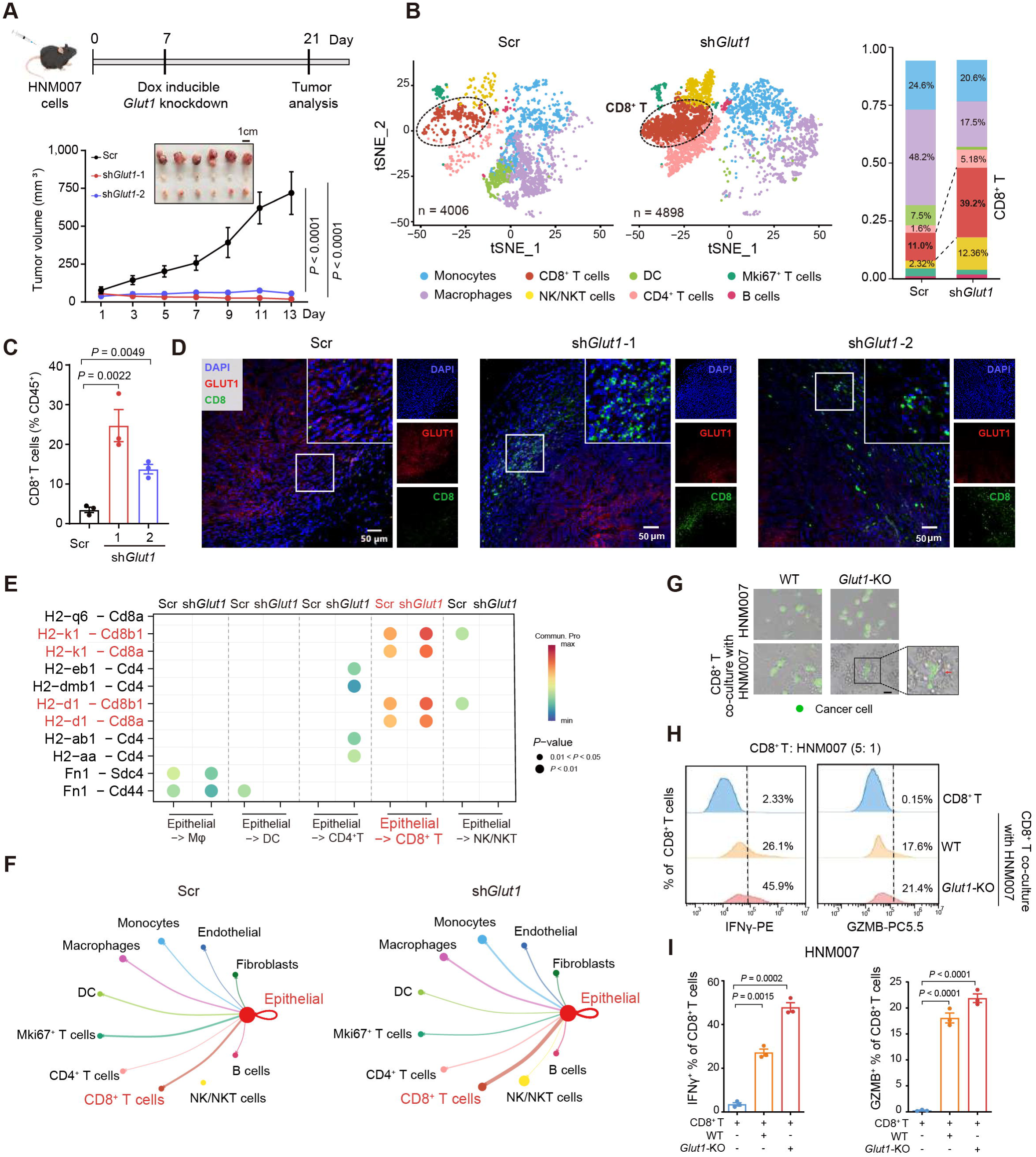
GLUT1 suppresses CD8⁺ T cell infiltration and cytotoxic function. (A) Experimental schematic of the subcutaneous tumor model in C57BL/6J mice. Mice were subcutaneously injected with 5 × 10⁵ HNM007 cells, followed by tumor growth monitoring. Representative tumor images and growth curves are shown. N = 6. *P* values indicate statistical significance between the indicated groups, as determined by one-way ANOVA followed by Tukey’s multiple comparisons test. (B) UMAP plot of cell types identified in mouse tumor tissues via scRNA-seq (left), and stacked plot showing the proportional changes of immune cell subsets after *Glut1* knockdown (sh*Glut1*) (right). (C and D) Flow cytometry (C) and IF staining (D) analyses showing increased CD8⁺ T cell infiltration after *Glut1* knockdown. (E) Dot plot showing predicted ligand-receptor interactions among epithelial, stromal, and immune cell populations in control and sh*Glut1* tumors. (F) Cell-cell communication networks centered on epithelial cells in Scramble control (Scr) and sh*Glut1* groups. Edges represent predicted signaling interactions between epithelial cells and surrounding immune or stromal populations, and line thickness reflects the intensity of intercellular crosstalk. (G) Co-culture of sh*Glut1* HNM007 tumor cells with CD8⁺ T cells, tumor cell viability was assessed by Calcein AM (green). (H and I) Flow cytometry (H) and statistical analysis of three independent assays (I) showing increased frequencies of IFNγ⁺ and GZMB⁺ CD8⁺ T cells following *Glut1* knockdown. Data are presented as mean ± SEM, and statistical comparisons were performed using one-way ANOVA with Tukey’s multiple comparison test (C and I).

To directly evaluate CD8⁺ T cell cytotoxicity, we performed *ex vivo* co-culture assays using SIINFEKL (OVA_257-264_)-pulsed ESCC cells (WT or *Glut1*-KO HNM007 cells) and OVA_257-264_-specific CD8⁺ T cells isolated from OT-I mice (Figure 2G). *Glut1*-KO ESCC cells exhibited markedly higher susceptibility to T cell-mediated cytotoxicity, resulting in a prominent reduction in viable tumor cells (Figures 2G, S2F, and S2G). Furthermore, CD8⁺ T cells co-cultured with the *Glut1*-KO cancer cells displayed a substantial increase in cytotoxic populations positive for GZMB and IFNγ (Figures 2H, 2I, and S2H). These results demonstrate that GLUT1 suppresses antitumor immunity by impairing CD8⁺ T cell infiltration and cytotoxic function in the ESCC TME.

### GLUT1 suppresses cellular senescence to sustain tumor progression in ESCC

To determine the tumor-intrinsic mechanisms by which GLUT1 impacts CD8⁺ T cell function, we first investigated the basis for the aberrant upregulation of GLUT1 in ESCC. Neither frequent mutation nor copy number amplification accounted for elevated GLUT1 expression in ESCC patients or across 1,119 squamous cell carcinoma samples (Figure S3A). Instead, we observed a pronounced enrichment of the active enhancer mark H3K27ac around the *GLUT1* locus in both ESCC tissues and cell lines relative to their normal counterparts, accompanied by high-density occupancy of RNAPII, H3K4me3, and H3K4me1 (Figure S3B). GLUT1 was then identified as a super-enhancer (SE)-associated gene, along with well-characterized ESCC oncogenes including *KLF5* and *SOX2* ^31,32^ (Figure S3C). H3K27ac signal intensity strongly correlated with GLUT1 mRNA and protein expression across examined ESCC cell lines (Figure S3D), indicating that SE-driven transcriptional activation leads to aberrant GLUT1 expression in ESCC.

We then performed a series of functional experiments to interrogate the functional consequences of GLUT1 in both human and murine ESCC tumor cells. High GLUT1 expression was detected in 8 of the 12 ESCC cell lines examined, including KYSE140, T.T, KYSE70, and TE-1 (Figure S4A). Transcriptome sequencing analysis using wild-type (WT) and *GLUT1*-KO TE-1 cells showed that 294 genes were commonly downregulated after *GLUT1* knockout (two independent gRNA targets exhibited high consistency) when compared with the WT control (Figure 3A). Further KEGG enrichment analysis of the downregulated genes revealed cellular senescence as the most significantly affected pathway by *GLUT1* knockout, followed by aging-related diseases, such as Parkinson’s, Alzheimer’s, and Huntington’s diseases. Moreover, most genes in these pathways have been reported to function as anti-senescence regulators, including AKT1, FOXM1, and HIPK1 (Figure 3B), suggesting that GLUT1 may contribute to the resistance of ESCC cells to cellular senescence.

**Figure 3.**
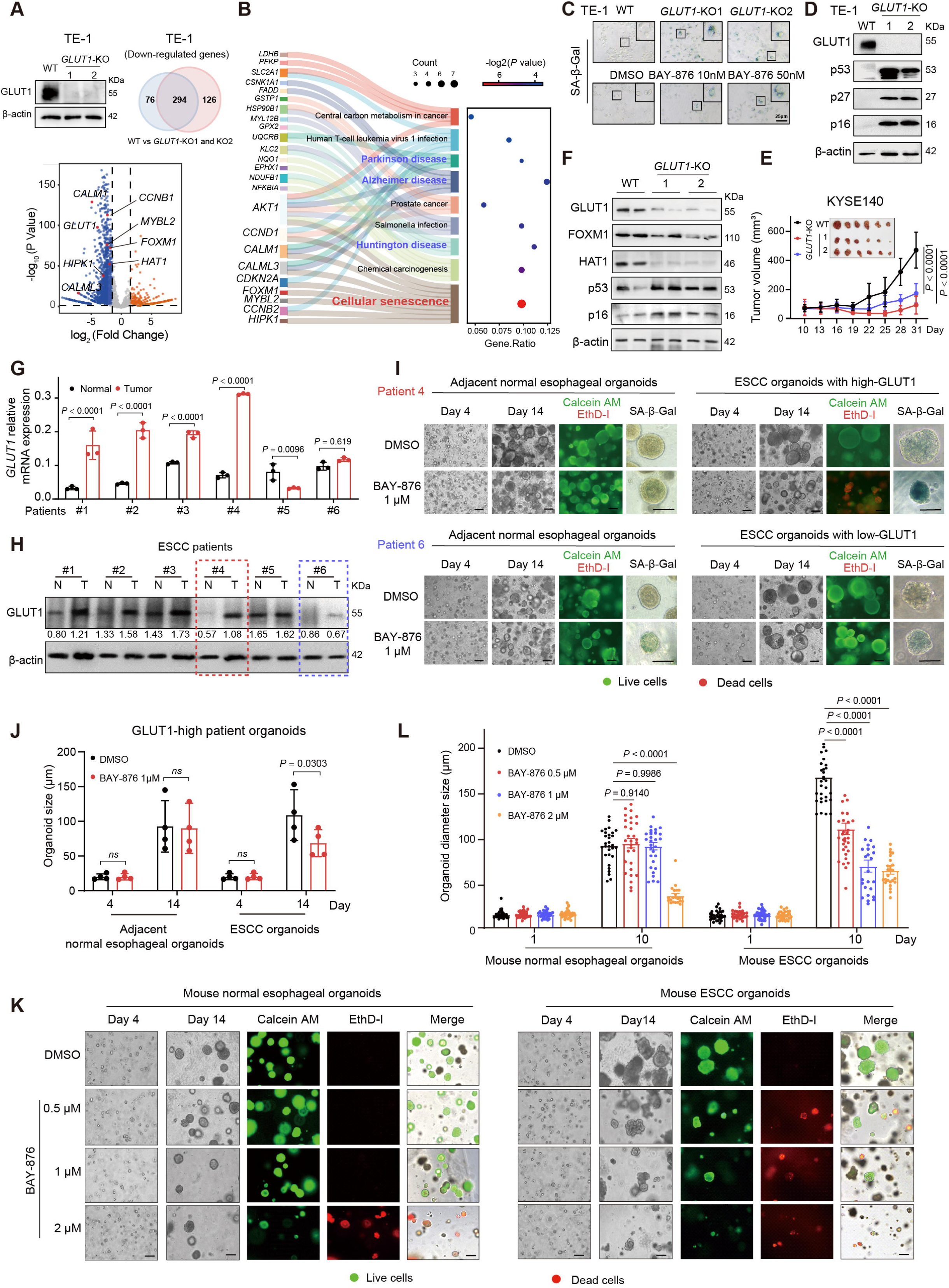
GLUT1 inhibition induces cellular senescence and suppresses ESCC tumor growth. (A) Western blot validation of *GLUT1* knockout (KO) in TE-1 cells. Venn diagram showing commonly downregulated genes in two independent *GLUT1*-KO cells compared to wild-type (WT) cells. Volcano plot showing differentially expressed genes following *GLUT1* depletion. (B) KEGG pathway enrichment analysis of downregulated genes. (C) Representative images of senescence-associated β-galactosidase (SA-β-Gal) staining in TE-1 cells after *GLUT1* knockout or treatment with the GLUT1 inhibitor BAY-876. (D) Western blot analysis showing protein expression of senescence markers p53, p27, and p16 following *GLUT1* depletion. (E) Tumor growth curves and representative images of xenografts derived from WT and *GLUT1*-KO KYSE140 cells in BALB/c nude mice. Each mouse was subcutaneously inoculated with 2 × 10^6^ cells. N = 6. (F) Western blot analysis of protein expression in xenograft tumor tissues in (E). (G and H) GLUT1 expression in paired tumor and adjacent normal tissues in six ESCC patients, measured at the mRNA (G) and protein (H) levels. (I) Representative images of patient-derived organoids (PDOs) with high or low GLUT1 expression treated with BAY-876. Live/dead staining shows Calcein AM (green) and EthD-I (red), along with representative SA-β-Gal staining images. (J) Quantification of PDO size following BAY-876 treatment in GLUT1-high organoids in (I, upper panel). (K and L) Representative images (K) and quantifications (L) of mouse WT and ESCC organoids treated with BAY-876. Data are presented as mean ± SEM, and statistical comparisons were performed using one-way ANOVA with Tukey’s multiple comparison test (E, G, J, and L).

Indeed, GLUT1 deletion or pharmacological inhibition with the GLUT1-selective compound BAY-876 induced a pronounced senescent phenotype, characterized by enlarged, flattened morphology, increased SA-β-Gal activity, and higher expression of senescence markers (p53, p27, p16) (Figure 3C and 3D). Moreover, deletion of *GLUT1* markedly inhibited cell proliferation, invasion, and colony formation in *GLUT1*-high human TE-1 and KYSE140, as well as murine HNM007 cells (Figure S4B-S4E). Conversely, overexpression of *GLUT1* enhanced these malignant phenotypes in *GLUT1*-low KYSE30 and KYSE450 cells (Figure S4F-S4G). Importantly, these effects were faithfully restored by exogenous overexpression of *GLUT1* in *GLUT1*-knockout (KO) cells (Figure S5A-S5I). Furthermore, pharmacological inhibition with BAY-876 suppressed malignant phenotypes and induced senescence (Figure S5J-S5P), indicating that senescence induction by targeting GLUT1 reduces cellular oncogenicity of ESCC.

The regulatory effect of GLUT1 on the malignant phenotype of ESCC was exerted through canonical glycolysis, as the enhanced oncogenicity was significantly abolished upon treatment with the glycolytic inhibitor 2-deoxy-D-glucose (2-DG) in GLUT1-overexpressing cells (Figure S5Q-S5W).

Consistent with the *in vitro* findings, *GLUT1*-KO tumors exhibited significantly reduced tumor volume, decreased Ki67 staining, and upregulated expression of senescence-associated proteins compared with the WT counterparts in both murine ESCC xenograft and allograft models (Figures 3E, 3F, and S5X). Given the clinical relevance of GLUT1 expression described above, we generated patient-derived organoids (PDOs) from ESCC patients and validated the induction of cellular senescence with the GLUT1 inhibitor, BAY-876. Compared with corresponding normal organoids, elevated expression of GLUT1 at both mRNA and protein levels was observed in 4 out of 6 PDOs (Figure 3G and 3H). Notably, tumor organoids with higher GLUT1 expression (from patient 4) exhibited greater sensitivity to BAY-876 and a more pronounced senescent phenotype relative to its normal organoids. However, such a difference was not observed in organoids derived from patient 6, which had low expression of GLUT1 (Figure 3H-3J). Consistent results were obtained in mouse tissue-derived organoids. Low-dose BAY-876 triggered a remarkable senescent phenotype and significantly inhibited the growth of ESCC tumor organoids, while exerting no obvious effect on normal esophageal organoids (Figures 3K, 3L, and S5Y). These results demonstrate that GLUT1 suppresses cellular senescence in ESCC to facilitate tumor progression, and suggest that high expression of GLUT1 may serve as a potential predictor of sensitivity to BAY-876 treatment.

### GLUT1 forms a positive feedback loop with HAT1 and FOXM1 cooperatively promoting chromatin accessibility to antagonize tumor cell senescence

To elucidate the molecular mechanism by which GLUT1 regulates tumor cell senescence, we verified the expression of anti-senescence genes identified from our transcriptome data in *GLUT1*-KO TE-1 and KYSE140 cells (Figure 3B). As expected, *GLUT1* deletion resulted in significant downregulation of these established anti-senescence genes, including *FOXM1*, *HIPK1*, and *AKT1* (Figures 4A, 4B, S6A, and S6B). In addition, we found that HAT1, a histone acetyltransferase implicated in chromatin remodeling and transcription factor recruitment, was also downregulated following *GLUT1* deletion (Figures 4A, 4B, S6A, and S6B). Interestingly, we noticed that Acetyl-CoA (Acetyl Coenzyme A), the essential acetyl group donor required for HAT1 enzymatic activity ^33,34^, is derived from glycolysis-generated pyruvate downstream of GLUT1-mediated glucose uptake. We therefore hypothesized that GLUT1 may fuel the FOXM1-driven transcription program through modulating Acetyl-CoA production and subsequent HAT1-dependent epigenetic remodeling, thereby conferring resistance to cellular senescence (Figure 4C).

**Figure 4.**
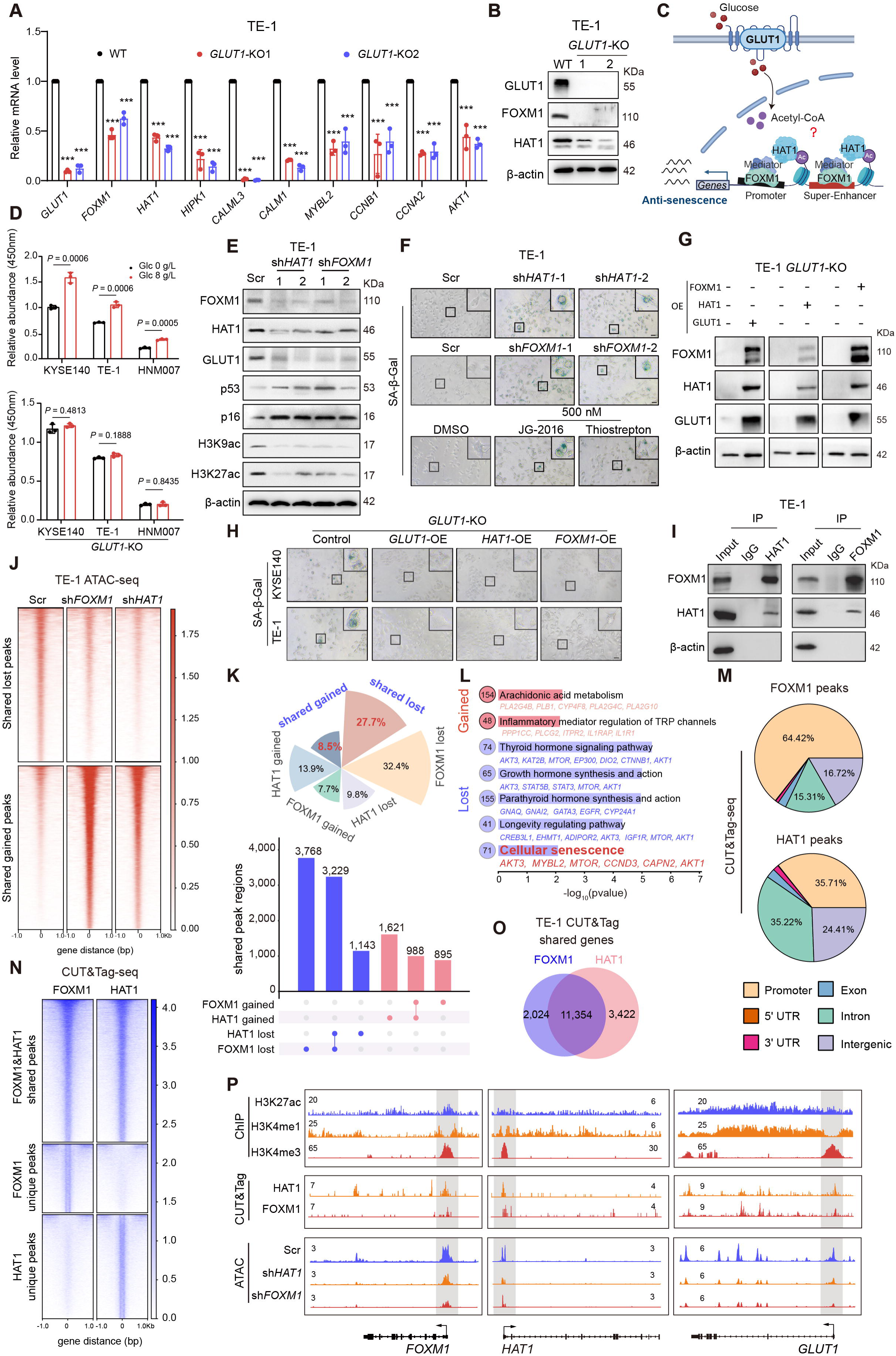
GLUT1 regulates cellular senescence through forming an epigenetic positive feedback loop with HAT1 and FOXM1. (A) RT-qPCR analysis of senescence-associated gene expression following *GLUT1* knockout. (B) Western blot analysis showing reduced FOXM1 and HAT1 protein levels upon *GLUT1* depletion. (C) Schematic model illustrating the regulation of the HAT1-FOXM1 signaling axis by GLUT1. (D) Intracellular Acetyl-CoA levels measured in WT and *GLUT1*-KO KYSE140, TE-1, and HNM007 cells cultured with or without glucose (8 g/L) for 12 h. (E) Western blot analysis of downstream targets following *HAT1* or *FOXM1* knockdown in TE-1 cells. (F) Representative SA-β-Gal staining images following *HAT1* or *FOXM1* knockdown or pharmacological inhibition with JG-2016 or Thiostrepton. (G and H) Western blot analysis (G) and SA-β-Gal staining images (H) showing protein levels and SA-β-Gal activity by exogenous overexpression of *GLUT1*, *HAT1*, or *FOXM1* in *GLUT1*-KO cells. (I) Co-immunoprecipitation showing the interaction between HAT1 and FOXM1. (J) ATAC-seq heatmaps showing chromatin accessibility changes following *FOXM1* or *HAT1* knockdown. Signals are centered at peak summits and extended ±1 kb. (K) UpSet plot and pie chart showing the overlap of chromatin accessibility changes regulated by FOXM1 and HAT1 based on ATAC-seq analysis. Bars indicate the number of genomic regions with increased or decreased accessibility upon FOXM1 or HAT1 knockdown, and the pie chart summarizes the proportions of shared gain, shared loss, and FOXM1- or HAT1-specific changes. (L) KEGG pathway enrichment analysis of genes associated with shared accessibility changes. (M) CUT&Tag-seq analysis showing genome-wide binding profiles, signals are centered at peak summits and extended ±1 kb. (N) Genomic distribution of FOXM1- and HAT1-binding peaks in TE-1 cells. (O) Venn diagram showing the overlap target genes bound by FOXM1 and HAT1. (P) IGV tracks showing ChIP-seq, ATAC-seq, and CUT&Tag signals at *FOXM1*, *HAT1*, and *GLUT1* loci in TE-1 cells. Data are presented as mean ± SEM of three independent experiments, and statistical comparisons were performed using one-way ANOVA with Tukey’s multiple comparison test (A and D).

To test this, we first examined intracellular production of Acetyl-CoA. Indeed, glucose supplementation enhanced glucose uptake and markedly increased Acetyl-CoA levels in GLUT1-high ESCC cells when compared with the non-glucose group. However, this difference was abolished following *GLUT1* deletion (Figure 4D), indicating that glucose-derived Acetyl-CoA production is GLUT1-dependent. Functionally, either the knockdown of *HAT1* or *FOXM1*, or the pharmacological inhibition of HAT1 or FOXM1 using small-molecule inhibitor JG-2016 (for HAT1) or Thiostrepton (for FOXM1), recapitulated the effect of *GLUT1*-KO by significantly suppressing ESCC cell proliferation and colony formation (Figure S6C-S6N). In addition, the downregulation or inhibition of HAT1 or FOXM1 triggered cellular senescence, as shown by suppressed expression of anti-senescence genes (Figure S7A-S7D), increased levels of senescence-associated proteins such as p53 and p16, and elevated SA-β-Gal activity (Figures 4E, 4F, S7E, and S7F).

Nuclear HAT1 is known to catalyze histone acetylation at specific residues, notably H3K9 and H3K27, which are critical for transcriptional activation ^34,35^. Western blot analysis showed that the protein levels of H3K9ac and H3K27ac were substantially decreased following the deletion of *GLUT1*, *HAT1*, or *FOXM1* (Figures 4E, S7E, and S7G). Notably, re-expression of *GLUT1*, *FOXM1*, or *HAT1* in *GLUT1*-KO cells not only effectively reversed the corresponding senescent phenotype, but also restored the expression of itself and the other two genes (Figures 4G, 4H, S7H, and S7I). Together, these results suggest that high expression of GLUT1 activates the anti-senescent HAT1/FOXM1 axis; in turn, FOXM1 and HAT1 may form a positive feedback loop with GLUT1 to maintain its high expression, thereby sustaining the hypermetabolic and anti-senescent state to promote ESCC progression.

FOXM1 is an established transcription factor known to suppress cellular senescence ^36,37^. To ascertain whether FOXM1 functionally cooperates with HAT1 in regulating chromatin accessibility and the expression of downstream anti-senescence genes, we conducted a series of analyses. The co-immunoprecipitation (Co-IP) and IF assays first confirmed the physical interaction between HAT1 and FOXM1, as well as their co-localization with the active histone marks H3K9ac and H3K27ac (Figures 4I and S7J-S7L). ATAC-seq analysis revealed that 36.2% of gained and lost peaks were shared following the downregulation of *HAT1* or *FOXM1* (Figure 4J and 4K). KEGG analysis identified cellular senescence as a significantly enriched pathway of those genes associated with lost accessibility, such as the aforementioned anti-senescent genes *CCND3* and *AKT1* (Figure 4L). Moreover, CUT&Tag profiling identified 11,354 genes co-bound by HAT1 and FOXM1 (Figure 4M-4O). Integration of these data with ChIP-seq profiles for active histone markers further revealed the co-occupancy of HAT1 and FOXM1 around active promoter and enhancer regions, including *GLUT1* and their own gene loci. Importantly, their chromatin accessibility (Figure 4P) and subsequent expression (Figures 4E and S7A-S7E) were consistently decreased upon downregulation of either *HAT1* or *FOXM1*.

Collectively, these results demonstrate that GLUT1 forms a positive feedback loop with HAT1 and FOXM1, cooperatively opening chromatin to drive their own expression, thereby resisting cellular senescence and enhancing the malignant phenotype of ESCC cells.

### Disruption of the GLUT1-HAT1/FOXM1 axis triggers tumor cell senescence to potentiate CD8⁺ T cell-based antitumor immunity

To investigate whether GLUT1 modulates CD8⁺ T cell cytotoxicity via the HAT1/FOXM1 anti-senescence axis, we measured the regulatory effects of HAT1 and FOXM1 on the CD8⁺ T cell-mediated tumor killing by co-culturing HNM007 cells and CD8⁺ T cells derived from OT-I mice. As noted above, *GLUT1* knockout in HNM007 cells markedly enhanced CD8⁺ T cell-mediated tumor killing (Figures 2H and S2F-S2H). Importantly, the enhanced cytotoxicity of CD8⁺ T cells was largely abrogated upon the restoration of *GLUT1*, *HAT1*, or *FOXM1* expression. Accordingly, tumor cell survival was significantly increased, accompanied by decreased proportions of GZMB^+^ and IFNγ^+^ CD8⁺ T cells (Figures 5A and S8A-S8C), demonstrating that the GLUT1-HAT1/FOXM1 signaling axis restrains the antitumor activity of CD8⁺ T cells.

**Figure 5.**
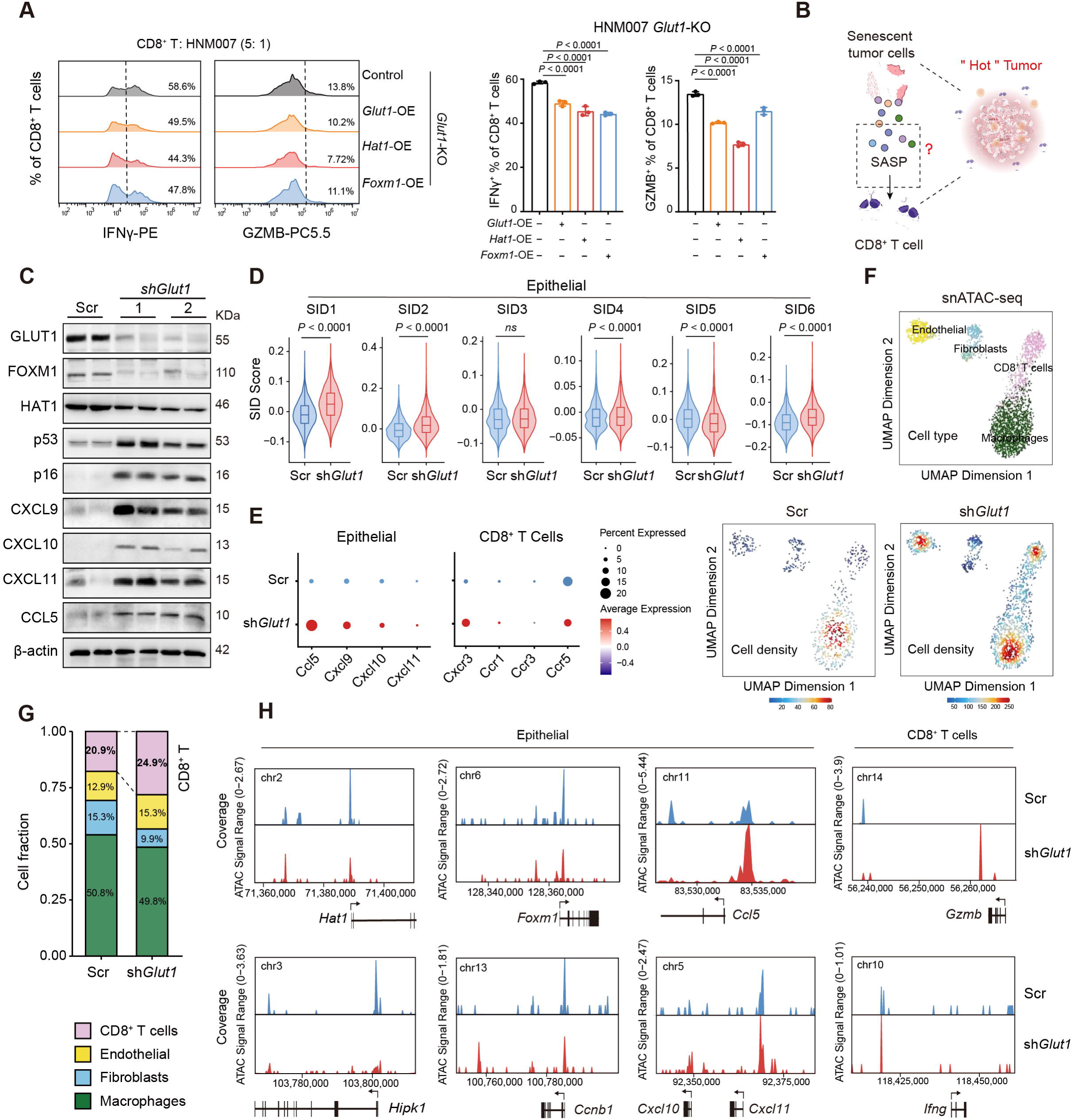
GLUT1 depletion promotes CD8⁺ T cell cytotoxicity. (A) Flow cytometry (left) and statistical analysis (right) showing the frequencies of GZMB⁺ and IFNγ⁺ CD8⁺ T cells in the indicated groups. Statistical comparisons were performed using one-way ANOVA with Tukey’s multiple comparison test. (B) Schematic illustration depicting the secretion of senescence-associated secretory phenotype (SASP) factors by senescent tumor cells, which may promote the recruitment and activation of CD8⁺ T cells within the tumor microenvironment. (C) Western blot analysis of tumor tissues from control and sh*Glut1* mouse allografts in (Figure 2A). (D) Violin plots showing the distribution of major senescence identities (SID) scores (SID1-SID6) in epithelial cells either from Scr or sh*Glut1* mouse allografts. Statistical significance between groups was assessed using a two-sided Student’s t-test. *P* values are indicated above each panel. (E) Dot plots showing the indicated chemokine expression in epithelial cells and CD8⁺ T cells in Scr and sh*Glut1* allograft groups. (F) UMAP plot of non-epithelial cell types identified in mouse allograft tissues via snATAC-seq, and density UMAP plots depict the distinct distribution patterns of the Scr and sh*Glut1* groups. (G) Stacked plot showing the proportion changes for chromatin accessibility in non-epithelial cell types of (F). (H) Aggregate snATAC-seq browser tracks showing *Glut1* knockdown-induced changes in chromatin accessibility at anti-senescence-associated loci, including *Hat1*, *Foxm1*, *Hipk1*, and *Ccnb1*, and SASP-related loci, including *Cxcl10*, *Cxcl11*, and *Ccl5*, in epithelial cells, as well as at effector gene loci, including *Gzmb* and *Ifng*, in CD8⁺ T cells.

Prompted by these findings, we propose that GLUT1 downregulation in ESCC triggers cellular senescence and the subsequent SASP factor release, thereby further recruiting and activating CD8⁺ T cells (Figure 5B). This hypothesis is supported by previous studies, in which CXCL9, CXCL10, CXCL11, and CCL5 have been defined as key SASP components governing CD8⁺ T cell recruitment in the contexts of antitumor immunity and autoimmunity ^38–40^. Consistent with these findings, our western blot analysis verified significant upregulation of these SASP chemokines including CXCL9, CXCL10, CXCL11, and CCL5, as well as senescence markers (p53, p16) in *Glut1*-knockdown allograft tumors compared with control tumors (Figure 5C). Moreover, knockdown of *HAT1* or *FOXM1* recapitulated these molecular alterations in both human and murine cell lines (Figure S8D-S8I), further validating that the GLUT1-HAT1/FOXM1 axis regulates SASP chemokines associated with CD8⁺ T cell recruitment.

Furthermore, we utilized a previously established machine learning-based tool, Senescent Cell Identification (SenCID) ^41^, to analyze our scRNA-seq data derived from murine allograft tumors *in vivo* (Figure 2). Among the six major senescence identities (SIDs) defined by SenCID, knockdown of *GLUT1* markedly elevated the SID scores of four senescence subtypes compared with the control group (Figure 5D). Further analyses demonstrated increased expression of the SASP ligands *Ccl5*, *Cxcl9*, *Cxcl10*, and *Cxcl11* in *Glut1*-knockdown tumor epithelial cells. Meanwhile, their corresponding receptors (*Cxcr3*, *Ccr1*, *Ccr3*, *Ccr5*) were concurrently elevated in tumor-infiltrating CD8⁺ T cells (Figure 5E), establishing a molecular basis for strengthened chemokine-mediated cytotoxicity of CD8⁺ T cells.

To further verify these findings, single-nucleus ATAC-seq (snATAC-seq) was performed using murine allograft tumor tissues. Cell-type annotation and compositional analysis revealed an increased proportion of CD8⁺ T cells following *Glut1* knockdown (Figure 5F and 5G). Consistently, Integrative Genomics Viewer (IGV) visualization showed that *Glut1* knockdown reduced chromatin accessibility at anti-senescence-associated genes (e.g., *Hat1*, *Foxm1*, *Hipk1*, *Ccnb1*) in epithelial cells, while increasing chromatin accessibility at SASP-related genes (e.g., *Cxcl10*, *Cxcl11*, *Ccl5*) in epithelial cells, and at effector genes (*Gzmb* and *Ifng*) in CD8⁺ T cells (Figure 5H). Together, these results demonstrate that repression of the GLUT1-HAT1/FOXM1 signaling axis triggers tumor cell senescence and SASP secretion, thereby facilitating the recruitment, activation, and cytotoxicity of CD8⁺ T cells.

### Targeting GLUT1 sensitizes ESCC to α-PD1 immunotherapy

Given the crucial role of GLUT1 in suppressing cellular senescence to restrain cytotoxic function of CD8⁺ T cells in the ESCC TME, we next investigated whether GLUT1 inhibition could enhance the therapeutic efficacy of α-PD1 blockade, a strategy known to reinvigorate CD8⁺ T cell function. To this end, HNM007 cells were inoculated into syngeneic immunocompetent C57BL/6J mice. Once tumors reached an appropriate volume, the tumor-bearing mice were treated with BAY-876, α-PD1, or the combination of BAY-876 and α-PD1 (Figure 6A).

**Figure 6.**
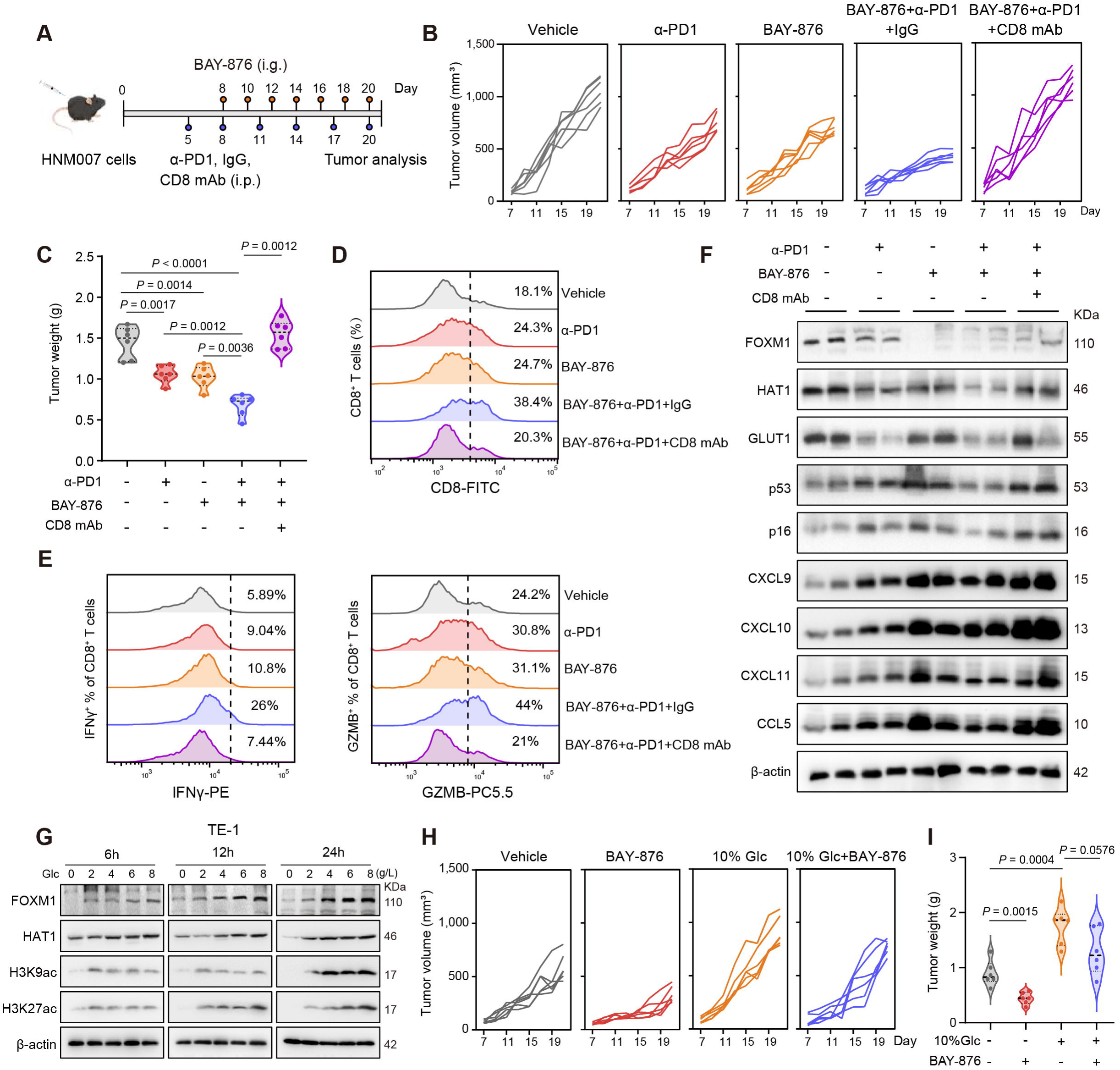
GLUT1 inhibition enhances α-PD1 immunotherapy through CD8⁺ T cell dependent antitumor immunity. (A) Experimental scheme of the *in vivo* treatment regimen. C57BL/6J mice were subcutaneously injected with 5 × 10⁵ HNM007 cells and subsequently treated with vehicle, BAY-876, α-PD1, or their combination (BAY-876 + α-PD1) with or without CD8-depleting antibody according to the indicated schedule. (B) Individual tumor growth curves for each treatment group. Data are presented as mean ± SEM, and statistical comparisons were performed using one-way ANOVA with Tukey’s multiple comparison test. N = 6. (C) Tumor weights at the study endpoint in (B). Each dot represents an individual tumor. *P* values indicate statistical significance between the indicated groups, as determined by one-way ANOVA followed by Tukey’s multiple comparisons test. (D) Flow cytometry quantification of tumor-infiltrating CD8⁺ T cells. (E) Assessment of CD8⁺ T cell effector function by IFNγ and GZMB expression. (F) Western blot analysis showing expression levels of FOXM1, HAT1, GLUT1, p53, p16, and SASP chemokines CXCL9, CXCL10, CXCL11, and CCL5 in tumor tissues. (G) Western blot analysis of FOXM1, HAT1, H3K9ac, and H3K27ac expression in TE-1 cells cultured with increasing glucose concentrations (Glc, 0-8 g/L) at the indicated time points (6 h, 12 h, and 24 h). (H and I) Tumor growth curves (H) and tumor weights (I) in mice under different glucose conditions, with or without BAY-876 treatment. Data are presented as mean ± SEM, and statistical comparisons were performed using one-way ANOVA with Tukey’s multiple comparison test. N = 6.

As shown in Figures 6 and S9A, monotherapy with either BAY-876 or α-PD1 yielded moderate suppression of HNM007 tumor growth. However, their combination significantly retarded tumor progression, demonstrating a robust synergistic antitumor effect (Figures 6B, 6C, and S9A). Consistently, IF staining and flow cytometry analysis further validated that combined treatment with BAY-876 and α-PD1 markedly enhanced CD8⁺ T cell infiltration and elevated the frequencies of IFNγ^+^ and GZMB^+^ CD8⁺ T cells within tumor tissues relative to monotherapy (Figures 6D, 6E, and S9B). Moreover, western blot analysis confirmed that BAY-876 treatment decreased the expression of anti-senescence factors FOXM1 and HAT1, while increasing protein levels of the senescence markers p53 and p16, as well as the SASP chemokines responsible for CD8⁺ T-cell recruitment (Figure 6F), supporting the activation of senescence-initiated antitumor immune responses. However, the potent antitumor efficacy of BAY-876 and α-PD1 combination was completely abrogated upon CD8⁺ T cell depletion using a specific CD8 neutralizing antibody (Figure 6B-6F), verifying that CD8⁺ T cells are indispensable for the therapeutic response. These results demonstrate that GLUT1 inhibition disrupts the HAT1/FOXM1 axis to trigger tumor cell senescence, potentiating CD8⁺ T cell-dependent antitumor immunity and sensitizing ESCC tumors to α-PD1 immunotherapy.

### Dietary glucose restriction potentiates the antitumor function of CD8⁺ T cells potentially via modulating GLUT1-HAT1/FOXM1 axis

Given that GLUT1 functions as a major glucose transporter and current small-molecule inhibitors exhibit suboptimal specificity, we hypothesized that modulation of glucose availability may serve as a viable alternative strategy to target the GLUT1-HAT1/FOXM1 metabolic axis and potentiate antitumor immunity. We initially cultured TE-1, KYSE140, HNM007, and AKR cells in glucose-deprived medium with gradient glucose supplementation. Glucose stimulation concentration- and time-dependently upregulated the expression of HAT1, FOXM1, H3K9ac, and H3K27ac (Figures 6G and S10A-S10G), whereas this inductive effect was abrogated in *GLUT1*-KO cells (Figure S10H-S10J). Furthermore, SA-β-Gal activity in senescent ESCC cells declined gradually as glucose concentrations increased (Figure S10K). These data suggest that modulating glucose availability can recapitulate GLUT1 expression-dependent senescent phenotypes and core molecular alterations of the GLUT1-HAT1/FOXM1 axis.

Consistently, high glucose promoted ESCC tumor growth *in vivo*, as evidenced by increased tumor volume and weight in tumor-bearing mice administered 10% glucose in drinking water compared with those receiving 1% glucose (Figure S11A-S11C). Following the treatment with the GLUT1 inhibitor BAY-876, tumor growth was mildly suppressed in the 10% glucose group, yet this inhibitory effect was negligible relative to the normal glucose control group (Figures 6H, 6I, S11D, and S11E). Subsequent IF staining, western blot, and flow cytometry analyses confirmed that high glucose intake elevated HAT1 and FOXM1 expression, while reducing protein levels of the senescence markers (p53 and p16), as well as SASP chemokines (CXCL9, CXCL10, CXCL11 and CCL5) responsible for CD8⁺ T cell recruitment, thereby impairing CD8⁺ T cell infiltration and activation (Figures S11F-S11H).

Low-carbohydrate diet (LCD) has evolved from mere nutritional support to an active combinatorial therapeutic strategy, owing to its potential to alleviate metabolic competition within the TME and improve treatment tolerance ^42^. To explore the functional implication of the LCD strategy in our study, mice were randomly assigned to three groups fed a standard diet (SD), an LCD, or an LCD supplemented with 10% glucose, respectively. Compared with the SD group, the LCD group exhibited significantly suppressed tumor growth, accompanied by downregulated expression of HAT1 and FOXM1, elevated levels of cellular senescence markers, and enhanced infiltration and activation of CD8⁺ T cells. Notably, these favorable antitumor effects were largely abrogated by 10% glucose supplementation (Figures 7A, 7B, and S11I-S11M), indicating that dietary glucose indeed modulates the HAT1/FOXM1 anti-senescent axis, as well as the subsequent infiltration and activation of CD8⁺ T cells.

**Figure 7.**
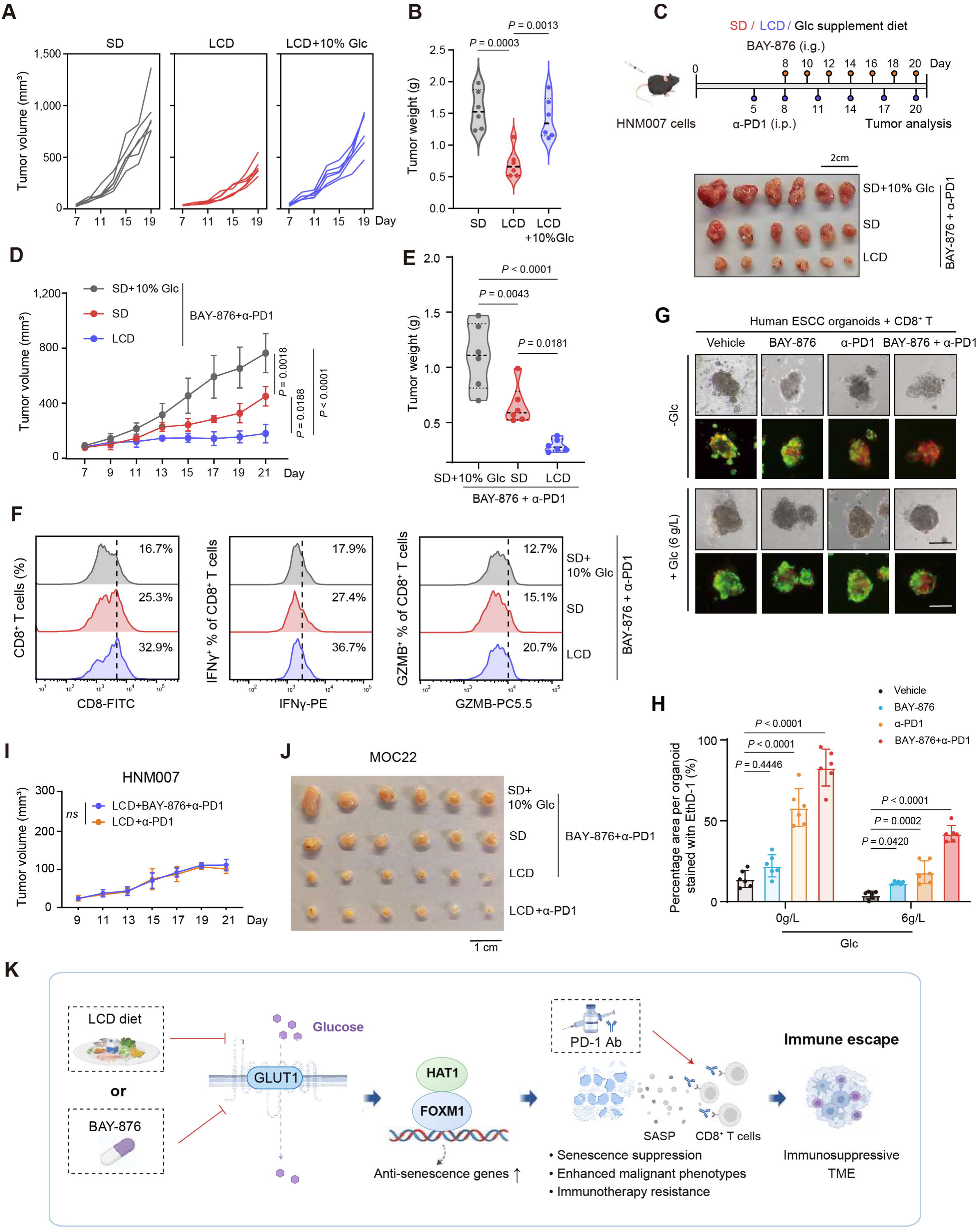
Dietary glucose restriction potentiates the antitumor function of CD8⁺ T cells via the modulating GLUT1-HAT1/FOXM1 anti-senescence axis. (A and B) Tumor growth curves (A) and tumor weights (B) in mice subjected to dietary glucose interventions including standard diet (SD), low-carbohydrate diet (LCD), or 10% glucose (Glc) supplementation. Data are presented as mean ± SEM, and statistical comparisons were performed using one-way ANOVA with Tukey’s multiple comparison test. (C) Experimental schematic of the dietary glucose interventions and treatment regimens (left). C57BL/6J mice were subcutaneously injected with 5 × 10⁵ HNM007 cells and treated with BAY-876 and α-PD1 antibody under SD, LCD, or glucose supplemented conditions. Photographs of isolated tumors on day 20 (right). (D and E) Tumor volumes (D) and tumor weight (E) following combination therapy under different dietary conditions. Data are presented as mean ± SEM, and statistical comparisons were performed using one-way ANOVA with Tukey’s multiple comparison test. N = 6. (F) Flow cytometry analysis of CD8⁺ T cell infiltration and effector function by IFNγ and GZMB expression under different dietary conditions. (G) Representative bright-field and fluorescence images of ESCC patient-derived organoids (PDOs) co-cultured with CD8⁺ T cells under glucose deprivation (-Glc) or + Glc (6 g/L) conditions and treated with vehicle, BAY-876, α-PD1, or their combination. Live or dead cells were respectively stained with Calcein AM (green) and EthD-I (red). Scale bar, 100 μm. (H) Quantification of the EthD-1-positive area percentage per organoid in (G). (I) Tumor growth curves of HNM007 tumor-bearing mice treated with α-PD1 or BAY-876 + α-PD1 under LCD conditions. Data are presented as mean ± SEM, and statistical comparisons were performed using one-way ANOVA with Tukey’s multiple comparison test. N = 6. (J) Representative tumor images from MOC22 oral squamous cell carcinoma tumor model under different dietary and treatment conditions (10% glucose, SD, LCD, and LCD + α-PD1) with or without BAY-876 + α-PD1. (K) Proposed model illustrating the role of the GLUT1-HAT1/FOXM1 axis in regulating tumor senescence and immune escape in ESCC.

### Dietary glucose restriction improves immunotherapy efficacy

We next evaluated the efficacy of combination therapy with BAY-876 and α-PD1 under different dietary regimens. The LCD + BAY-876 + α-PD1 group achieved the most robust antitumor response, followed by the SD + BAY-876 + α-PD1 group, whereas the SD + 10% glucose + BAY-876 + α-PD1 group showed the poorest therapeutic efficacy (Figures 7C-7E and S11N). Further mechanistic analysis demonstrated that the LCD + BAY-876 + α-PD1 regimen most effectively triggered tumor cell senescence and SASP secretion, thereby maximally enhancing CD8⁺ T cell infiltration and activation (Figures 7F, S11O, S12A).

In light of recent evidence indicating that a low-glucose diet may remodel the pulmonary microenvironment via modulating NK cells and macrophages, thereby facilitating metastatic dissemination ^43^, we obtained lung tissues from the experimental mice mentioned above. Flow cytometry analysis revealed no significant differences in the abundance of NK cells or macrophages within the lungs among the three groups (Figure S12B). Nevertheless, small metastatic foci were detected in the lungs of mice fed a high-glucose diet (10% glucose + BAY-876 + α-PD1) (Figure S12C). Concurrently, we performed gene set variation analysis to evaluate the effect of glucose metabolic activity on postoperative 2-year recurrence in ESCC patients (from the TCGA-ESCC dataset) using the KEGG glycolysis/gluconeogenesis pathway hsa00010. Consistent with the previous results ^43^, low glucose metabolic activity was not correlated with a 2-year postoperative recurrence in patients with ESCC. In contrast, high glucose metabolic activity showed a potential trend toward increased recurrence risk (Figure S12D). These results indicate that a low-glucose diet does not promote lung metastasis in the clinical setting of ESCC.

To validate the clinical relevance of our findings, we established a co-culture system consisting of ESCC PDOs and matched autologous peripheral blood mononuclear cells (PBMCs). We then evaluated the cytotoxic effects induced by BAY-876, α-PD1 or their combination under glucose-deprived or 6 g/L glucose conditions. Overall, stronger cytotoxic effects were observed under glucose-deprived conditions compared with the 6 g/L glucose groups. Notably, the combination of BAY-876 and α-PD1 elicited the most pronounced tumor cell death under glucose deprivation (Figures 7G, 7H, and S12E). Consistent results were obtained in both murine ESCC and oral squamous cell carcinoma organoids generated from 4-NQO-induced primary tumors. Flow cytometry analysis further demonstrated a marked increase in GZMB^+^ and IFNγ^+^ cytotoxic populations in the combination treatment group under glucose-deprived conditions (Figure S12F-S12I).

To test whether dietary glucose restriction could replace pharmacological GLUT1 inhibition *in vivo*, ESCC tumor-bearing mice were subjected to LCD feeding and α-PD1 treatment with or without BAY-876 administration. Remarkably, LCD yielded comparable tumor suppression effect to BAY-876 when combined with α-PD1 (Figures 7I, S13A, and S13B). Consistently, flow cytometry analysis showed no significant alterations in the frequency or activation status of CD8⁺ T cells between the two treatment arms (Figure S13C). These results suggest that GLUT1 inhibition does not act synergistically with glucose restriction; rather, the two interventions converge functionally within the identical metabolic-senescence pathway, representing a pattern of vertical blockade.

As a validation, we repeated the *in vivo* experiments using the murine oral squamous cell carcinoma cell line MOC22. Of note, MOC22 tumors were sensitive to combined treatment with BAY-876 and α-PD1 across all four groups. Nevertheless, the combination of BAY-876 and α-PD1 in LCD-fed mice still yielded the most favorable tumor-suppression trend, consistent with our observations in the ESCC model. Importantly, α-PD1 combined with LCD achieved comparable antitumor efficacy to the regimen containing BAY-876 (Figures 7J and S13D), supporting no additive effect between glucose restriction and GLUT1 inhibition. Collectively, our findings highlight dietary glucose restriction as a viable alternative to GLUT1 inhibition for potentiating immunotherapy efficacy by targeting the GLUT1-HAT1/FOXM1 anti-senescence axis (Figure 7K).

## Discussion

Both GLUT1-mediated glycolysis ^23,44–46^ and cellular senescence ^19,20^ have been independently confirmed to regulate immune responses in both tumor cells and immune cells. In this study, we demonstrate GLUT1 as a key mediator in ESCC tumor cells connecting glucose metabolism to cellular senescence and antitumor immunity, representing a novel function of GLUT1 beyond glucose transport. Herein, GLUT1 in ESCC tumor cells can remodel TME and suppress adaptive antitumor immune response by resisting cellular senescence, in which nuclear HAT1/FOXM1 anti-senescence axis is a key downstream node of GLUT1 signaling. In GLUT1-high ESCC tumors, anti-senescent genes, including FOXM1, were transcriptionally activated upon glucose uptake, while SASP chemokines responsible for CD8⁺ T cell recruitment, such as CCL5, CXCL10, and CXCL11 were inhibited through remodeling of chromatin accessibility. Deletion of *GLUT1* or treatment with BAY-876, α-PD1 alone or their combination resulted in increased CD8⁺ T cell infiltration and cytotoxic activity in both *ex vivo* PDO co-culture and *in vivo* murine allograft models, particularly under low-glucose condition. Although our study focused on the GLUT1-upregulated HAT1/FOXM1 anti-senescence pathway in T cell-mediated immune responses, the concurrent alterations observed in macrophages and NK cells suggest the involvement of additional, yet uncharacterized mechanisms. Future studies are warranted to delineate how GLUT1-mediated anti-senescence signaling interacts with distinct immune cell populations, either directly or indirectly, to establish an immune-evasive TME.

Our findings carry significant implications for the clinical management of ESCC. We identify GLUT1 as a reliable biomarker to predict non-responsiveness to immunotherapy and unfavorable prognosis in patients with ESCC. Consistently, the significant correlation of high GLUT1 expression with advanced stage, increased invasiveness, and unfavorable overall and disease-free survival has also been reported across multiple solid tumors ^47,48^. In addition, GLUT1-high ESCC cells show higher sensitivity to treatment of BAY-876 in both murine and human organoids relative to their normal counterparts. Moreover, GLUT1 is significantly upregulated in immunotherapy nonresponsive ESCC patients, possessing a superior discriminatory performance relative to PD-L1.

Therapeutically, strategies targeting the GLUT1-regulated immune response hold great promise. Our data demonstrated that combined GLUT1 inhibition and α-PD1 treatment resulted in pronounced antitumor effects in both co-cultured PDOs and murine ESCC tumors, and such effects are more potent under low-glucose conditions compared with high-glucose conditions. Consistently, GLUT1 inhibition prevents lactate excretion from tumor glycolysis, thereby alleviating metabolic immune suppression ^49^; LDH inhibitors selectively suppress tumor glycolysis and GLUT1 expression in tumor cells while enhancing glucose uptake, GLUT1 expression, and proliferation in tumor-infiltrating T cells, owing to the higher basal LDH expression and glycolysis activity of tumor cells compared with infiltrating T cells ^45^. Therefore, high GLUT1 expression and glycolysis levels in tumor cells also create therapeutic opportunity for tumor-specific targeting of glycolysis in ESCC and other squamous cell carcinomas. In addition, targeting the GLUT1-HAT1/FOXM1 axis boosts antitumor immunity by triggering tumor cell senescence, and also offers a mechanistic rationale for how GLUT1 inhibition or a glucose restriction diet enhances immunotherapy. However, to facilitate clinical translation, future preclinical studies and clinical trials should systematically determine how targeting GLUT1-mediated glucose metabolism affects antitumor immunity, therapeutic efficacy, and treatment safety.

A central challenge in therapeutically targeting GLUT1 is its ubiquitous expression and essential function in tissues such as erythrocytes, raising substantial concerns over on-target toxicity ^50,51^. Our study establishes a mechanistic rationale for a viable therapeutic window. In ESCC, GLUT1 deletion or inhibition triggers a cellular senescence program via the HAT1/FOXM1 transcriptional axis. Critically, this nuclear signaling pathway is irrelevant in anucleate erythrocytes, highlighting a cancer-specific signaling role of GLUT1 that is distinct from its canonical metabolic function. Moreover, the profound glycolytic addiction of tumor cells confers a differential vulnerability compared to normal tissues. Supporting this, BAY-876 treatment at efficacious doses induced significant tumor cell death and senescence without causing severe systemic toxicity or marked weight loss in mice. In addition, our findings suggest that dietary glucose restriction represents a feasible alternative to GLUT1 inhibition, given their shared regulatory axis and equivalent antitumor effect in boosting immunotherapy. This strategy will fundamentally overcome the on-target toxicity of GLUT1 inhibition.

Glucose metabolism modulation emerges as a promising anticancer therapeutic strategy that leverages the Warburg effect ^45,49,52,53^. The controversial point is that glucose deprivation promotes distant metastasis by establishing a pre-metastatic niche dominated by lung macrophage in specific murine hepatocellular carcinoma models ^43^. Our study provides direct experimental evidence relevant to the ongoing debate on dietary glucose modulation in cancer therapy. On phenotype, glucose restriction diet (LCD) suppressed ESCC tumor growth in combination with BAY-876 and α-PD1, and no lung metastases were observed. However, small metastatic foci were detected in the lungs of mice under high glucose condition. On cell populations, flow cytometry showed no significant difference for NK and macrophage proportions between high or low glucose groups. On metabolism, gene set variation analysis demonstrated no significant correlation between glucose metabolism activity and a 2-year postoperative recurrence in patients with ESCC. Supporting our results, such an association has not been previously observed in ESCC and another squamous cell carcinoma ^43^. These results highlight the substantial metabolic heterogeneity across tumor types. In the context of ESCC, our findings not only support the glucose restriction paradigm to enhance immunotherapy but also emphasize the necessity for context-specific metabolic investigations.

### Limitations of the study

We revealed a negative correlation between GLUT1 expression and immunotherapy response based on scRNA-seq data from 42 ESCC patients, and further validated this association using tissue specimens from an independent cohort of 25 ESCC patients receiving immunochemotherapy. Nevertheless, the generalizability of GLUT1 as a stratification biomarker for immunotherapy non-response remains to be validated in large clinical cohorts of ESCC and other squamous cell carcinomas. While results from our murine ESCC model and patient-derived ESCC organoid co-culture systems support that dietary glucose restriction potentiates immunotherapy efficacy, optimized immunotherapy strategies based on dietary intervention still require further evaluation in clinical trials. Additionally, given the ubiquitous involvement of GLUT1 and glucose metabolism in tumor progression, our findings merit further translational exploration across pan-cancer settings.

## Methods

### Patient samples

Human tissue microarrays (TMAs) comprised formalin-fixed paraffin-embedded (FFPE) cores from 85 ESCC cases (ESCC cohort 1) and 27 LUSC cases. ESCC cohort 1 tissues were obtained from Linzhou, Henan Province, China, and ethical approval was granted by the Ethics Committee of the National Cancer Center/Cancer Hospital, CAMS & PUMC (No. 16-084/1163). LUSC TMA slides were purchased from Outdo Biotech (Shanghai, China), and the use of these slides was approved by the Clinical Research Ethics Committee of Outdo Biotech (Nos. SHYJS-CP-1407020 and SHXC202IYF01).

ESCC cohort 2 comprised FFPE whole-tissue sections from 25 patients who received neoadjuvant immunochemotherapy at the First Affiliated Hospital of Anhui Medical University and Hefei Cancer Hospital, Chinese Academy of Sciences. Based on clinicopathological information provided by the pathology department, patients were classified into the pathological complete response group (CR, N = 6) and the non-complete response group (NCR, N = 19). Tissue specimens were preserved in FFPE blocks, sectioned, and mounted on glass slides for subsequent IF analyses. For organoid culture and co-culture assays, biopsy or surgical specimens were obtained from treatment-naive patients with ESCC at the First Affiliated Hospital of Anhui Medical University.

This study was approved by the Ethics Committee of First Affiliated Hospital of Anhui Medical University (no. PJ 2025-06-19 for ESCC cohort 2 and PDOs), and the Ethics Committee of Hefei Cancer Hospital, Chinese Academy of Sciences (no. PJ-KY2024-083 for ESCC cohort 2). Written informed consent was obtained from all patients. The collection and use of all human samples were conducted in accordance with the principles of the Declaration of Helsinki.

### Mouse models

BALB/c nude mice and C57BL/6J mice were purchased from Gem Pharmatech (Nanjing, China). OT-I TCR transgenic mice (C57BL/6-Tg(TcraTcrb)1100Mjb/J) were generously provided by Dr. Zhengfan Jiang (Beijing, China) and housed in a specific pathogen-free facility in Hefei Institutes of Physical Science, Chinese Academy of Sciences (Hefei, China). All animal experiments were performed with the approval of the animal care regulations of Hefei Institutes of Physical Science, Chinese Academy of Sciences (DWLL(E)-2024-37).

For gene function investigation *in vivo*, immunodeficient BALB/c nude mice and immunocompetent C57BL/6J mice were respectively utilized. Wild-type (WT) and *GLUT1* knockout (*GLUT1*-KO) KYSE140 cells were subcutaneously injected into BALB/c nude mice. Scramble control (Scr) and *GLUT1*-knockdown (*GLUT1*-KD) HNM007 cells were injected into immunocompetent C57BL/6J mice, with sh*GLUT1* expression induced by doxycycline (Dox) administration in drinking water. To evaluate whether *GLUT1*-KD enhances the efficacy of PD-1 monoclonal antibody (α-PD1) therapy, tumor-bearing mice were randomly assigned to treatment with either α-PD1 (BE0146, BioXcell) or IgG isotype control (IgG2a, BE0089, BioXcell). To determine whether the tumor-suppressive effects of *GLUT1* inhibition depend on CD8⁺ T cells, a subset of mice received either a CD8⁺ T cell-depleting mAb (BE0061, BioXcell) or IgG isotype control (IgG2b, BE0090, BioXcell) in combination with α-PD1. All antibodies were administered via intraperitoneal injection (100 μg per dose per mouse) every three days for two weeks. For CD8⁺ T cell-depletion experiments, anti-CD8 mAb treatment was initiated on day 5 after tumor cell inoculation.

For *in vivo* experiments involving dietary glucose restriction, mice were randomly assigned to different dietary regimens: a standard diet (SD: 11 kJ% Fat, 22 kJ% Protein, 67 kJ% Carbohydrates), a standard diet supplemented with 10% glucose in drinking water (SD + 10% glucose), or a low-carbohydrate diet (LCD: 16 kJ% Fat, 50 kJ% Protein, 34 kJ% Carbohydrate). All diets were provided by Beijing Keao Xieli Feed Co., Ltd. Specific treatment schedules are detailed in the main figures and corresponding legends.

### Cell lines

KYSE cell line series were generously gifted by Dr. Y. Shimada (Kyoto University, Japan). The AKR and HNM007 cell lines were kindly provided by Dr. Anil K. Rustgi (Columbia University Irving Medical Center, USA). All cell lines were maintained at 37 °C in a humidified atmosphere of 5% CO_2_. Cells were cultured in either RPMI-1640 or Dulbecco’s Modified Eagle Medium (DMEM), supplemented with 10% FBS, penicillin, and streptomycin. All the cell lines were authenticated via short tandem repeat (STR) analysis before experimental manipulation.

For primary CD8⁺ T cells isolation and activation, splenocytes were harvested from OT-I TCR transgenic mice. Briefly, mouse spleen was gently homogenized with the plunger of a 1 mL syringe in complete RPMI 1640 medium containing 10% FBS. Red blood cells were eliminated using ACK Lysis Buffer (Beyotime, C3702), and the resultant splenocytes were stimulated with 300 ng/mL SIINFEKL peptide (OVA_257-264_; Sigma-Aldrich) for 24 h to induce antigen-specific CD8⁺ T cells. The activated cells were further cultured for 48 h in medium containing IL-2 (100 U/mL; Peprotech). CD8⁺ T cells were subsequently obtained using the MojoSort™ Mouse CD8 T Cell Isolation Kit (BioLegend, 480007) and expanded for another 3 days in IL-2-supplemented medium prior to co-culture assays.

### Transient shRNA knockdown by plasmid transfection

For transient knockdown of *GLUT1*, *HAT1*, and *FOXM1*, shRNAs were designed via the GPP Web Portal (https://portals.broadinstitute.org/gpp/public/gene/search). The corresponding oligonucleotides (sequences listed in Supplementary Table 1) were synthesized, annealed, and cloned into the XhoI/HpaI sites of the LentiLox 3.7 vector. For transfection, cells were seeded in 6-well plates and cultured until reaching 40-60% confluence, followed by plasmid transfection using Hieff Trans® Transfection Reagent (YESEN, China) according to the manufacturer’s instructions.

### Generation of stable inducible knockdown cell lines

To generate tetracycline-inducible *GLUT1* knockdown cell lines, non-targeting shRNA (Scramble) and *GLUT1*-specific shRNA were separately inserted into the TRIPZ vector, which was kindly provided by Dr. Boshi Wang (Shanghai Cancer Institute, Renji Hospital, Shanghai Jiao Tong University School of Medicine). Target sequences are provided in Supplementary Table 1. Recombinant lentiviruses were produced by co-transfecting HEK293T cells at 70-90% confluence with TRIPZ constructs together with packaging plasmids psPAX2 and pMD2.G. Supernatants containing lentiviral particles were collected at 48 h post-transfection, filtered through a 0.45-μm filter, and subsequently used for target cell infection. Stable polyclonal cell populations were selected with puromycin (Macklin), and the knockdown efficiency of GLUT1 was verified by RT-qPCR and Western blot.

### Generation of CRISPR/Cas9-mediated knockout cell lines

Single-guide RNAs (sgRNAs) targeting human and mouse GLUT1 were designed using CHOPCHOP (http://chopchop.cbu.uib.no/); target sequences are listed in Supplementary Table 1. Lentiviral particles were packaged in HEK293T cells by co-transfection of the LentiCRISPR v2 vector and packaging plasmids. Target cells were transduced with the viral supernatant and selected with puromycin. Single puromycin-resistant clones were isolated and expanded. Knockout efficiency was validated by Western blot analysis.

### RNA purification and reverse transcriptase quantitative PCR (RT-qPCR)

Total RNA was extracted using the Steady Pure Universal RNA Extraction Kit (Accurate Biology). Complementary DNA (cDNA) was synthesized from the extracted RNA with Evo M-MLV RT Premix (Accurate Biology), following the manufacturer’s protocol. Quantitative real-time PCR (RT-qPCR) was performed on a Roche LightCycler 480 instrument using the SYBR® Green Premix Pro Taq HS qPCR Kit (Accurate Biology). Each reaction was carried out in triplicate, and β-actin was used as an internal reference gene. All primer sequences are listed in Supplementary Table 1.

### Western blot analysis

Protein extracts were prepared using standard protocols and separated by 8-15% SDS-polyacrylamide gel electrophoresis (SDS-PAGE). Separated proteins were then electrotransferred onto 0.22 μm polyvinylidene fluoride (PVDF) membranes (Merck Millipore, Billerica, MA, USA). Membranes were blocked with 5% non-fat milk for 1 h at room temperature and then incubated with primary antibodies overnight at 4 °C. After three washes with TBS-T buffer, membranes were incubated with horseradish peroxidase (HRP)-conjugated secondary antibodies for 1 h at room temperature. Protein bands were visualized using a Tanon 5100 chemiluminescence imaging system (Tanon, Shanghai, China). The following primary antibodies were used: anti-HAT1 (HUABIO, HA722332, 1:1000), anti-FOXM1 (Proteintech, 13147-1-AP, 1:3000), anti-GLUT1 (Proteintech, 21829-1-AP; 1:4000), anti-p53 (Proteintech, 60283-2-Ig, 1:2000), anti-p16 (HUABIO, ET1602-9, 1:5000), anti-CCL5 (Abclonal, A14192, 1:1000), anti-CXCL9 (Abclonal, A1864, 1:1000), anti-CXCL10 (Abclonal, A21986, 1:1000), anti-CXCL11 (Abclonal, A6201, 1:1000), anti-H3K9ac (HUABIO, HA722132, 1:2000), and anti-H3K27ac (Abcam, ab177178, 1:10000). Specific antibody information is provided in Supplementary Table 2.

### RNA-seq and differential expression analysis

Total RNA was isolated from TE-1 cells expressing scramble control (Scr) or *GLUT1-*KO (KO-1 and KO-2) using the Steady Pure Universal RNA Extraction Kit. RNA quality was assessed prior to library preparation. Sequencing libraries were prepared according to the manufacturer’s instructions and sequenced on DNBSEQ-T7 platform. Raw reads were aligned to the human reference genome (hg38) using STAR, and gene-level read counts were generated using featureCounts. KEGG pathway enrichment analysis was performed using clusterProfiler based on the selected gene sets.

### Cell proliferation and colony formation assays

Cells were seeded into 96-well plates at a density of 2,000-5,000 cells per well and cultured in quintuplicate. A 10-μL aliquot of MTT substrate (5 mg/mL) was added to each well and incubated for 4 h at 37 ℃. The formed formazan crystals were solubilized with 10% SDS, and the optical density (OD) was measured at 490 nm. For the colony formation assay, 2,000 cells were plated into each well of 6-well plates and maintained under standard culture conditions for 14 days. The resulting colonies were fixed with methanol, stained with crystal violet, and quantified using ImageJ 1.47v software.

### Cell invasion assay

Cell invasion was evaluated using Matrigel-coated Transwell chambers (8 μm pore size; Corning, USA). Briefly, cells were harvested and resuspended in serum-free medium. A total of 5 × 10⁴ cells in 200 μL serum-free medium were seeded into the upper chamber, while the lower chamber was supplemented with 500 μL complete medium containing 10% FBS as a chemoattractant. After 48 h of incubation at 37 °C, non-invading cells on the upper membrane surface were carefully removed with a cotton swab. The invaded cells on the lower surface were fixed with 4% paraformaldehyde, stained with 0.1% crystal violet, and photographed. The number of invaded cells was quantified by counting five randomly selected fields per membrane under a light microscope.

### Organoid culture

Organoids were generated from normal and tumor tissues following previously established protocols ^54^. Briefly, tissues were rinsed with Hank’s Balanced Salt Solution (HBSS), mechanically minced, and digested in prewarmed digestion buffer. The harvested single cells were resuspended in Matrigel and seeded into 24-well plates. Once the Matrigel droplets solidified, organoid culture medium was added, and fresh medium replenished every other day.

### Live/Dead staining of organoids

Mouse and human esophageal organoids were cultured in 24-well plates for 4 days before drug treatment. On day 10 of culture, the medium was removed, and the organoids were gently washed three times with 1× PBS. The organoids were then stained with 1 µM Calcein-AM and 5 µM Ethidium homodimer-1 (EthD-1, Amersco/UElandy, L6023M) prepared in 1× PBS, and incubated for 30 min at room temperature in the dark. Fluorescence images were acquired using an Olympus SpinSR10 spinning disk confocal super-resolution microscope (Olympus, Japan). The percentage of dead cells (EthD-1-positive, red fluorescence) was quantified by measuring the red fluorescence signal intensity relative to the total organoid area using ImageJ 1.47v software.

### Co-immunoprecipitation (Co-IP)

Cells were lysed on ice for 30 min using NP-40 immunoprecipitation lysis buffer supplemented with 5% PMSF (Beyotime, Shanghai, China). The lysates were sonicated on ice (35% power, 5 min total duration with 5 s ON/10 s OFF cycles) to shear genomic DNA and reduce viscosity. After centrifugation, a portion of the supernatant was collected as the “Input” control. The remaining lysate was pre-cleared by incubation with Protein A/G PLUS-Agarose beads (Santa Cruz, sc-2003) for 1 h at 4 °C. The pre-cleared lysates were then incubated overnight at 4 °C with specific antibodies against HAT1 (HUABIO, HA722332), FOXM1 (Proteintech, 13147-1-AP), or control IgG (Cell Signaling Technology, 2729s) together with Protein A/G PLUS-Agarose beads to form immune complexes. The beads were subsequently collected, washed thoroughly, and resuspended in sample buffer. Immunoprecipitated proteins were eluted by boiling at 100 °C for 10 min and analyzed by immunoblotting.

### Immunohistochemistry and immunofluorescence

Formalin-fixed paraffin-embedded tissues were sectioned at a thickness of 5 μm. The slides were baked at 65°C for 2 h, deparaffinized in xylene, and rehydrated through a graded ethanol series (100%, 90%, 80%, and 70%). Antigen retrieval was performed using Tris-EDTA buffer under microwave heating, maintaining a sub-boiling temperature for 20 min. The slides were then permeabilized with 0.1% Triton X-100 (Beyotime) and blocked with 10% bovine serum albumin (BSA). Subsequently, the sections were incubated with primary antibodies overnight at 4°C.

For immunohistochemistry, the slides were incubated with secondary antibodies (HRP polymer anti-rabbit IgG, ASP1613, Abcepta; HRP polymer anti-mouse IgG, ASP1615, Abcepta) at 37°C for 1 h. For immunofluorescence, the slides were incubated with fluorescent labeled secondary antibodies (Thermo Fisher) at 37°C for 1 h in the dark, followed by nuclear counterstaining with 4′,6-diamidino-2-phenylindole (DAPI; Beyotime, C1005). After washing three times with PBS, coverslips were mounted onto the glass slides. Immunofluorescence images were acquired using an Olympus SpinSR10 spinning disk confocal super-resolution microscope (Olympus, Japan). The primary antibodies used in this study included: anti-GLUT1 (Proteintech, 21829-1-AP; 1:400), anti-CD8a (Proteintech, 66868-1-Ig; 1:400), and anti-Ki67 (Abcam, ab16667; 1:200).

Protein expression was quantified by assessing both staining intensity and the proportion of positive cells. Staining intensity was graded on a scale of 0 to 3 (0 = negative, 1 = weak, 2 = moderate, 3 = strong). The percentage of positively stained cells was determined semi-quantitatively. A final histological score (H-score) was then calculated by multiplying the intensity grade by the percentage of positive cells. All assessments were performed independently by two observers to ensure consistency.

### Senescence-associated β-galactosidase staining

Cellular senescence was assessed using a Senescence-associated β-Galactosidase (SA-β-Gal) Staining Kit (Beyotime, China) following the manufacturer’s protocol. Briefly, cells were fixed and subsequently incubated overnight at 37 °C in the dark with the staining solution containing 0.05 mg/mL X-gal. After incubation, the cells were washed twice with PBS and imaged under a light microscope.

### Measurement of Acetyl Coenzyme A (Acetyl-CoA) levels

Following the manufacturer’s protocol, Acetyl-CoA was quantified in glucose-treated TE-1, KYSE140, and HNM007 cells using an ELISA kit (Elabscience, E-EL-0125). Standards and samples were added to the microtiter plate and incubated at 37°C for 90 min. After aspirating the supernatant, 100 µL of diluted biotinylated antibody was supplemented, followed by incubation at 37°C for 60 min. The plate was then washed three times, then incubated with 100 µL of enzyme conjugate working solution for 30 min prior to five additional washes. Afterwards, 90 µL of substrate solution was added and incubated for 15 min, and the reaction was terminated by adding 50 µL of stop solution. The absorbance value at 450 nm was measured using a microplate reader. Acetyl-CoA content was calculated according to the standard curve, and the differences between the control and experimental groups were further analyzed.

### Chromatin immunoprecipitation (ChIP)

Chromatin immunoprecipitation (ChIP) was carried out as previously described ^31^. Briefly, cells were harvested and cross-linked with 4% formaldehyde for 10 min at room temperature followed by quenching with 125 mM glycine for 5 min. Washed cells were resuspended in lysis buffer (1% SDS, 5 mM EDTA, and 50 mM Tris-HCl, pH 8.1) supplemented with protease inhibitors and then sonicated on ice for 15 min to shear cross-linked DNA to a length of approximately 300 bp. For immunoprecipitation, sheared chromatin was incubated with anti-Histone H3 (tri methyl K4) (Abcam, ab8580), anti-Histone H3 (acetyl K27) (Abcam, ab4729), and anti-Histone H3 (mono methyl K4) (Abcam, ab8895) antibodies overnight at 4 °C, and chromatin immune complexes were conjugated to Dynabeads Protein A/G. After extensive washing, the immunoprecipitated DNA was eluted, reverse cross-linked, and purified using a Qiagen PCR Purification Kit for subsequent sequencing. Sequencing reads were aligned to the reference genome with Bowtie2, and MACS3 (Model-based Analysis of ChIP-Seq) was employed to identify statistically significant enrichment peaks by comparing the ChIP sample against a control input sample.

### Assay for Transposase Accessible Chromatin (ATAC)-sequencing and analysis

ATAC-seq was performed using the High-Sensitivity Open Chromatin Profile Kit 2.0 (for Illumina®; Novoprotein, Cat# N248) following the manufacturer’s protocol. Briefly, 5×10^4^ control (Scramble), *HAT1*-knockdown (sh*HAT1*), and *FOXM1*-knockdown (sh*FOXM1*) TE-1 cells were collected, washed three times in cold PBS, and lysed for 10 min at 4 °C in lysis buffer. Freshly isolated nuclei were incubated with Tn5 transposase reaction mix at 37 °C for 30 min to fragment chromatin and integrate sequencing adapters into accessible genomic regions. Upon reaction stop, tagmented DNA was purified using tagment DNA extract beads. The purified fragments were amplified via PCR with indexed primers to generate sequencing libraries.

The resulting library fragments were subsequently evaluated using agarose gel electrophoresis and then sequenced on the GenoLab M platform (GeneMind, Shenzhen, China) in single-end (SE) mode. For quality control, fastp (v0.23.0) was applied to process the raw sequencing data following standard next-generation sequencing (NGS) quality control procedures. The clean single-end reads were aligned to the human reference genome (hg38) using BWA (v0.7.17). PCR duplicates were removed with Picard (v2.25.5), and reads mapping to the mitochondrial genome were excluded from downstream analysis. Peak calling was performed using MACS2 with the parameters --SPMR --nomodel --extsize 200 --q 0.01. The resulting peak calls were further filtered against the ENCODE blacklist for the hg38 genome to remove artifactual signals. BigWig (bw) files for data visualization were generated using bamCoverage from deepTools (v3.5.3). Heatmaps were also plotted using deepTools to visualize signal enrichment patterns. Peak annotation was conducted using ChIPSeeker (v1.38.0). Potential differentially accessible regions (DARs) were identified using MAnorm2. All sequencing tracks were visualized with the Integrative Genomics Viewer (IGV v2.3.61).

### Cleavage Under Targets and Tagmentation assay (CUT&Tag)

CUT&Tag was performed using the NovoNGS® CUT&Tag® 4.0 High-Sensitivity Kit (Novoprotein, N259-YH01) in accordance with the manufacturer’s protocol. Briefly, harvested cells were washed and fixed with 1% formaldehyde for 10 min at room temperature, followed by quenching with 125 mM glycine. After thorough washing, cells were captured using activated Concanavalin A-coated magnetic beads, permeabilized, and incubated overnight at 4 °C with primary antibodies targeting: anti-HAT1 (Cell Signaling Technology, 41490T) and anti-FOXM1 (Cell Signaling Technology, 20459S). Subsequently, samples were incubated with secondary antibody-conjugated pA-Tn5 transposase complex for 1 h at room temperature. Upon further washing, tagmentation was initiated by adding MgCl₂ and proceeded at 37 °C for 1 h. The reaction was terminated by adding stop buffer containing Proteinase K, followed by incubation at 55 °C for 2 h to reverse cross-links and elute DNA fragments. DNA was purified using tagmented DNA extraction beads, and libraries were constructed via PCR amplification. Final libraries were quantified and submitted for high-throughput sequencing.

### T-cell cytotoxicity assays

HNM007 cells were transfected with either a non-targeting control vector (WT) or *GLUT1*-KO constructs. For rescue experiments in *GLUT1*-KO cells, exogenous *GLUT1*, *HAT1*, or *FOXM1* was separately overexpressed. All cells were stimulated with 10 ng/mL IFNγ for 24 h. Afterwards, HNM007 cells were pulsed with 1 ng/mL OVA_257-264_ peptide for 2 h and seeded into 24-well plates. Pre-activated OT-I CD8⁺ T cells were co-cultured with the tumor cells at different effector-to-target (E:T) ratios (5:1 and 10:1). After 48 h of co-culture, both HNM007 cells and CD8⁺ T cells were harvested for subsequent cytotoxicity assessment. Viable HNM007 cells were quantified via crystal violet staining with absorbance measured at OD 570 nm using a microplate reader. The cytotoxic function of CD8⁺ T cells was assessed by flow cytometry to detect cells positive for granzyme B (GZMB; BioLegend, 372212) and IFNγ (BioLegend, 505823). A detailed schematic of the experimental setup is provided in the main figures and corresponding legends.

### Flow cytometry

Single-cell suspensions were isolated from murine HNM007 allograft tumors by mechanical grinding, followed by enzymatic digestion using 5 mg/mL collagenase IV (Invitrogen) and 0.25% Trypsin-EDTA (Gibco), supplemented with 10 U/mL DNase I (Sigma-Aldrich). Before staining, immune cells were pre-blocked with Fc Block™ (anti-mouse CD16/32) antibody (BD Biosciences). Live cells were labeled with Fixable Viability Stain 450 (562247, BD Biosciences) and subsequently stained on ice for 20 min with APC-conjugated CD45 (103112, BioLegend) and FITC-conjugated CD8 (100706, BioLegend) antibodies. For intracellular staining of GZMB and IFNγ, cells were fixed and permeabilized with the True-Nuclear™ Transcription Factor Buffer Set (424401, BioLegend), and further incubated with PerCP/Cyanine5.5-conjugated GZMB (372212, BioLegend) and AF700-conjugated IFNγ (505823, BioLegend) antibodies. In the T-cell cytotoxicity assays, CD8⁺ T cells were isolated and stained with PerCP/Cyanine5.5-conjugated GZMB antibody. Data were acquired on a CytoFLEX instrument (Beckman, USA) and analyzed using FlowJo v10.8.1 software.

### scRNA-seq and data analysis

Mouse tumors were washed with ice-cold PBS and cut into small pieces, which were then subjected to enzymatic digestion with 5 mg/mL collagenase IV (Invitrogen) and 0.25% Trypsin-EDTA (Gibco), supplemented with 10 U/mL DNase I (Sigma-Aldrich). Cells were filtered through a 100 μm filter and washed with PBS. To control the final number of cells captured, approximately 30,000 cells per sample were loaded. Single-cell libraries were constructed using the DNBelab C Series Single Cell Library Preparation Kit (MGI Tech) following the manufacturer’s protocol, and sequencing was performed on the BGISEQ-2000 platform.

Raw sequencing data were processed using the DNBC4tools pipeline (v2.1.3) to generate a gene expression count matrix for each cell per sample. The gene expression matrices of all samples were imported into Seurat (v4.3.0) and merged for subsequent analyses. To filter out low-quality cells, we set the following quality control criteria: (1) cells with number of detected genes per cell falling outside the range of 500 to 6,000 were excluded; (2) cells exhibiting a mitochondrial gene content exceeding 10% were removed; and (3) only genes detected in at least 5 cells were retained for downstream analysis. After quality control, data were normalized, and the top 2,000 highly variable genes were identified using the FindVariableFeatures function. The data were then scaled using the ScaleData function. To minimize batch effects across all samples, principal component analysis (PCA) was first performed, followed by the Harmony algorithm based on the initial PCA results to minimize batch effects ^55^. The resulting harmonized components were then applied to cell clustering and UMAP visualization. Cell types were annotated manually using conventional markers as reported previously. Cell-cell communication analysis was performed using CellChat (v1.6.1) to compare the signaling networks between the Control (Scr) and *Glut1*-knockdown (sh*Glut1*) groups. After merging the CellChat objects, we calculated the total number of interactions and interaction strength among cell types. Dot plots were generated using CellChat to visualize differential signaling patterns and predicted ligand-receptor pairs. Cell-cell communication networks centered on epithelial cells were established to visualize interactions with immune or stromal populations, with line thickness representing the intensity of intercellular crosstalk. Net analysis was performed to identify sender and receiver cell types involved in key immune-related signaling axes, such as the H2-K1-Cd8a and H2-K1-Cd8b1.

### Analysis of scRNA-seq from human ESCC samples

Publicly available ESCC single-cell RNA-seq datasets, including HRA003312, HRA004396, and HRA004740, were included in this analysis. We acquired raw sequencing data from 42 samples, comprising 35 ESCC tumors, 7 adjacent normal tissue samples from HRA003312 and HRA004396 datasets, while for the HRA004740 dataset, we acquired the corresponding expression matrix in 10x MEX format from OMIX005710. Raw reads were aligned to the GRCh38 reference genome, and gene expression was quantified using STARsolo (2.7.10b). The Seurat pipeline (v4.3.0) ^55^ was applied to each sample for downstream integrated analyses based on R (v4.3.0). To get high-quality data, low-quality cells were filtered out using the following criteria: (1) cells with fewer than 500 or more than 6,000 detected genes were excluded; (2) cells with mitochondrial gene content exceeding 20% were removed. Additionally, only genes detected in at least five cells were retained for downstream analysis. Gene expression and clustering results were visualized on a UMAP plot using RunUMAP. We next binned ESCC tumors into *GLUT1*-high and *GLUT1*-low groups based on the median expression level of *GLUT1* across samples, and compared their T cell subtypes distribution.

Differentially expressed genes (DEGs) were identified using a pseudo-bulk approach. Based on the clinical response information and classification criteria reported by Liu et al.,^28^ patients were categorized into the Complete Response (CR, N = 9) and Non-Complete Response (NCR, N = 23) groups. For tumor epithelial cells, sample-level raw count profiles were generated using AggregateExpression from Seurat with slot = “counts”. Differential expression analysis between the Complete Response (CR) and Non-Complete Response (NCR) groups was performed using DESeq2 (v1.42.1). To compare tumor and normal tissues, the same aggregation strategy was applied to all epithelial cells to generate pseudo-bulk count matrices for group-wise comparisons. In both analyses, genes with raw counts ≥ 10 in at least three samples were retained. DEGs were defined as genes with *P* value < 0.05 and an absolute log_2_ fold change >1.

For correlation analysis, the proportion of CD8⁺ T cells was calculated for each patient as the ratio of CD8⁺ T cells to total cells. Average expression levels of target genes in epithelial cells were obtained using AggregateExpression with slot = “data”, which sums exponentiated expression values to represent expression in non-log space. Spearman’s rank correlation analysis was used to evaluate associations between gene expression and CD8⁺ T cells proportions, and the results were visualized using ggplot2 (v4.0.2).

### Single-cell ATAC sequencing and data analysis

Single-cell ATAC-seq (scATAC-seq) was performed following an adapted single-nucleus protocol ^56^. Briefly, cells were lysed in 100 µL of lysis buffer A (10 mM Tris-HCl pH 7.5, 10 mM NaCl, 3 mM MgCl₂, 0.1% NP40, 0.1% Tween-20, 0.01% Digitonin), and the lysis was quenched with 1 mL of PBS containing 1% BSA. Nuclei were pelleted by centrifugation (400 × g, 5 min, 4 °C), washed twice with PBS, and resuspended in PBS with 1% BSA. After trypan blue staining for quantification, scATAC-seq libraries were prepared using the DNBelab C Series Single-Cell ATAC Library Prep Set (MGI, 1000021878) and sequenced on the DNBSEQ-T7 platform.

Reference genomes were downloaded from the UCSC database (mm10 for mouse). All mitochondrial DNA regions in both reference genomes were masked. Raw sequencing data were aligned using chromap (v0.2.3) with the parameters “--preset atac --bc-error-threshold 0 --trim-adapters -x”. Only reads mapped to the nuclear genome were retained. Cell barcode assignment was performed using d2c (v1.4.4).

To ensure high-quality chromatin accessibility profiles, cells were filtered based on the following criteria using the ArchR (v1.0.2) ^57^: (1) a Transcription Start Site (TSS) enrichment score ≥ 2; (2) a minimum of 500 unique nuclear fragments (minFrags = 500). Following initial quality control, doublets were identified and filtered using ‘addDoubletScores’ and ‘filterDoublets’ functions, with a filter ratio of 1. Dimensionality reduction was performed using Iterative Latent Semantic Indexing (LSI) via the addIterativeLSI function. To minimize the batch effects across samples, we applied Harmony integration using the addHarmony function. Initial clustering was conducted using addClusters, and data were visualized via Uniform Manifold Approximation and Projection (UMAP). To enable high-resolution analysis of the TME, particularly the dynamic shifts in CD8⁺ T cell populations, we subset the dataset to exclude the predominant epithelial cell fraction. For this non-epithelial subset, UMAP and density distribution patterns were recalculated to compare the Scr and sh*Glut1* groups, and changes in cell type proportions were quantified using stacked bar plots. Marker genes were identified via the getMarkerFeatures function, with the Gene Activity Score used as a surrogate for gene expression. Finally, to visualize group-specific chromatin accessibility, aggregate browser tracks were generated for key loci, including *Hat1*, *Foxm1*, *Ccl5*, *Gzmb*, and *Ifng*.

### TCGA data analysis

mRNA expression profiles and corresponding clinical information for ESCC patients were obtained from The Cancer Genome Atlas (TCGA) through the Genomic Data Commons (GDC) portal. Samples were restricted to patients with TxN0M0 staging, and cases with documented neoadjuvant chemotherapy were excluded based on pathological annotations. Glycolysis activity was evaluated using Gene Set Variation Analysis (GSVA)/ssGSEA based on the KEGG “Glycolysis/Gluconeogenesis” gene set (hsa00010). Patients were stratified into high- and low-glucose metabolism groups according to the median glycolysis scores, including high-glucose metabolism (N = 34) and low-glucose metabolism (N = 35) groups for cumulative incidence function (CIF) analysis. Recurrence-free survival (RFS) was analyzed using Kaplan-Meier curves and compared by log-rank tests ^43^.

In addition, TCGA transcriptomic datasets from ESCC, HNSC, and LUSC cohorts were analyzed to compare the expression patterns of the 14 GLUT family members (*GLUT1*-*GLUT14*) between tumor and normal tissues.

## Quantification and statistical analysis

Statistical analyses were performed with GraphPad Prism (v9.0) and R (v4.0.0). Unless stated otherwise, quantitative data are presented as mean ± SEM. For box-and-whisker plots, the center line indicates the median, the box indicates the interquartile range, and the whiskers indicate the minimum and maximum values. Violin plots show the distribution of individual values, with the solid line indicating the median and the dashed lines indicating the first and third quartiles. The data presentation, sample size, and statistical test used for each analysis are specified in the corresponding figure legend. Statistical significance was defined as a two-sided *P* value < 0.05. Differences between two groups were evaluated by Student’s t-test, while one- or two-way ANOVA was employed for multi-group comparisons. For survival analyses, Kaplan-Meier curves were plotted, and log-rank tests were used to calculate *P* values between groups. The hazard ratio (HR) along with a 95% confidence interval (CI) was estimated using Cox proportional hazards regression. Details of the statistical tests are provided in the figure legends.

## Materials availability

This study did not generate new unique reagents.

## Data and code availability

All materials and reagents used in this study are listed in the Key Resources Table (Supplementary Table 2). The raw sequencing data generated in this study have been deposited in the Genome Sequence Archive (GSA) of the China National Center for Bioinformation and are publicly accessible at https://ngdc.cncb.ac.cn/gsa. These datasets include ChIP-seq profiles of H3K27ac, H3K4me3, and H3K4me1 in TE-1 cells; ATAC-seq data from *HAT1* knockdown and *FOXM1* knockdown TE-1 cells; and CUT&Tag data for HAT1 and FOXM1 in TE-1 cells, under accession number HRA017636. In addition, scRNA-seq and snATAC-seq data from mouse tumor models (Scr, sh*Glut1*-1, and sh*Glut1*-2) have been deposited under accession number CRA041703. Previously published scRNA-seq datasets from ESCC patients are available under accession numbers HRA003312, HRA004396 and HRA004740.

## Acknowledgments

We acknowledge the animal core facility at Hefei Institutes of Physical Science (HFIPS), Chinese Academy of Sciences (CAS). We thank Dr. Xin Zhang (CAS Key Laboratory of High Magnetic Field and Ion Beam Physical Biology, HFIPS, CAS) for offering the confocal microscope imaging system. We thank Dr. Boshi Wang (Shanghai Cancer Institute, Renji Hospital, Shanghai Jiao Tong University School of Medicine) for sharing the TRIPZ vector. This work was supported by grants from the Strategic Priority Research Program of the Chinese Academy of Sciences (XDC0200000), the National Natural Science Foundation of China (82572978, 82273010, 82302896, 82503236, 82504042), the Anhui Provincial Natural Science Foundation (2408085J044, 2508085QH287), and the HFIPS Director’s Fund of Hefei Institutes of Physical Science, Chinese Academy of Sciences (BJPY2024B06, YZJJQY202403).

## Author contributions

Y.-Y.J. and Y.J. conceived the study. J.-X.D., J.Z., J.H., S.K., and K.M. performed experiments and analyzed data. C.Y., D.-D.W., F.W., J.M., and J.F. assisted with experiments and validation. Y.-W.Z., W.-Q.W., and H.P. contributed to data analysis and interpretation. Y.-Y.J., Y.J., and M.W. supervised the study. J.-X.D., J.Z., and J.H. wrote the original draft. Y.J., Y.-Y.J., K.M., and M.W. reviewed and edited the manuscript. All authors approved the final version of the manuscript.

## References

1. Siegel, R.L., Miller, K.D., Wagle, N.S. & Jemal, A. Cancer statistics, 2023. CA Cancer J Clin 73, 17–48 (2023).

2. Chang, J. et al. Single-cell multi-stage spatial evolutional map of esophageal carcinogenesis. Cancer Cell 43, 380–397.e7 (2025).

3. Forde, P.M. et al. Neoadjuvant PD-1 Blockade in Resectable Lung Cancer. N Engl J Med 378, 1976–1986 (2018).

4. Menzies, A.M. et al. Pathological response and survival with neoadjuvant therapy in melanoma: a pooled analysis from the International Neoadjuvant Melanoma Consortium (INMC). Nat Med 27, 301–309 (2021).

5. Doki, Y. et al. Nivolumab Combination Therapy in Advanced Esophageal Squamous-Cell Carcinoma. N Engl J Med 386, 449–462 (2022).

6. Xu, J. et al. Tislelizumab plus chemotherapy versus placebo plus chemotherapy as first-line treatment for advanced or metastatic oesophageal squamous cell carcinoma (RATIONALE-306): a global, randomised, placebo-controlled, phase 3 study. Lancet Oncol 24, 483–495 (2023).

7. Yin, J. et al. Neoadjuvant adebrelimab in locally advanced resectable esophageal squamous cell carcinoma: a phase 1b trial. Nat Med 29, 2068–2078 (2023).

8. Verschoor, Y.L. et al. Neoadjuvant atezolizumab plus chemotherapy in gastric and gastroesophageal junction adenocarcinoma: the phase 2 PANDA trial. Nat Med 30, 519–530 (2024).

9. Wei, D.D. et al. Perioperative immunotherapy for esophageal squamous cell carcinoma. Front Immunol 15, 1330785 (2024).

10. Di Micco, R., Krizhanovsky, V., Baker, D. & d’Adda di Fagagna, F. Cellular senescence in ageing: from mechanisms to therapeutic opportunities. Nat Rev Mol Cell Biol 22, 75–95 (2021).

11. Hanahan, D. & Weinberg, R.A. Hallmarks of cancer: the next generation. Cell 144, 646–74 (2011).

12. Shahbandi, A. et al. Breast cancer cells survive chemotherapy by activating targetable immune-modulatory programs characterized by PD-L1 or CD80. Nat Cancer 3, 1513–1533 (2022).

13. Pei, S. et al. Age-related decline in CD8(+) tissue resident memory T cells compromises antitumor immunity. Nat Aging 4, 1828–1844 (2024).

14. Ma, F. et al. Tumor extracellular vesicle-derived PD-L1 promotes T cell senescence through lipid metabolism reprogramming. Sci Transl Med 17, eadm7269 (2025).

15. Chen, H.A. et al. Senescence Rewires Microenvironment Sensing to Facilitate Antitumor Immunity. Cancer Discov 13, 432–453 (2023).

16. Marin, I. et al. Cellular Senescence Is Immunogenic and Promotes Antitumor Immunity. Cancer Discov 13, 410–431 (2023).

17. Sturmlechner, I. et al. p21 produces a bioactive secretome that places stressed cells under immunosurveillance. Science 374, eabb3420 (2021).

18. Wang, C. et al. Inducing and exploiting vulnerabilities for the treatment of liver cancer. Nature 574, 268–272 (2019).

19. Prieto, L.I., Sturmlechner, I., Goronzy, J.J. & Baker, D.J. Senescent cells as thermostats of antitumor immunity. Sci Transl Med 15, eadg7291 (2023).

20. Zhou, P. et al. Targeting senescent EGR1(+) B cells enhances immunotherapy efficacy in esophageal squamous cell carcinoma. Cell Rep Med 7, 102532 (2026).

21. Deng, D. et al. Crystal structure of the human glucose transporter GLUT1. Nature 510, 121–125 (2014).

22. Zhang, Z. et al. DHHC9-mediated GLUT1 S-palmitoylation promotes glioblastoma glycolysis and tumorigenesis. Nat Commun 12, 5872 (2021).

23. De Leo, A. et al. Glucose-driven histone lactylation promotes the immunosuppressive activity of monocyte-derived macrophages in glioblastoma. Immunity 57, 1105–1123.e8 (2024).

24. Guo, D. et al. Aerobic glycolysis promotes tumor immune evasion by hexokinase2-mediated phosphorylation of IκBα. Cell Metab 34, 1312–1324.e6 (2022).

25. Cascone, T. et al. Increased Tumor Glycolysis Characterizes Immune Resistance to Adoptive T Cell Therapy. Cell Metab 27, 977–987.e4 (2018).

26. Cotton, S. et al. Target Score-A Proteomics Data Selection Tool Applied to Esophageal Cancer Identifies GLUT1-Sialyl Tn Glycoforms as Biomarkers of Cancer Aggressiveness. Int J Mol Sci 22, 1664 (2021).

27. Minakata, N. et al. Immunohistochemistry and oxygen saturation endoscopic imaging reveal hypoxia in submucosal invasive esophageal squamous cell carcinoma. Cancer Med 12, 15809–15819 (2023).

28. Liu, Z. et al. Progenitor-like exhausted SPRY1(+)CD8(+) T cells potentiate responsiveness to neoadjuvant PD-1 blockade in esophageal squamous cell carcinoma. Cancer Cell 41, 1852–1870.e9 (2023).

29. Ji, G. et al. Single-cell profiling of response to neoadjuvant chemo-immunotherapy in surgically resectable esophageal squamous cell carcinoma. Genome Med 16, 49 (2024).

30. Wang, C. et al. Single-cell profiling identifies biomarkers for immunochemotherapy in esophageal squamous cell carcinoma. Cancer Lett 633, 217988 (2025).

31. Jiang, Y.Y. et al. TP63, SOX2, and KLF5 Establish a Core Regulatory Circuitry That Controls Epigenetic and Transcription Patterns in Esophageal Squamous Cell Carcinoma Cell Lines. Gastroenterology 159, 1311–1327.e19 (2020).

32. Jiang, Y.Y. et al. Targeting super-enhancer-associated oncogenes in oesophageal squamous cell carcinoma. Gut 66, 1358–1368 (2017).

33. Hsieh, W.C. et al. Glucose starvation induces a switch in the histone acetylome for activation of gluconeogenic and fat metabolism genes. Mol Cell 82, 60–74.e5 (2022).

34. Gruber, J.J. et al. HAT1 Coordinates Histone Production and Acetylation via H4 Promoter Binding. Mol Cell 75, 711–724.e5 (2019).

35. Ma, Y. et al. The Nuclear Localization of ACLY Guards Early Embryo Development Through Recruiting P300 and HAT1 to Promote Histone Acetylation and Transcription. Adv Sci (Weinh) 12, e14367 (2025).

36. Ferreira, F.J. et al. FOXM1 expression reverts aging chromatin profiles through repression of the senescence-associated pioneer factor AP-1. Nat Commun 16, 2931 (2025).

37. Xie, F. et al. Targeting FOXM1 condensates reduces breast tumour growth and metastasis. Nature 638, 1112–1121 (2025).

38. Lugassy, J. et al. Development of DPP-4-resistant CXCL9-Fc and CXCL10-Fc chemokines for effective cancer immunotherapy. Proc Natl Acad Sci U S A 122, e2501791122 (2025).

39. Mowat, C., Mosley, S.R., Namdar, A., Schiller, D. & Baker, K. Anti-tumor immunity in mismatch repair-deficient colorectal cancers requires type I IFN-driven CCL5 and CXCL10. J Exp Med 218, e20210108 (2021).

40. House, I.G. et al. Macrophage-Derived CXCL9 and CXCL10 Are Required for Antitumor Immune Responses Following Immune Checkpoint Blockade. Clin Cancer Res 26, 487–504 (2020).

41. Tao, W., Yu, Z. & Han, J.J. Single-cell senescence identification reveals senescence heterogeneity, trajectory, and modulators. Cell Metab 36, 1126–1143.e5 (2024).

42. Lien, E.C. et al. Low glycaemic diets alter lipid metabolism to influence tumour growth. Nature 599, 302–307 (2021).

43. Wu, C.Y. et al. Glucose restriction shapes pre-metastatic innate immune landscapes in the lung through exosomal TRAIL. Cell 188, 5701–5716.e19 (2025).

44. Morrissey, S.M. et al. Tumor-derived exosomes drive immunosuppressive macrophages in a pre-metastatic niche through glycolytic dominant metabolic reprogramming. Cell Metab 33, 2040–2058.e10 (2021).

45. Verma, S. et al. Pharmacologic LDH inhibition redirects intratumoral glucose uptake and improves antitumor immunity in solid tumor models. J Clin Invest 134, e177606 (2024).

46. Guerrero, J.A. et al. GLUT1 overexpression in CAR-T cells induces metabolic reprogramming and enhances potency. Nat Commun 15, 8658 (2024).

47. Wang, L. et al. SLC2A1(+) tumour-associated macrophages spatially control CD8(+) T cell function and drive resistance to immunotherapy in non-small-cell lung cancer. Nat Cell Biol 28, 338–348 (2026).

48. Gonzalez-Menendez, P. et al. GLUT1 protects prostate cancer cells from glucose deprivation-induced oxidative stress. Redox Biol 17, 112–127 (2018).

49. Li, T. et al. Metabolism/Immunity Dual-Regulation Thermogels Potentiating Immunotherapy of Glioblastoma Through Lactate-Excretion Inhibition and PD-1/PD-L1 Blockade. Adv Sci (Weinh) 11, e2310163 (2024).

50. Mueckler, M. & Thorens, B. The SLC2 (GLUT) family of membrane transporters. Mol Aspects Med 34, 121–138 (2013).

51. Cairns, R.A., Harris, I.S. & Mak, T.W. Regulation of cancer cell metabolism. Nat Rev Cancer 11, 85–95 (2011).

52. Feng, T. et al. Fructose and glucose from sugary drinks enhance colorectal cancer metastasis via SORD. Nat Metab 7, 2018–2032 (2025).

53. Fowle-Grider, R. et al. Dietary fructose enhances tumour growth indirectly via interorgan lipid transfer. Nature 636, 737–744 (2024).

54. Jiang, Y. et al. Establishing mouse and human oral esophageal organoids to investigate the tumor immune response. Dis Model Mech 17, dmm05031 (2024).

55. Stuart, T. et al. Comprehensive Integration of Single-Cell Data. Cell 177, 1888–1902.e21 (2019).

56. Mazid, M.A. et al. Rolling back human pluripotent stem cells to an eight-cell embryo-like stage. Nature 605, 315–324 (2022).

57. Granja, J.M. et al. ArchR is a scalable software package for integrative single-cell chromatin accessibility analysis. Nat Genet 53, 403–411 (2021).

