## Supplementary Figure for "High glucose confers senescence resistance via GLUT1 epigenetic rewiring to blunt immunotherapy responses in esophageal squamous cell carcinoma"

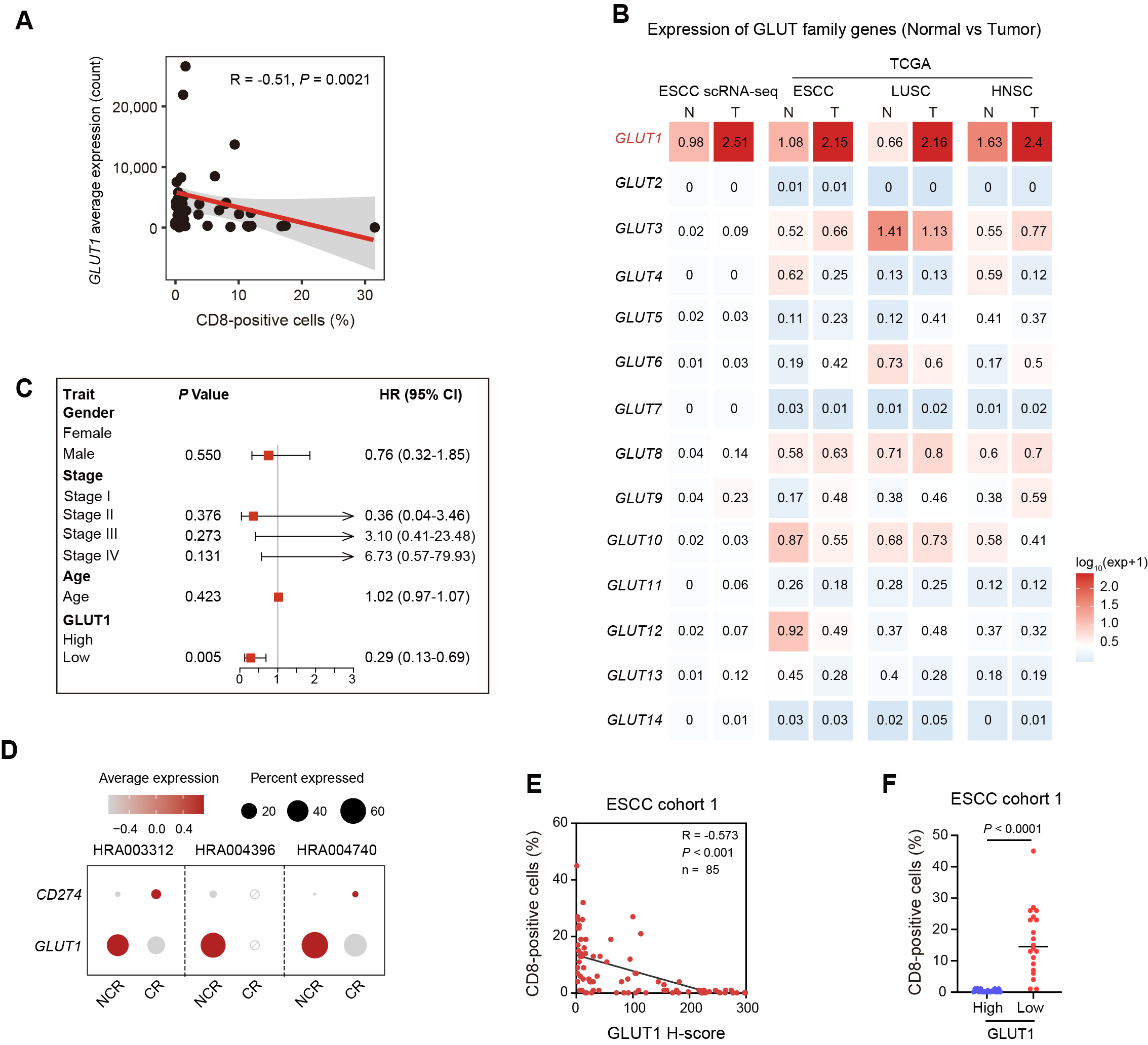

**Figure S1. GLUT1 correlates with poor immunotherapy response and unfavorable prognosis, related to Figure 1.**

(A) Correlation between CD8⁺ T cell ratio and GLUT1 average expression. Each point represents an individual sample. The red line indicates the linear regression fit, with the shaded area representing the 95% confidence interval. A modest but statistically significant negative correlation is observed (R = - 0.51, *P* = 0.0021).

(B) Heatmap showing the expression profiles of GLUT family genes ESCC tumor and normal tissues. scRNA-seq data from HRA003312, HRA004396, and HRA004740. Bulk RNA-seq data from The Cancer Genome Atlas (TCGA). ESCC: esophageal squamous cell carcinoma; HNSC: head and neck squamous cell carcinoma; LUSC: lung squamous cell carcinoma.

**(C)** Forest plot of multivariable Cox proportional hazards analysis evaluating the association of clinicopathological variables and **GLUT1** expression with survival. Hazard ratios (HRs), 95% confidence intervals (CIs), and P values are shown.

(D) Dot plot showing the average expression and percentage of cells expressing *GLUT1* and *CD274* (*PD-L1*) across three independent ESCC cohorts (HRA003312, HRA004396, and HRA004740) in complete response (CR) and non-complete response (NCR) groups. Dot color represents scaled average expression (blue to red), and dot size indicates the percentage of expressing cells. The circled slash symbol denotes the absence of the CR group in the HRA004396 dataset.

(E) Correlation analysis between GLUT1 H-score and CD8⁺ T cell percentage in the ESCC cohort. Spearman correlation coefficient and *P* value are shown.

(F) Comparison of CD8⁺ T cell infiltration between GLUT1 high and low tumors. Each dot represents an individual sample; horizontal lines indicate mean values. Statistical analysis was performed using Student’s test.

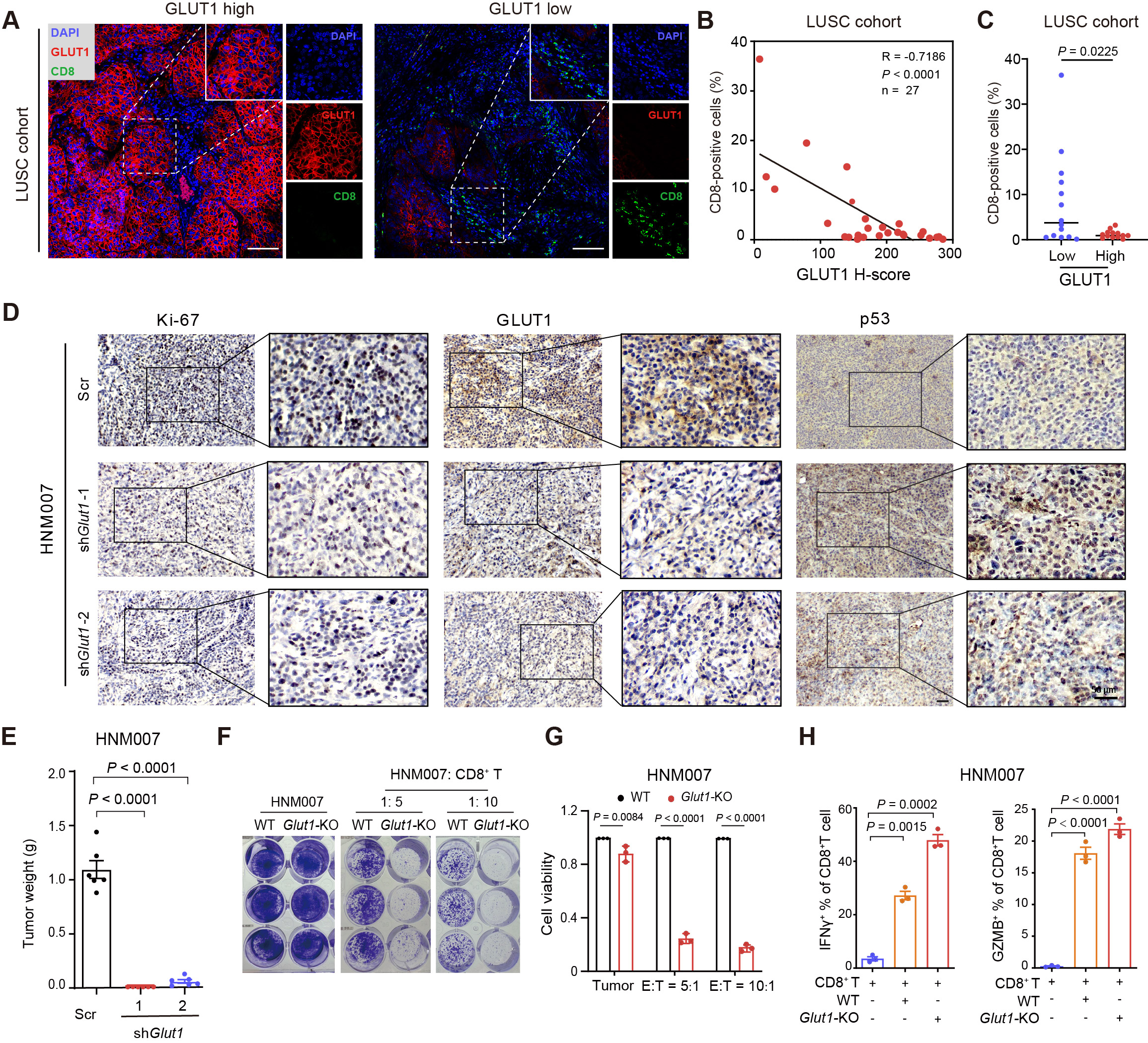

**Figure S2. GLUT1 suppresses antitumor immunity and promotes tumor growth *in vivo* and in clinical cohorts, related to Figure 2.**

(A) Representative IF images of GLUT1 (red), CD8⁺ T cells (green), and DAPI (blue) in LUSC samples with high or low GLUT1 expression.

(B) Correlation analysis between GLUT1 expression (H-score) and CD8⁺ T cell infiltration in (A), N = 27.

(C) Quantification of CD8⁺ T cell infiltration in (A), N = 27.

(D) Immunohistochemical (IHC) staining showing the protein levels of Ki-67, GLUT1, and p53 in Scr and sh*Glut*1 allografts. Representative images and enlarged views are shown. Scale bar, 50 μm.

(E) Comparison of tumor weights from HNM007 allografts expressing control (Scr) or two independent sh*Glut1* constructs in C57BL/6J mice.

(F) Crystal violet staining showing the viability of WT or *Glut1*-KO HNM007 cells co-cultured with CD8⁺ T cells at indicated effector-to-target (E: T) ratios (5:1 and 10:1).

(G) Quantification of tumor cell viability in (F).

(H) Flow cytometric analysis of IFNγ and GZMB production in co-cultures in (F).

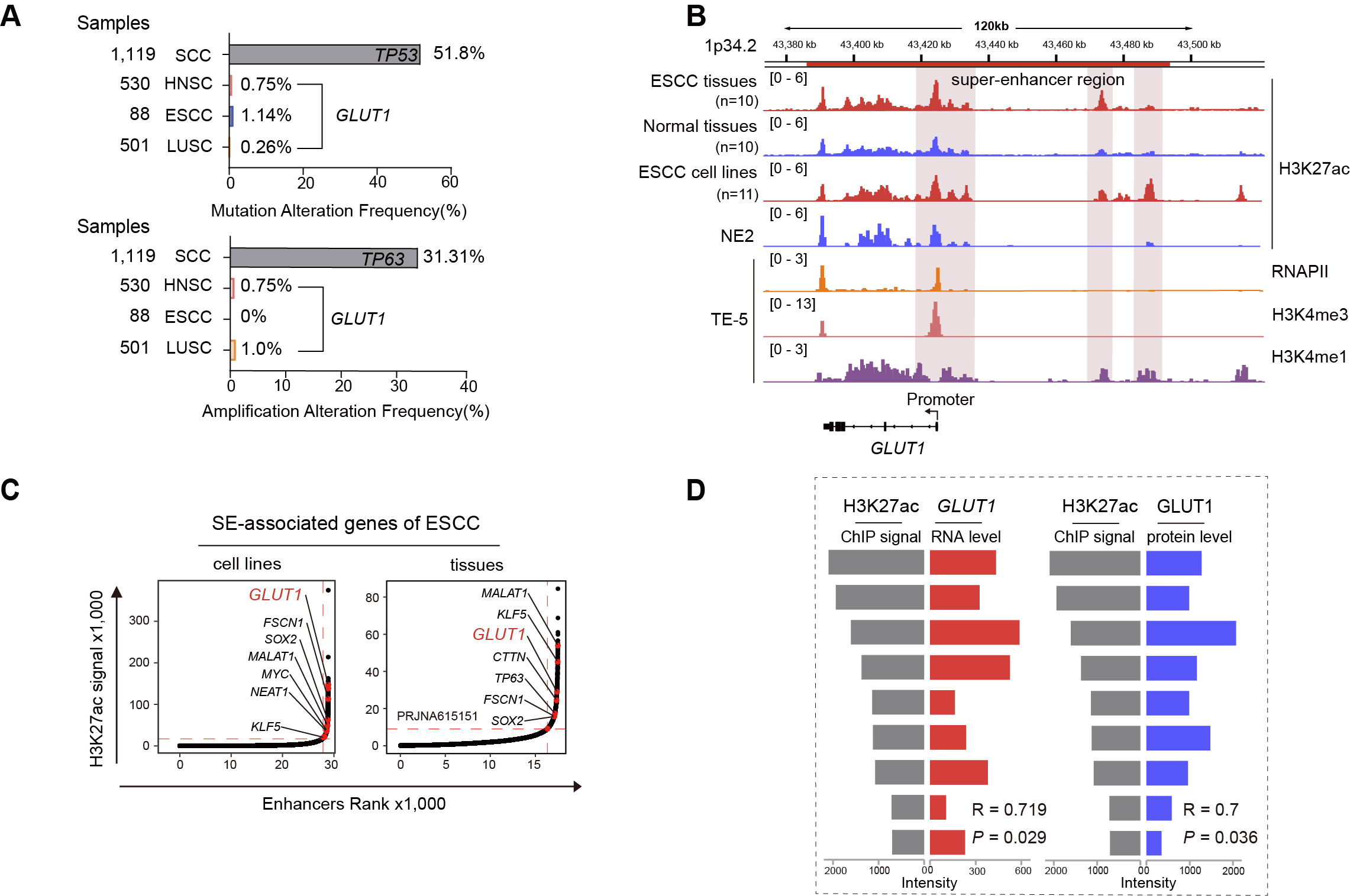

**Figure S3. Super-enhancer-associated activation of *GLUT1* in ESCC, related to Figure 3.**

(A) Frequency of genomic alterations across squamous cell carcinoma (SCC) cohorts, including HNSC, ESCC, and LUSC. Top panel shows mutation frequencies, highlighting TP53 as the most frequently mutated gene. Bottom panel shows amplification frequencies, with TP63 as a major amplified gene. *GLUT1* alteration frequencies are indicated for comparison across cancer types.

(B) Genomic landscape of the *GLUT1* locus. ChIP-seq profiles display H3K27ac signal across ESCC tissues, normal tissues, ESCC cell lines, and an immortalized esophageal epithelial cell (NE2), highlighting super-enhancer (SE) region upstream of *GLUT1*. Additional tracks show RNA polymerase II (RNAPII), H3K4me3, and H3K4me1 enrichment, indicating active transcriptional regulation. The promoter region is annotated.

(C) Identification of SE-associated genes in ESCC. Plots show enhancer ranking based on H3K27ac ChIP-seq signal intensity in ESCC cell lines (left) and tissues (right). Genes associated with high-ranking SEs are labeled.

(D) Correlation between H3K27ac signal density and *GLUT1* expression in mRNA (left) and protein level (right). Pearson correlation coefficients (R) and corresponding *P* values are indicated.

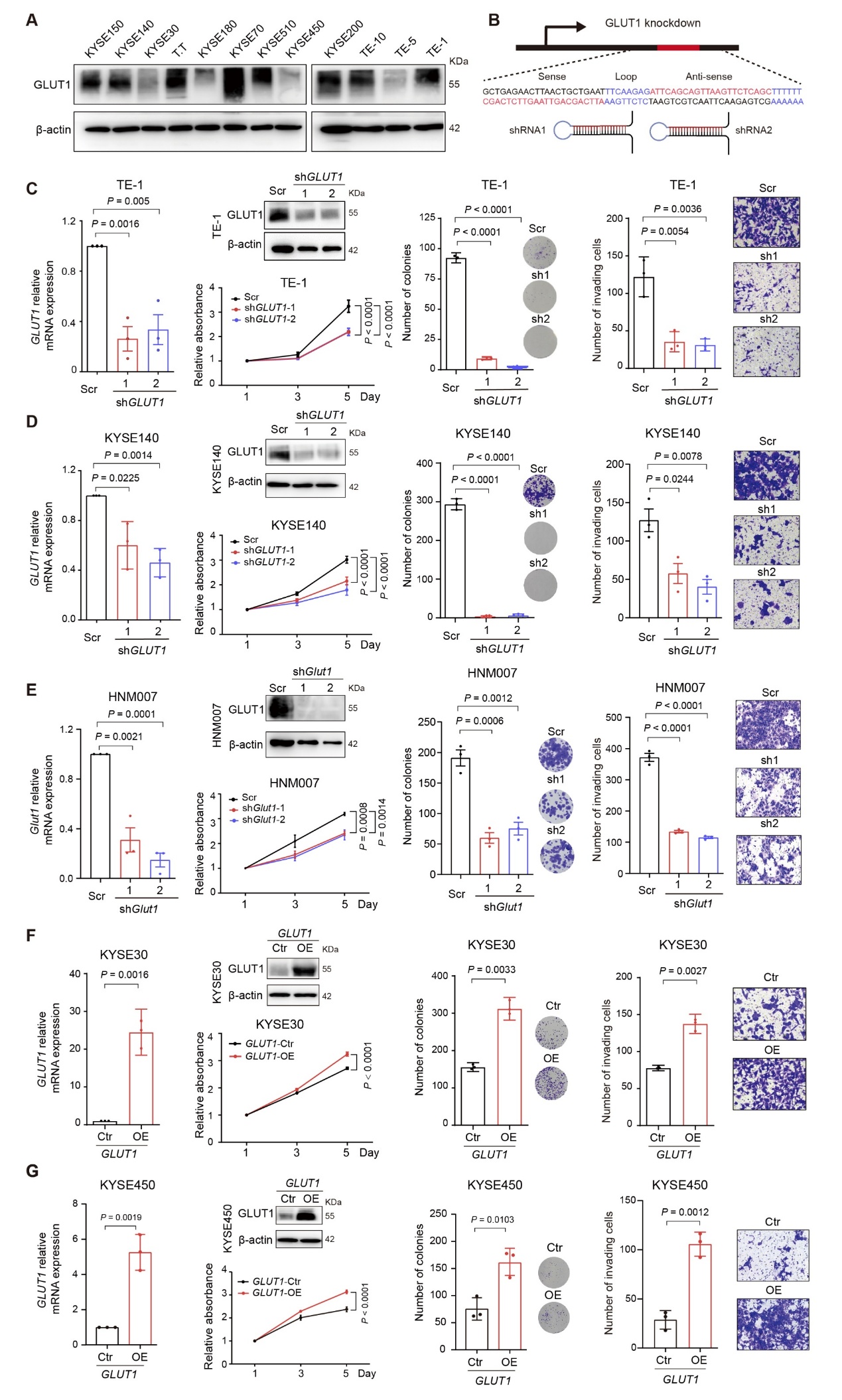

**Figure S4. GLUT1 promotes proliferation, clonogenicity, and invasion in ESCC, related to Figure 4.**

(A) Western blot analysis of GLUT1 protein expression in a panel of ESCC cell lines. β-actin was used as a loading control.

(B) Schematic illustration of the shRNA constructs targeting GLUT1. Two independent shRNAs (shRNA1 and shRNA2) targeting GLUT1 were designed.

(C and D) Knockdown of *GLUT1* suppresses malignant phenotypes in ESCC cells. In TE-1 (C) and KYSE140 (D) cells, GLUT1 silencing by two independent shRNAs was validated by RT-qPCR and western blotting. Cell proliferation was assessed by growth curve analysis, colony forming ability was evaluated by colony formation assays, and invasive capacity was measured by transwell invasion assays. Representative stained images and corresponding quantitative analyses are shown.

(E) Knockdown of *Glut1* inhibits malignant phenotypes in murine HNM007 cells. *Glut1* silencing efficiency was confirmed by RT-qPCR and western blotting. Cell proliferation, colony formation, and invasion were then analyzed as in panels C and D. Representative images and quantification are shown.

(F and G) Ectopic overexpression of GLUT1 enhances tumor cell growth and invasiveness in ESCC cells. In KYSE30 (F) and KYSE450 (G) cells, GLUT1 overexpression was confirmed by RT-qPCR and western blotting, followed by assessment of cell proliferation, colony formation, and transwell invasion assays. Representative images and quantitative results are shown. Data are presented as mean ± SEM of three independent experiments, and statistical comparisons were performed using student’s test and one-way ANOVA with Tukey’s multiple comparison test (C-G). Statistical significance is indicated in each panel by *P* values.

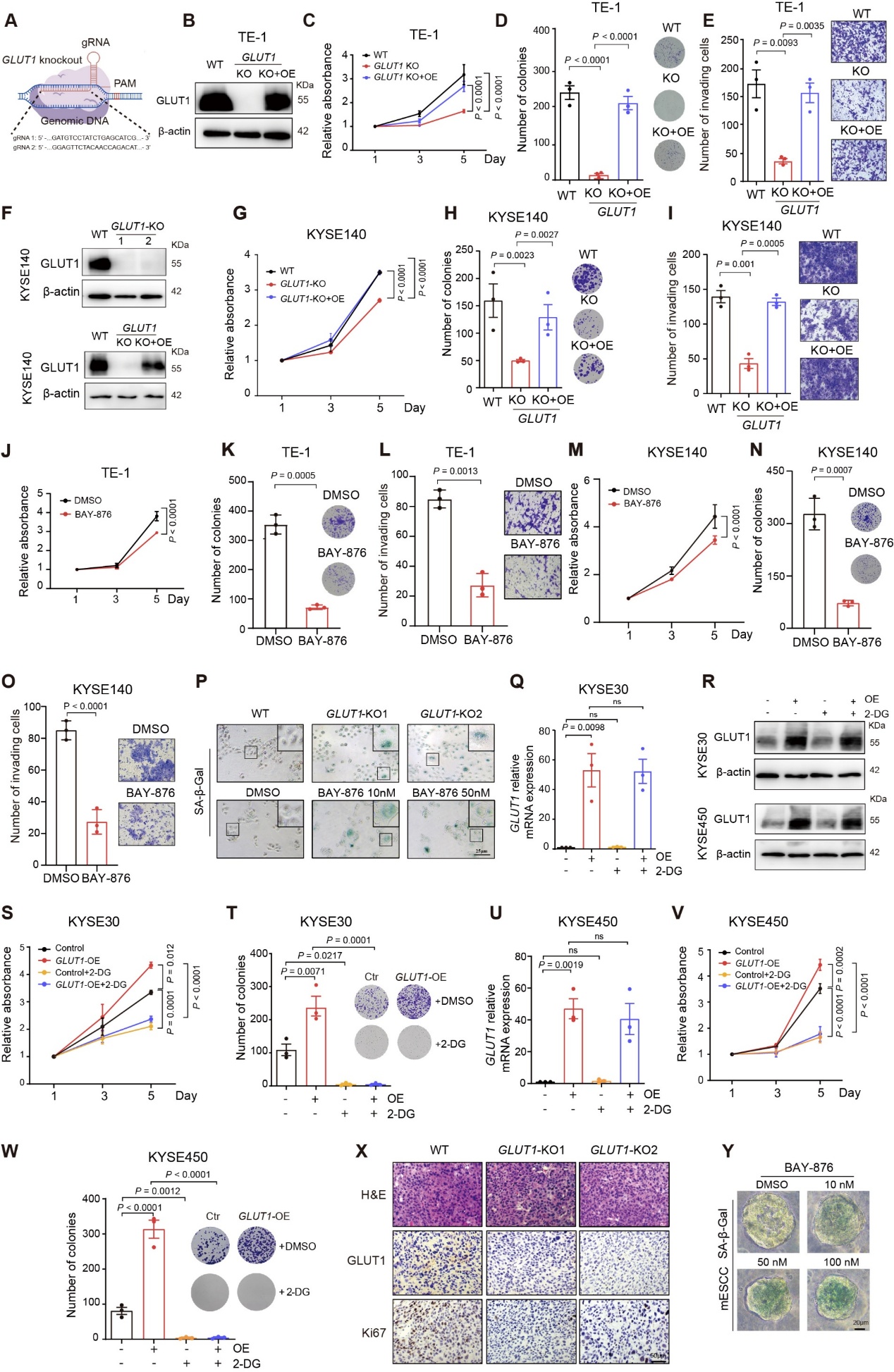

**Figure S5. Genetic and pharmacological inhibition of GLUT1 suppresses malignant phenotypes of ESCC, related to Figure 4.**
**(A)** Schematic illustration of the CRISPR/Cas9 strategy used to generate ***GLUT1***-KO cells. Two independent guide RNAs (gRNAs) targeting the ***GLUT1*** genomic locus adjacent to the protospacer adjacent motif (PAM) are shown.
**(B)** Western blot validation of ***GLUT1***-KO and re-expression in **TE-1** cells. **β-actin** was used as a loading control.
**(C-E)** Functional characterization of ***GLUT1*** depletion and rescue in **TE-1** cells. Cell proliferation was assessed by growth curve analysis (**C**), colony forming ability by colony formation assay (**D**), and invasive capacity by transwell invasion assay (**E**). Comparisons were made between Wild-type (WT), ***GLUT1*** knockout (KO), and ***GLUT1*** knockout with re-expression (KO+OE) cells. Representative images and quantitative analyses are shown.
**(F)** Western blot validation of ***GLUT1***-KO and rescue in **KYSE140** cells.
**(G-I)** Functional analyses of **GLUT1** depletion and rescue in **KYSE140** cells. Cell proliferation (**G**), colony formation (**H**), and invasion (**I**) were evaluated in WT, ***GLUT1***-KO, and ***GLUT1***-KO+OE cells. Representative images and corresponding quantification are shown.
**(J-P)** Pharmacological inhibition of GLUT1 with BAY-876 suppresses tumor cell growth (J and M), colony formation (K and N) and invasion (L and O) and induces cell senescence, as indicated by representative SA-β-Gal staining (P) in **TE-1** cells and **KYSE140** cells. Representative images and quantification are shown. Insets show enlarged views of the boxed regions (P). Scale bar, 25 μm.
**(Q-W)** The oncogenicity effect of **GLUT1** depends on glucose metabolism and is abrogated by 2-deoxy-D-glucose (2-DG). In **KYSE30** and **KYSE450** cells, ***GLUT1*** overexpression (OE) was confirmed by RT-qPCR (**Q**) and western blot (**R**). Cell proliferation assays (**S**, **V**) and colony formation assays (**T**, **W**) showing enhanced growth and clonogenicity **with *GLUT1*** overexpression, whereas the effects was markedly attenuated by treatment with 2-DG.
**(X)** Representative images of hematoxylin and eosin (H&E) and IHC staining for **GLUT1** and **Ki67** in xenograft tumors derived from WT, ***GLUT1***-KO1, and ***GLUT1***-KO2 KYSE140 cells. Scale bar, 50 μm.
**(Y)** Representative images of SA-β-Gal staining of mouse ESCC (mESCC) organoids treated with BAY-876 at the indicated concentrations. Scale bar, 20 μm.

Statistical significance is indicated by P values in each panel. “ns” indicates no significant difference. Data are presented as mean ± SEM of three independent experiments, and statistical comparisons were performed using student’s test (K, L, N, and O) and one-way ANOVA with Tukey’s multiple comparison test (C-E, G-J, M, Q, S-W).

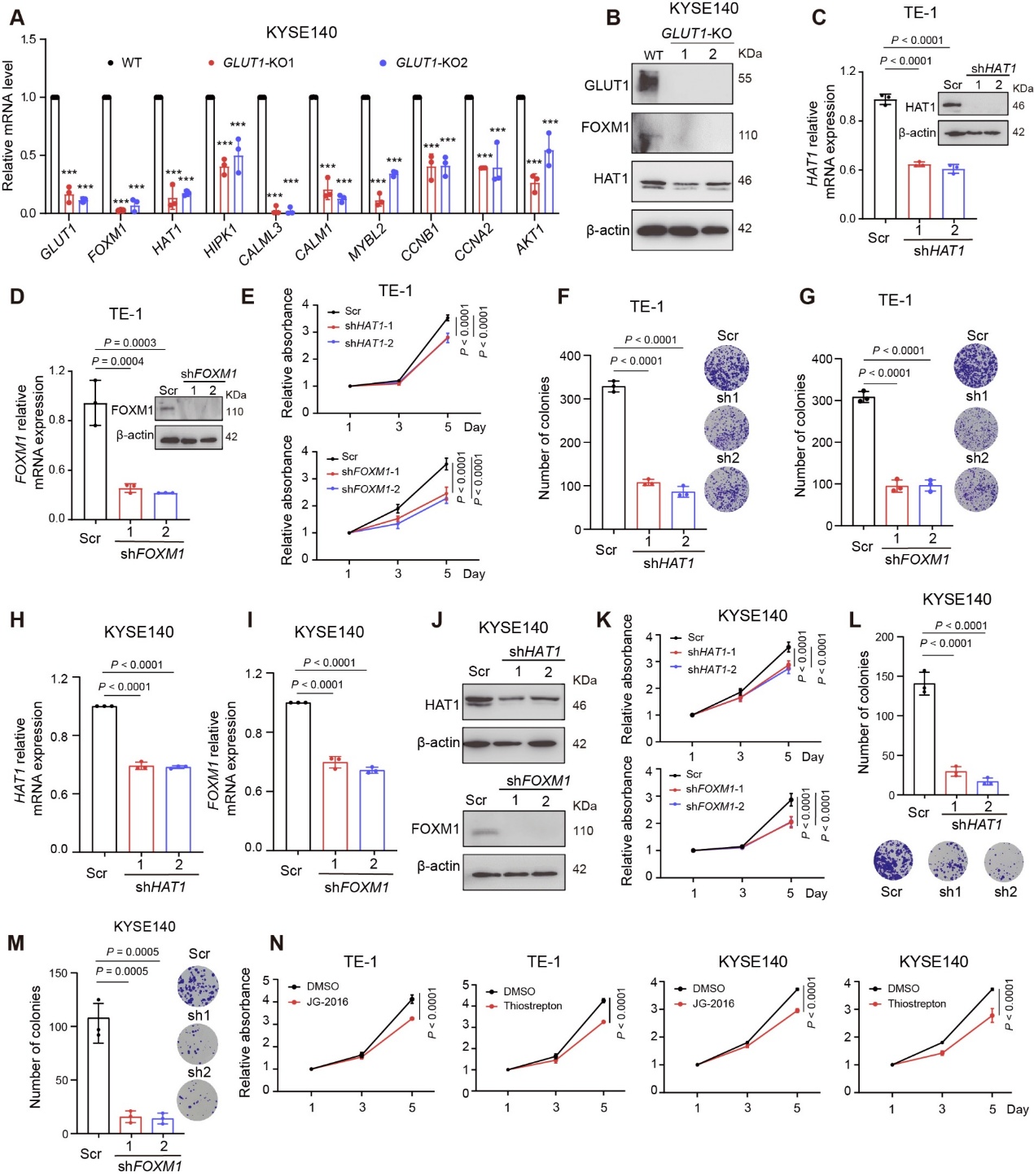

**Figure S6. GLUT1-HAT1/FOXM1 signaling promotes proliferation and colony formation in ESCC cells, related to Figure 4.**

(A) Relative mRNA levels of the indicated genes in *GLUT1*-KO compared to WT KYSE140 cells, as measured by RT-qPCR.

(B) Western blot analysis of GLUT1, FOXM1, and HAT1 protein levels in WT and *GLUT1*-KO KYSE140 cells.

(C) RT-qPCR and western blot analysis of HAT1 expression in TE-1 cells transduced with control (Scr) or two independent sh*HAT1* constructs.

(D) RT-qPCR and western blot analysis of FOXM1 knockdown in TE-1 cells using two independent sh*FOXM1* constructs.

(E) Cell proliferation of TE-1 cells expressing sh*HAT1* (top) or sh*FOXM1* (bottom) measured by MTT assay at the indicated time points.

(F and G) Colony formation assay showing the inhibitory effects of *HAT1* (F) and *FOXM1* (G) knockdown on clonogenic growth in TE-1 cells.

(H and I) RT-qPCR analysis of *HAT1* and *FOXM1* expression levels in KYSE140 cells after *HAT1* and *FOXM1* knockdown.

(J) Western blot validation of HAT1 and FOXM1 knockdown in KYSE140 cells.

(K) MTT proliferation assays showing that *HAT1* or *FOXM1* silencing suppresses KYSE140 cell growth.

(L and M) Colony formation assays demonstrating that *HAT1* and *FOXM1* knockdown markedly reduces clonogenic capacity of KYSE140 cells.

(N) Proliferation assays indicating that pharmacological inhibition of FOXM1 using JG-2016 or thiostrepton suppresses cell growth in TE-1 and KYSE140 cells compared to the DMSO control.

Data are presented as mean ± SEM from three independent experiments. Statistical significance was determined using Student’s t-test. ****P* < 0.001; *****P* < 0.0001. Data are presented as mean ± SEM of three independent experiments, and statistical comparisons were performed using one-way ANOVA with Tukey’s multiple comparison test (A, C-I, K, N).

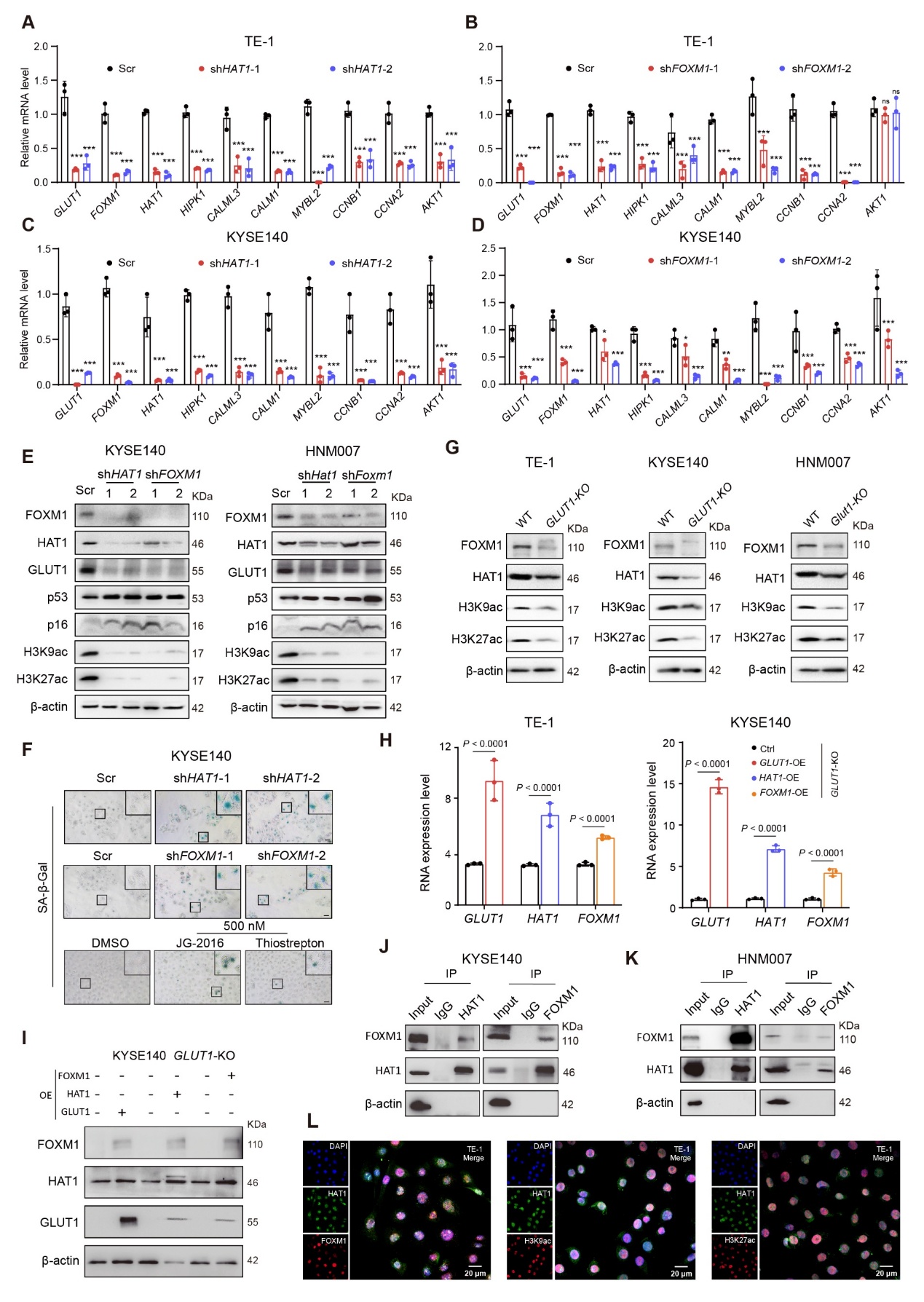

**Figure S7. The HAT1/FOXM1 axis regulates cellular senescence, histone acetylation, and downstream transcriptional programs in ESCC cells, related to Figure 4.**

(A and B) RT-qPCR analysis of the indicated genes in TE-1 cells transduced with sh*HAT1* (A) or sh*FOXM1* (B). Relative mRNA levels were normalized to Scr control.

(C and D) RT-qPCR analysis of the indicated genes in KYSE140 cells following *HAT1* (C) or *FOXM1* (D) knockdown.

(E) Western blot analysis in KYSE140 and HNM007 cells showing the effects of HAT1 or FOXM1 silencing on the protein expression of FOXM1, HAT1, GLUT1, and senescence markers p53, p16, as well as histone acetylation marks (H3K9ac and H3K27ac).

(F) Representative images of SA-β-Gal staining in KYSE140 cells following *HAT1* or *FOXM1* knockdown, or pharmacological inhibition using JG-2016 and thiostrepton , respectively.

(G) Western blot analysis of FOXM1, HAT1, H3K9ac, and H3K27ac in WT and *GLUT1*-KO cells across TE-1, KYSE140, and HNM007 cell lines.

(H) RT-qPCR showing of the mRNA levels of *GLUT1,* *HAT1*, and *FOXM1* upon overexpression (OE) of indicated genes in *GLUT1*-KO TE-1 cells (left) and KYSE140 cells (right).

(I) Western blot validation of GLUT1, HAT1, and FOXM1 protein levels in *GLUT1*-KO KYSE140 cells following the indicated rescue experiments.

(J and K) Co-immunoprecipitation (Co-IP) assays in KYSE140 (J) and HNM007 (K) cells demonstrating the interaction between HAT1 and FOXM1. IgG served as a negative control.

(L) Representative IF images in TE-1 cells showing the subcellular localization and co-expression of HAT1 (green) and FOXM1 (red), as well as association with histone acetylation marks (H3K9ac and H3K27ac). Nuclei were stained with DAPI. Scale bar, 20 μm.

Data are presented as mean ± SEM from three independent experiments. Statistical significance was determined using Student’s t-test. **P* < 0.05 ***P* < 0.01; ****P* < 0.001; ns, not significant. Data are presented as mean ± SEM of three independent experiments, and statistical comparisons were performed using student’s test (K, L, N, and O) and one-way ANOVA with Tukey’s multiple comparison test (C-E, G-J, M, Q, S-W).

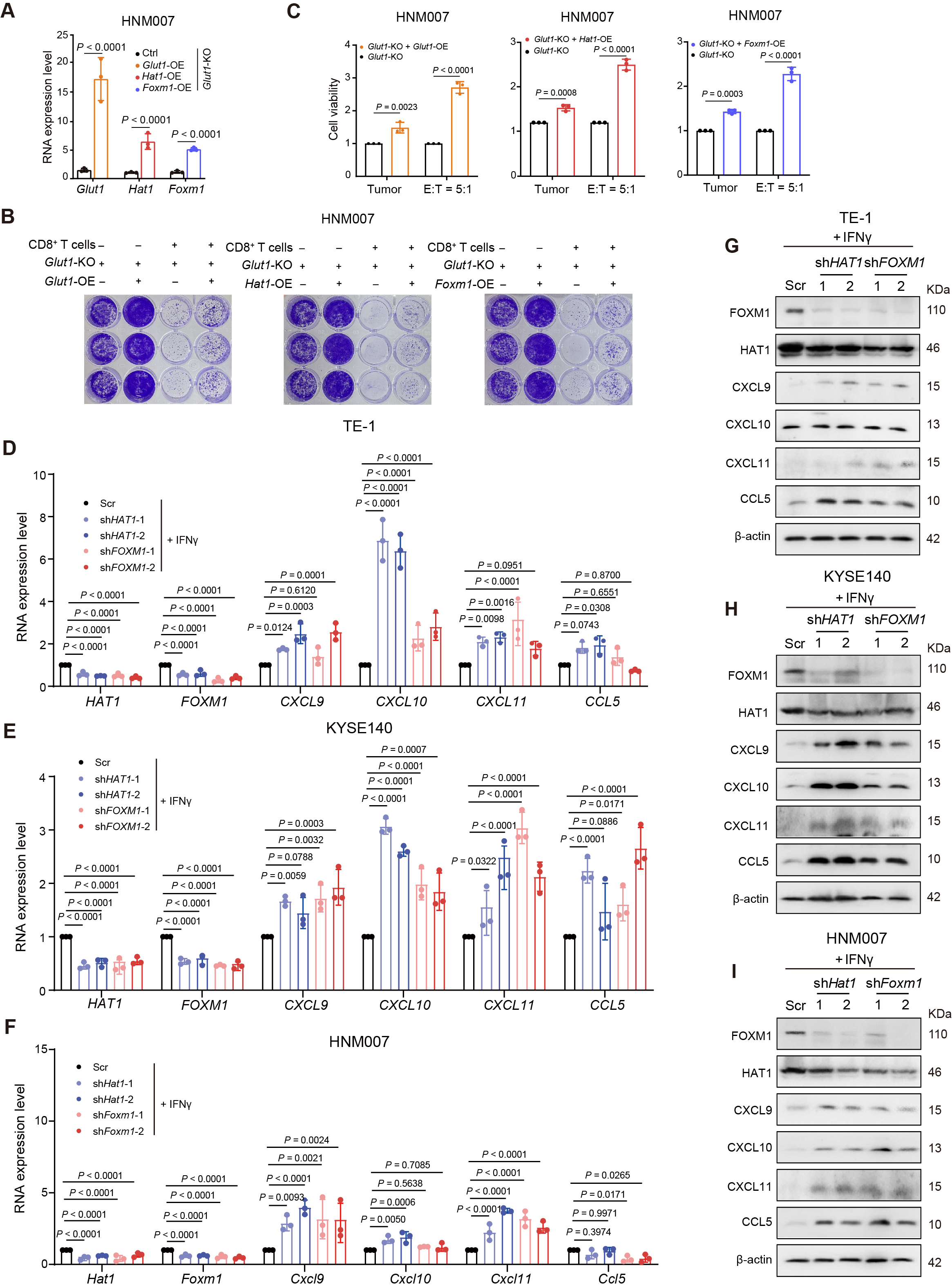

**Figure S8. The GLUT1-HAT1/FOXM1 axis regulates CD8⁺ T cell mediated cytotoxicity and SASP chemokine expression in ESCC cells, related to Figure 5.**

(A) RT-qPCR analysis showing the expression levels of *Glut1*, *Hat1*, and *Foxm1* in the *Glut1*-KO HNM007 cells with exogenous overexpression (OE) of the indicated genes.

(B) Crystal violet staining showing the viability of HNM007 cells with *Glut1*-KO or exogenous overexpression of Glut1, Hat1 or Foxm1 in *Glut1*-KO cells, upon co-culture with CD8⁺ T cells.

(C) Quantification of tumor cell viability in rescue experiments of (B).

(D-F) RT-qPCR analysis of *HAT1*, *FOXM1*, and SASP chemokines in TE-1, KYSE140 and HNM007 cells following *HAT1* or *FOXM1* knockdown.

(G-I) Western blot analysis showing the protein levels of SASP chemokines including CXCL9, CXCL10, CXCL11, and CCL5 in TE-1 (G), KYSE140 (H), and HNM007 (I) cells.

Data are presented as mean ± SEM of three independent experiments, and statistical comparisons were performed using one-way ANOVA with Tukey’s multiple comparison test (A, C, D-F).

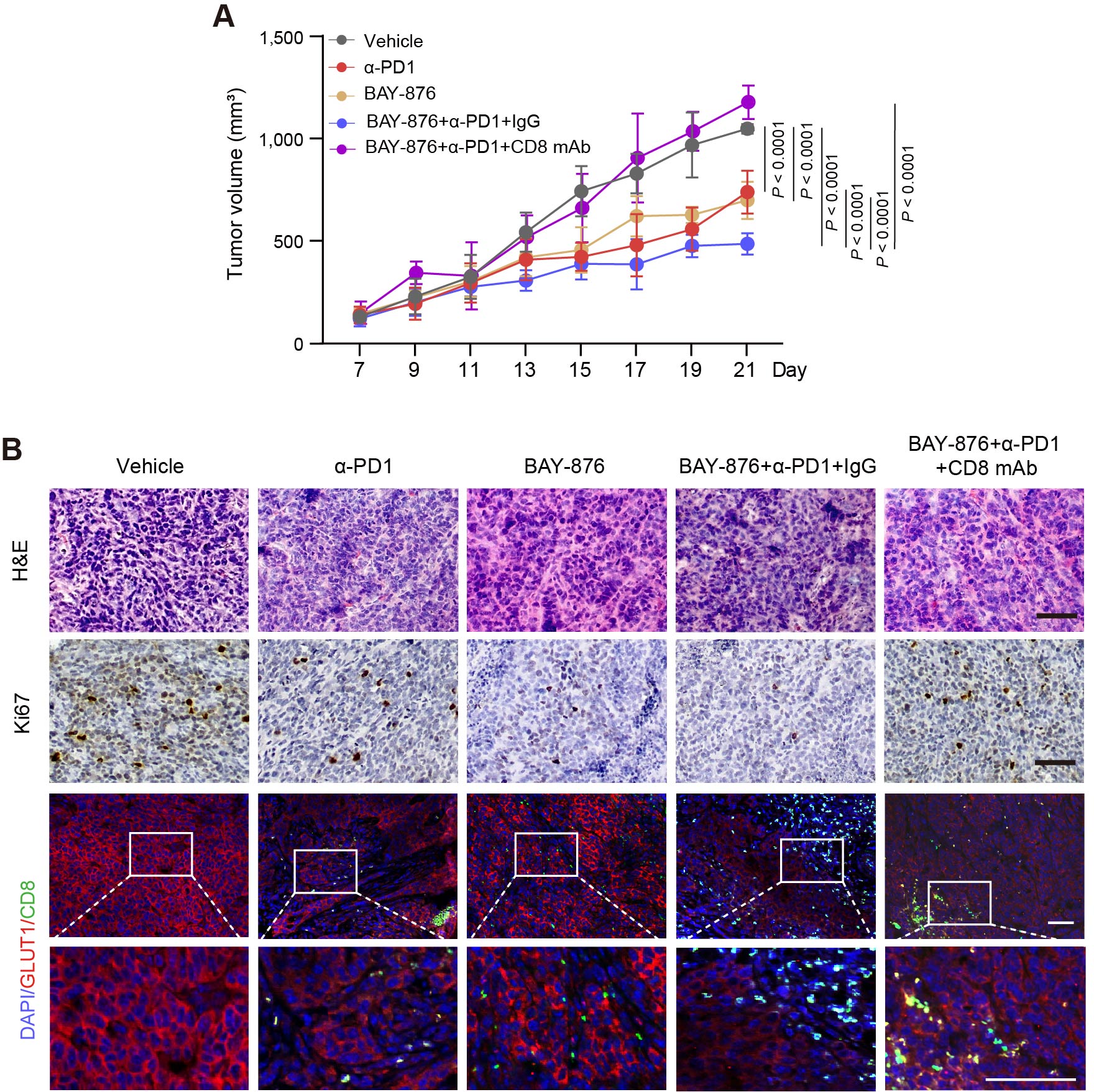

**Figure S9. BAY-876 enhances the therapeutic efficacy of α-PD1 in HNM007 syngeneic tumors, related Figure 6.**

(A) Statistical analysis of tumor growth for the indicated treatment groups in Figure 6B. N = 6.

(B) Representative H&E staining and IF analyses of tumor tissue sections from each treatment group. H&E staining shows tumor morphology; Ki-67 staining indicates proliferative activity; and IF staining of GLUT1 (red), CD8⁺ T cells (green), and DAPI (blue) shows the spatial distribution of CD8⁺ T cells and GLUT1 expression. Insets indicate magnified regions. Scale bar, 100 μm. Data are presented as mean ± SEM. using one-way ANOVA with Tukey’s multiple comparison test.

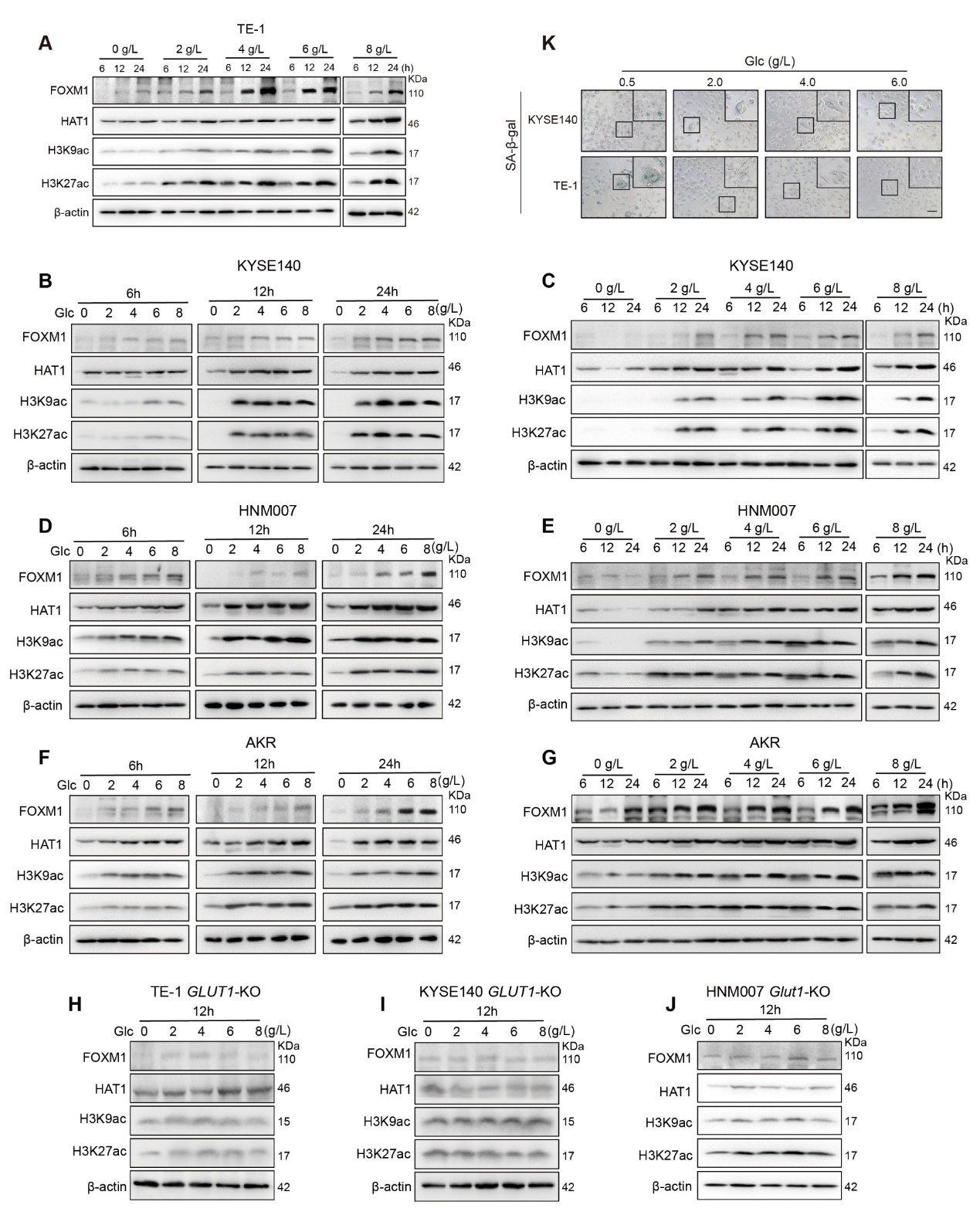

**Figure S10. Glucose availability modulation regulates the HAT1/FOXM1 axis and histone acetylation in a GLUT1-dependent manner, related to Figure 6.**

(A) Western blot analysis showing the protein levels of FOXM1, HAT1, H3K9ac, and H3K27ac at indicated time points (6, 12, 24 h) and glucose concentrations in KYSE140 cells.

(B-G) Western blot analysis showing the protein levels of FOXM1, HAT1, H3K9ac, and H3K27ac under different glucose concentrations (0-8 g/L) over time (6, 12, and 24 h) (A) or at indicated time points (6, 12, 24 h) and glucose concentrations in KYSE140 (B, C), HNM007 (D, E), and AKR (F, G) cells.

(H-J) Western blot analysis showing protein levels of FOXM1, HAT1, H3K9ac, and H3K27ac at indicated time points and glucose concentrations in *GLUT1*-KO TE-1, KYSE140, and HNM007 cells.

(K) Representative SA-β-Gal staining images showing the glucose-dependent regulation of cellular senescence.

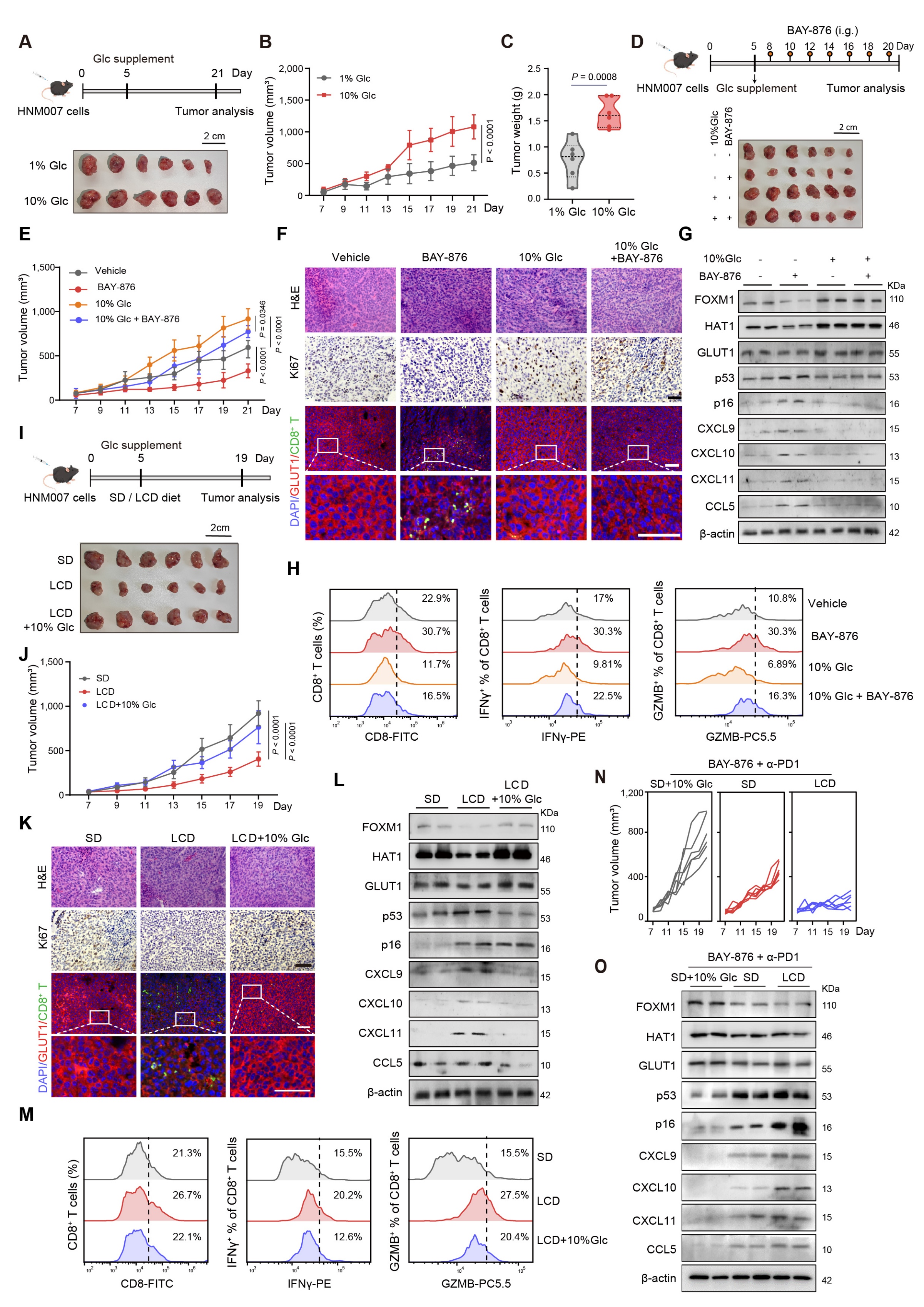

**Figure S11. High glucose promotes tumor growth and immune evasion via the GLUT1-HAT1/FOXM1 axis and modulates the response to metabolic and immunotherapy interventions, related to Figure 7.**

(A) Experimental schematic of tumors derived from HNM007 cells (1 × 10⁵ cells) implanted into mice under different glucose conditions (1% *vs* 10% glucose). Photographs of isolated tumors on day. N = 6.

(B) Tumor growth curves showing increased tumor volume in the 10% glucose group compared to the 1% glucose group.

(C) Tumor weight comparison between low- and high-glucose groups.

(D) Experimental design and tumor images from mice bearing HNM007 tumors (1 × 10⁵ cells) treated with BAY-876 (i.g.) under high-glucose conditions. N = 6.

(E) Tumor growth curves showing that BAY-876 suppresses tumor progression.

(F) Histological and immunofluorescence analyses (H&E, Ki-67, and multiplex IF for GLUT1/CD8⁺ T cells/DAPI) of tumor tissues from the indicated groups. Scale bars, 100 μm.

(G) Western blot analysis of tumor lysates showing the expression of FOXM1, HAT1, GLUT1, and senescence markers p53, p16, and chemokines (CXCL9, CXCL10, CXCL11, CCL5) under different treatment conditions.

(H) Flow cytometry analysis of tumor-infiltrating CD8⁺ T cells and their functional effector markers (IFNγ and GZMB) across treatment groups.

(I) Experimental schematic and tumor images of mice bearing HNM007 tumors (5 × 10⁵ cells) under different dietary conditions (SD vs LCD ± 10% glucose supplementation). N = 6.

(J) Tumor growth curves showing that LCD diet suppresses tumor growth, while glucose supplementation partially restores tumor progression.

(K) Histological and multiplex immunofluorescence analyses of tumor tissues from SD, LCD, and LCD + 10% glucose groups. Scale bars, 100 μm.

(L) Western blot analysis of tumor lysates showing the regulation of the Glut1, Hat1, Foxm1, p53/p16, and chemokine expression under the specified dietary conditions.

(M) Flow cytometry analysis of CD8⁺ T cell infiltration and effector function (IFNγ and GZMB) in tumors from different dietary groups.

(N) Tumor growth curves showing the therapeutic effects of BAY-876 combined with α-PD1 under different metabolic conditions (10% glucose, SD, and LCD).

(O) Western blot analysis of tumor tissues showing molecular changes following combination therapy (BAY-876 + α-PD1), including effects on FOXM1, HAT1, GLUT1, p53, p16, and chemokines.

Data are presented as mean ± SEM, and statistical comparisons were performed using student’s test (C), and one-way ANOVA with Tukey’s multiple comparison test (B, E, J). *P* values are indicated in the figures.

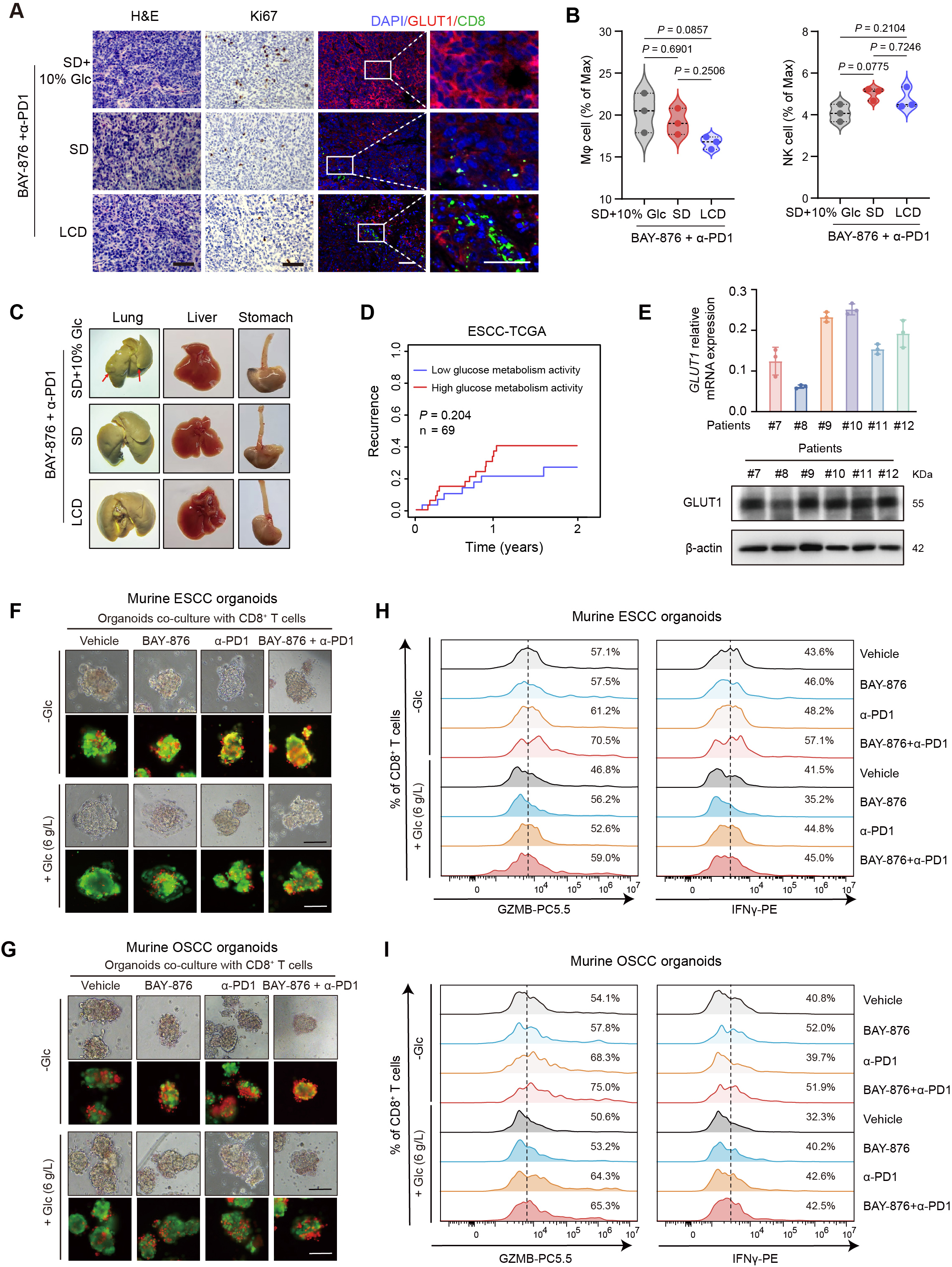

**Figure S12. Dietary glucose restriction enhances the efficacy of BAY-876 and α-PD1 therapy by promoting anti-tumor immunity, related to Figure 7.**

(A) Representative images of lung, liver, and stomach tissues from treated mice, showing reduced metastatic burden under LCD compared to the high-glucose conditions following BAY-876 + α-PD1 therapy.

(B) Quantification of tumor-infiltrating macrophages (MΦ) and NK cells in mice treated with BAY-876 + α-PD1 under different metabolic conditions (10% glucose, SD, and LCD). Statistical comparisons were performed using one-way ANOVA with Tukey’s multiple comparison test.

(C) Images of lung, liver, and stomach tissues from tumor-bearing mice treated with BAY-876 + α-PD1 under different dietary conditions: control diet with 10% glucose (SD + 10% Glc), SD, and LCD. Arrows indicate visible metastatic nodules in the lungs.

(D) Cumulative incidence function (CIF) analysis comparing recurrence risk between ESCC patients with high versus low glucose metabolism activity in the ESCC-TCGA cohort. Patients were stratified into high-activity (n = 34) and low-activity (n = 35) groups according to the median ssGSEA score of the glycolysis-related gene signature. CIF analysis showed a trend toward increased recurrence risk in patients with high glucose metabolism activity.

(E) Relative expression of GLUT1 in patient-derived samples (left) and corresponding western blot analysis of GLUT1 protein levels (right). β-actin serves as a loading control.

(F and G) Representative bright-field and live/dead fluorescence images of murine ESCC-derived (A) and OSCC-derived (C) organoids co-cultured with CD8⁺ T cells under glucose-deprived (-Glc) or glucose-supplemented (+Glc, 6 g/L) conditions and treated with vehicle, BAY-876, anti-PD-1 (α-PD1), or their combination. Green indicates live cells and red indicates dead cells. Scale bars, as indicated.

(H and I) Flow cytometry analysis of cytotoxic CD8⁺ T cells from co-culture systems shown in (A) and (C), quantified by GZMB and IFNγ expression under -Glc or +Glc conditions. Percentages indicate the proportion of GZMB⁺ or IFNγ⁺ CD8⁺ T cells.

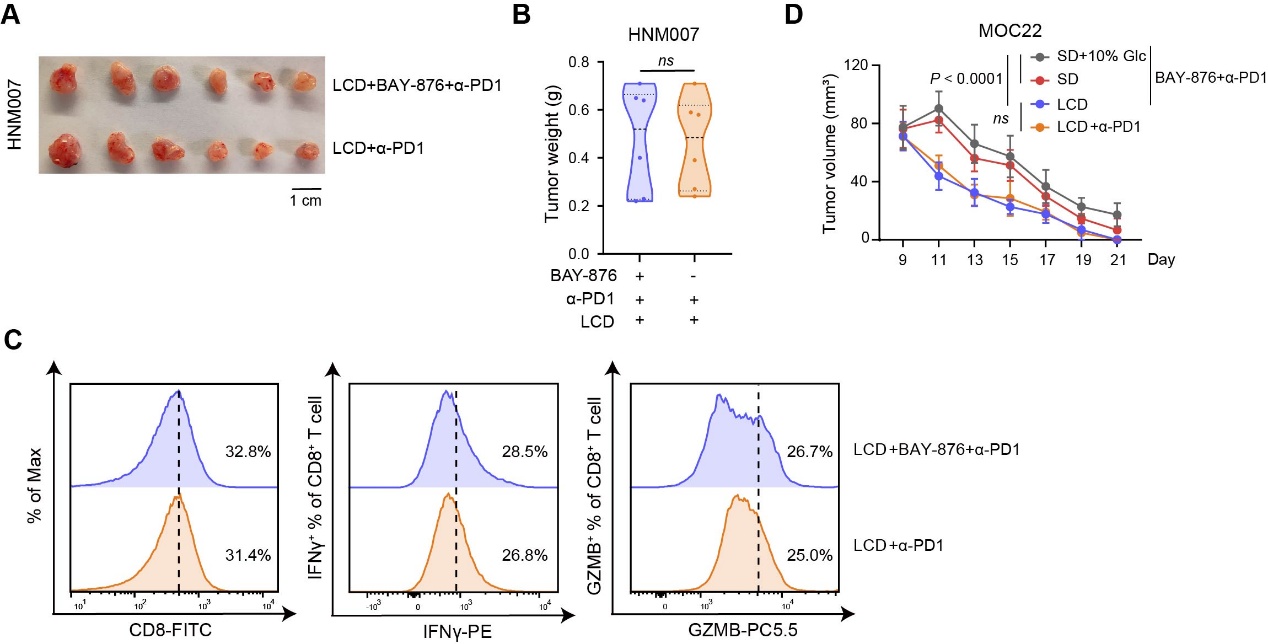

**Figure S13. Glucose availability modulates CD8⁺ T cell mediated cytotoxicity and the efficacy of GLUT1 inhibition in tumor organoid models.**

(A and B) Representative tumor images (A) and tumor weight quantification (B) showing enhanced tumor suppression in HNM007-bearing mice receiving the triple combination therapy (LCD + BAY-876 + α-PD1).

(C) Flow cytometry analysis of tumor-infiltrating CD8⁺ T cells, showing increased frequencies and enhanced effector function (IFNγ and GZMB) under combination treatment.

(D) Tumor growth curves of MOC22 tumor-bearing mice under different dietary conditions (10% Glc, SD, and LCD) in combination with BAY-876 + α-PD1 and LCD + α-PD1 therapy. Statistical significance is indicated by the corresponding *P* values, demonstrating that dietary glucose restriction enhances the antitumor efficacy of combined metabolic and immune checkpoint therapy. Statistical comparisons were performed using one-way ANOVA with Tukey’s multiple comparison test.
