## Supplementary Table for "High glucose confers senescence resistance via GLUT1 epigenetic rewiring to blunt immunotherapy responses in esophageal squamous cell carcinoma"

**Supplementary Table 1. All oligonucleotide sequences used in this study**

| **Short hairpin RNA or sgRNA target sequences** | |
| --- | --- |
| sh*HAT1*-1F(H) | TTTTGGTGCTATGGAGAAATTTTTCAAGAGAAAATTTCTCCATAGCACCAAATTTTTTC |
| sh*HAT1*-1R(H) | TCGAGAAAAAATTTGGTGCTATGGAGAAATTTTCTCTTGAAAAATTTCTCCATAGCACCAAAA |
| sh*HAT1*-2F(H) | TGCAAGGATTCAATGAAGATATTTCAAGAGAATATCTTCATTGAATCCTTGCTTTTTTC |
| sh*HAT1*-2R(H) | TCGAGAAAAAAGCAAGGATTCAATGAAGATATTCTCTTGAAATATCTTCATTGAATCCTTGCA |
| sh*FOXM1*-1F(H) | TGCAGTAGTGGGCCCAACAAATTTCAAGAGAATTTGTTGGGCCCACTACTGCTTTTTTC |
| sh*FOXM1*-1R(H) | TCGAGAAAAAAGCAGTAGTGGGCCCAACAAATTCTCTTGAAATTTGTTGGGCCCACTACTGCA |
| sh*FOXM1*-2F(H) | TCCAGCTGGGATCAAGATTATTTTCAAGAGAAATAATCTTGATCCCAGCTGGTTTTTTC |
| sh*FOXM1*-2R(H) | TCGAGAAAAAACCAGCTGGGATCAAGATTATTTCTCTTGAAAATAATCTTGATCCCAGCTGGA |
| *GLUT1*-sgF(H) | caccgGGATGCTCTCCCCATAGCGG |
| *GLUT1*-sgR(H) | aaacCCGCTATGGGGAGAGCATCCc |
| sh*Glut1*-1F(M) | TGAGGAGTTCTACAATCAAACATTCAAGAGATGTTTGATTGTAGAACTCCTCTTTTTTC |
| sh*Glut1*-1R(M) | TCGAGAAAAAAGAGGAGTTCTACAATCAAACATCTCTTGAATGTTTGATTGTAGAACTCCTCA |
| sh*Glut1-*2F(M) | TGCTCAGATCTATTCAGATAAGTTCAAGAGACTTATCTGAATAGATCTGAGCTTTTTTC |
| sh*Glut1-*2R(M) | TCGAGAAAAAAGCTCAGATCTATTCAGATAAGTCTCTTGAACTTATCTGAATAGATCTGAGCA |
| Dox-sh*Glut1*-F(M) | GTTGACAGTGAGCGGGCTCAGATCTATTCAGATAAGTAGTGAAGCCACAG |
| Dox-sh*Glut1*-R(M) | CGAGGCAGTAGGCATGCTCAGATCTATTCAGATAAGTACATCTGTGGCTTC |
| sh*Hat1*-1F(M) | TCGTGTTGAATATTCATCTAAATTCAAGAGATTTAGATGAATATTCAACACGTTTTTTC |
| sh*Hat1*-1R(M) | TCGAGAAAAAACGTGTTGAATATTCATCTAAATCTCTTGAATTTAGATGAATATTCAACACGA |
| sh*Hat1*-2F(M) | TGCTACATGACAGTCTATAATTTTCAAGAGAAAttatagactgtcatgtagcTTTTTTC |
| sh*Hat1*-2R(M) | TCGAGAAAAAAgctacatgacagtctataaTTTCTCTTGAAAATTATAGACTGTCATGTAGCA |
| sh*Foxm1*-1F(M) | TGGCAAAGACAGGAGAGCTATGTTCAAGAGACATAGCTCTCCTGTCTTTGCCTTTTTTC |
| sh*Foxm1*-1R(M) | TCGAGAAAAAAGGCAAAGACAGGAGAGCTATGTCTCTTGAACATAGCTCTCCTGTCTTTGCCA |
| sh*Foxm1*-2F(M) | TCCGGAATCAAGATTATCAACCTTCAAGAGAGGTTGATAATCTTGATTCCGGTTTTTTC |
| sh*Foxm1*-2R(M) | TCGAGAAAAAACCGGAATCAAGATTATCAACCTCTCTTGAAGGTTGATAATCTTGATTCCGGA |
| **Primer oligos used for qPCR** | |
| *GLUT1*-F(H) | GGCCAAGAGTGTGCTAAAGAA |
| *GLUT1*-R(H) | ACAGCGTTGATGCCAGACAG |
| *FOXM1*-F(H) | CAGGTGTTTAAGCAGCAGCA |
| *FOXM1*-R(H) | CTGAGCTCATGAGGGAAGCC |
| *AKT1*-F(H) | CGAGCTGTTCTTCCACCTGT |
| *AKT1*-R(H) | CGACCGCACATCATCTCGTA |
| *CALML3*-F(H) | CTGCGGGACATGATGAGTGA |
| *CALML3*-R(H) | CACACGGACAAACTCCTCGT |
| *CALM1*-F(H) | CACTGGGTCAGAACCCAACA |
| *CALM1*-R(H) | ACGTGACGTAGTTCTGCTGC |
| *MYBL2*-F(H) | GTTTGGACAGCAGGACTGGA |
| *MYBL2*-R(H) | TGGCAACATCTTGGCGATCT |
| *CDKN2A*-F(H) | ATGGAGCCTTCGGCTGACT |
| *CDKN2A*-R(H) | CGAAGCGCTACCTGATTCCA |
| *CDKN1A*-F(H) | ACTTCGACTTTGTCACCGAG |
| *CDKN1A*-R(H) | CGTTTGGAGTGGTAGAAATCTGTC |
| *HIPK1*-F(H) | TCCAGCGGAGAAGGGGATTA |
| *HIPK1*-R(H) | TCACTGCTTAGGCGGGAAAG |
| *CCNA2*-F(H) | CGGTACTGAAGTCCGGGAAC |
| *CCNA2*-R(H) | CAGGGCATCTTCACGCTCTA |
| *CCNB1*-F(H) | GCACTTCCTTCGGAGAGCAT |
| *CCNB1*-R(H) | CCAGGTGCTGCATAACTGGA |
| *HAT1*-F(H) | AGGCAGATGATGTTGAGGGC |
| *HAT1*-R(H) | TACGGTCGCAAAGAGCGTAG |
| *CXCL9*-F(H) | ATTGGTGCCCAGTTAGCCTC |
| *CXCL9*-R(H) | CCACCGGACAGCACTCTAAA |
| *CXCL10*-F(H) | ACCAGAGGGGAGCAAAATCG |
| *CXCL10*-R(H) | GCAGGGTCAGAACATCCACT |
| *CXCL11*-F(H) | TGTCTTTGCATAGGCCCTGG |
| *CXCL11*-R(H) | AGCCTTGCTTGCTTCGATTTG |
| *CCL5*-F(H) | TCATTGCTACTGCCCTCTGC |
| *CCL5*-R(H) | TCGGGTGACAAAGACGACTG |
| *ACTIN*-F(H) | AGCGAGCATCCCCCAAAGTT |
| *ACTIN*-R(H) | GGGCACGAAGGCTCATCATT |
| *Glut1*-F(M) | GCTTCTCCAACTGGACCTCAAAC |
| *Glut1*-R(M) | ACGAGGAGCACCGTGAAGATGA |
| *Hat1*-F(M) | TCTTCGACTGCTGGTGACTG |
| *Hat1*-R(M) | GGGAACAAGGGCTTCACTCT |
| *Foxm1*-F(M) | TTTCAGCCCAGTACGAACCC |
| *Foxm1*-R(M) | GGCTGAAAGGCTGGGAATCT |
| *Akt1*-F(M) | CAGGACCACGAGAAGCTGTT |
| *Akt1*-R(M) | TCCACACACTCCATGCTGTC |
| *Calml3*-F(M) | TGACTGAGGAGCAAATCGCC |
| *Calml3*-R(M) | ATCTCCTCCTCGCTGTCAGT |
| *Calm1*-F(M) | TCAGAACCCAACAGAAGCCG |
| *Calm1*-R(M) | GTTGACTTGTCCGTCGCCAT |
| *Mybl2*-F(M) | GTCGCCTGTCACAGAGAACA |
| *Mybl2*-R(M) | GGGAGTACTTCTGATGATGGATAC |
| *Cdkn2a*-F(M) | TCACACGACTGGGCGATTG |
| *Cdkn2a*-R(M) | CGTGAACGTTGCCCATCATC |
| *Cdkn1a*-F(M) | GCAAAGTGTGCCGTTGTCTC |
| *Cdkn1a*-R(M) | TTTCGGCCCTGAGATGTTCC |
| *Hipk1*-F(M) | AGTGCACAATCAGGCCAGTG |
| *Hipk1*-R(M) | AGGGTGGAGAGTTGCTACCAT |
| *Ccna2*-F(M) | TGCAGCTGTCTCTTTACCCG |
| *Ccna2*-R(M) | TGGCCCTCATGCTGTTAGTG |
| *Ccnb1*-F(M) | CAGGCAAGAGTGCCTCTGAA |
| *Ccnb1*-R(M) | TTTGGGTCAGCCCCATCATC |
| *Cxcl9*-F(M) | GCAGTGTGGAGTTCGAGGAA |
| *Cxcl9*-R(M) | AGTCCGGATCTAGGCAGGTT |
| *Cxcl10*-F(M) | GAGAGACATCCCGAGCCAAC |
| *Cxcl10*-R(M) | ATTCTCACTGGCCCGTCATC |
| *Cxcl11*-F(M) | CCTGACCCTCTGCTGTCTTG |
| *Cxcl11*-R(M) | AACCACAGAAGGTAGCGTGG |
| *Ccl5*-F(M) | CCTCACCATATGGCTCGGAC |
| *Ccl5*-R(M) | TCTTCTCTGGGTTGGCACAC |
| *Actin*-F(M) | GTGACGTTGACATCCGTAAAGA |
| *Actin*-R(M) | GCCGGACTCATCGTACTCC |

Note: M and H indicate mouse and human primer sequences, respectively.

**Supplementary Table 2. Key resources used in this study**

| Reagent or Resource | Source | Identifier |
| --- | --- | --- |
| **Antibodys** |  |  |
| GLUT1(SLC2A1) Polyclonal antibody | Proteintech | 21829-1-AP |
| FoxM1(D3F2B) Rabbit mAb | Cell Signaling Technology | 20459T |
| HAT1(E7Y80) Rabbit mAb | Cell Signaling Technology | 41490T |
| FOXM1 Polyclonal antibody | Proteintech | 13147-1-AP |
| KAT1 /HAT1 Recombinant Rabbit Monoclonal Antibody | HUABIO | HA722510 |
| CCL5/RANTES Rabbit pAb | A14192 | ABclonal |
| CXCL10/IP-10 Rabbit mAb | A21986 | ABclonal |
| CXCL11 Rabbit pAb | A6201 | ABclonal |
| CXCL9 Rabbit pAb | A1864 | ABclonal |
| Histone H3 (acetyl K9) Recombinant Rabbit Monoclonal Antibody | HUABIO | HA722132 |
| Anti-Histone H3 (acetyl K27) antibody | Abcam | ab177178 |
| Fixable Viability Stain 450 | BD Biosciences | 562247 |
| Donkey anti-Mouse IgG (H+L) Highly Cross-Adsorbed Secondary Antibody, Alexa FluorTM 488 | Invitrogen | A-21202 |
| Donkey anti-Mouse IgG (H+L) Highly Cross-Adsorbed Secondary Antibody, Alexa FluorTM 594 | Invitrogen | A-21203 |
| Donkey anti-Rabbit IgG (H+L) Highly Cross-Adsorbed Secondary Antibody, Alexa FluorTM 488 | Invitrogen | A-21206 |
| Donkey anti-Rabbit IgG (H+L) Highly Cross-Adsorbed Secondary Antibody, Alexa FluorTM 594 | Invitrogen | A-21207 |
| InVivoMab anti-mouse PD-1 | BioXcell | BE0146 |
| InVivoMAb rat IgG isotype control | BioXcell | BE0089; |
| InVivoMab anti-mouse CD8α | BioXcell | BE0061 |
| FITC anti-mouse CD8a Antibody; clone H1.2F3; clone | BioLegend | 100706 |
| PE anti-mouse INFγ Antibody | BioLegend | 505807 |
| PerCP/Cyanine5.5 anti-human Granzyme B Recombinant | BioLegend | 372212 |
| Mouse Anti-CD8α | Cell Signaling Technology | 70306 |
| GAPDH Monoclonal antibody | Proteintech | 60004-1-ig |
| HRP-conjugated IgG Fraction Monoclonal Mouse Anti-Rabbit IgG, Light Chain Specific | Proteintech | SA00001-7L |
| Rabbit secondary antibody | Abcepta | ASP1615 |
| Mouse secondary antibody | Abcepta | ASP1613 |
| **Bacterial and virus strains** |  |  |
| Stbl3 Chemically Competent Cell | Tsingke  Biotechnology | TSC-C06 |
| **Chemicals** |  |  |
| PolyJet DNA In Vitro Tranfection Reagent | SignaGen | SL100688 |
| BAY876 | Medchem Express | HY-100017 |
| Thiostrepton | Medchem Express | HY-B0990 |
| JG-2016 | Medchem Express | HY-154944 |
| Protein A/G PLUS-Agarose: sc-2003 | Santa Cruz | sc-2003 |
| N-Acetylcysteine (NAC), 0.5M | sigma | A9165-5G |
| Y-27632 2HCl | Selleck | S1049 |
| Collagenase, Type IV, powder | Thermo Fisher Scientiﬁc | 17104019 |
| Corning® Matrigel® Basement Membrane Matrix, LDEV-Free | Corning | 354234 |
| B-27 supplement, 50x | Thermo Fisher Scientiﬁc | 17504044 |
| N-2 Supplement, 100x | Thermo Fisher Scientiﬁc | 17502048 |
| **Critical commercial assays** |  |  |
| MTT | Macklin | 298-93-1 |
| Steady Pure Universal RNA Extraction Kit | Accurate Biology | AG21017 |
| Evo M-MLV RT Premix | Accurate Biology | AG11707 |
| SYBR® Green Premix Pro Taq HS qPCR Kit | Accurate Biology | AG11701 |
| Pro Taq HS PCR | Accurate Biology | AG11307 |
| Viability/Cytotoxicity Assay Kit for Animal Live & Dead Cells (Calcein AM, EthD-1 Method) | Amersco/UElandy | L6023M |
| Acetyl-CoA (A-CoA) Enzyme-Linked Immunosorbent Assay Kit | Elabscience | E-EL-0125 |
| **Recombinant DNA** |  |  |
| PLKO.1 | Addgene | Cat# 1864 |
| psPAX2 | Addgene | Cat# 12260 |
| PMD2.G | Addgene | Cat# 12259 |
| pLV3-CMV-SLC2A1(human)-3×FLAG-CopGFP-Puro | MIAOLING PLASMID | P54238 |
| pLV3-CMV-Slc2a1(mouse)-3×FLAG-CopGFP-Puro | MIAOLING PLASMID | P54201 |
| pLV3-CMV-HAT1(human)-3xFLAG-CopGFP-Puro | MIAOLING PLASMID | P67937 |
| pLV3-CMV-FOXM1(human)-3xFLAG-Puro | MIAOLING PLASMID | p60179 |
| pLV3-CMV-Foxm1(mouse)-3xFLAG-CopGFP-Puro | MIAOLING PLASMID | P56560 |
| pLV3-CMV-Hat1(mouse)-3xFLAG-CopGFP-Puro | MIAOLING PLASMID | p44394 |
